# Dual role of ZIC2 during neural induction: from pioneer transcription factor to enhancer activator

**DOI:** 10.1101/2025.07.23.666365

**Authors:** María Mariner-Faulí, Víctor Sánchez-Gaya, Sarah Malika Robert, Patricia Respuela-Alonso, Sara de la Cruz-Molina, Sara Lobato-Moreno, Matteo Trovato, Karin D. Prummel, Kyung-Min Noh, Judith B. Zaugg, Álvaro Rada-Iglesias

## Abstract

ZIC2, a member of the Zinc Finger of the Cerebellum family of transcription factors (TFs), plays crucial roles during neural development. In humans, defects in ZIC2 cause holoprosencephaly, a congenital brain malformation characterized by the defective cleavage of cerebral hemispheres due to problems in midline patterning. However, the gene regulatory network (GRN) controlled by ZIC2 and the regulatory mechanisms it employs during neural development remain largely unexplored. Here, we combined a mouse embryonic stem cell (mESC) *in vitro* differentiation model towards anterior neural progenitors (AntNPCs) with genome editing approaches, bulk and single cell (i.e. Multiome scATAC + scRNAseq) genomic methods to elucidate the precise GRN controlled by ZIC2 and the underlying mechanisms. We found that ZIC2 shows widespread binding throughout the genome already in mESC as well as upon pluripotency exit and neural induction. Despite its extensive binding in mESC, ZIC2 function is dispensable in pluripotent cells due to compensation by ZIC3. In contrast, ZIC2 plays a major regulatory function during neural induction, directly controlling the expression of master regulators implicated in the patterning and morphogenesis of specific brain regions, such as the midbrain (e.g., *En1, Lmx1b, Pax2, Wnt1*) and the roof plate (e.g., *Lmx1a, Wnt3a*). Mechanistically, ZIC2 plays a dual role in neural differentiation: (i) during pluripotency exit, ZIC2 acts as a pioneer TF, binding *de novo* to distal enhancers and promoting their chromatin accessibility; (ii) during neural induction, ZIC2 is essential for the activation of a subset of the previously primed enhancers, which in turn control the expression of major neural patterning regulators and signaling pathways (i.e. WNT) that prevent the premature differentiation of neural progenitors. Overall, our work shows that, by sequentially acting as a promiscuous pioneer and selective activator of enhancer elements, ZIC2 canalizes pluripotent cells towards neural progenitors with rostro-dorsal identities.

## Introduction

Neural induction, like other developmental processes, is orchestrated by precise spatio-temporal transcriptional regulation. This entails the dynamic interplay between the chromatin landscape and transcription factors (TFs) that bind enhancer elements whose activation/repression ultimately shapes cell fate during developmental transitions (1–3). Among the TFs implicated in neural induction, ZIC2 -a member of the Zinc Finger of the Cerebellum family (ZIC1 to ZIC5)-plays a crucial role (4). In humans and mice, *ZIC2* loss of function (LOF) causes holoprosencephaly (HPE; MIM 236100), a forebrain congenital defect characterized by the failure of the cerebral hemispheres and midline cortical structures to properly separate (5–7). HPE can be broadly categorized into two major types, based on both the extent of forebrain separation and the location of the developmental defect with respect to the dorsoventral axis. The classical forms of HPE —comprising alobar, semilobar, and lobar subtypes— are characterized by an incomplete separation of the cerebral hemispheres, in which the lack of separation is most severe ventrally (7). In contrast, the middle interhemispheric variant (MIHV) exhibits proper separation of the ventral forebrain but displays a failure in the division of the dorsal midline, along the posterior frontal and parietal lobes (7). Notably, while multiple genes have been associated with classical HPE, *ZIC2* remains the only gene linked to both classical and MIHV forms to date, underscoring its potential role in regulating distinct spatial and temporal gene expression programs during anterior brain development.

Various *Zic2*-defective mouse models have been established (8–11), which partially recapitulate the neural tube defects observed in human HPE patients. The phenotypic alterations found in these HPE mouse models provide hints on the different processes in which ZIC2 intervenes during brain development. Firstly, in the mid-gastrula node, ZIC2 is involved in establishing the Nodal gradient, and its total LOF impairs anterior notochord and prechordal plate formation, leading to failed SHH signaling and ventral forebrain defects resembling those associated with classical HPE (12, 13). Later, ZIC2 acts within the dorsal neuroectoderm to promote the formation of the roof plate and choroid plexus, which, if disrupted, cause dorsal forebrain defects resembling those found in patients with the middle interhemispheric variant of HPE (MIHV-HPE) (10). While ZIC2 has been associated with both HPE subtypes, the precise mechanisms by which ZIC2 regulates gene expression during brain development are largely unknown. Similarly, the potential interactions between ZIC2 and the complex signaling network controlling brain patterning remain elusive. In this regard, despite the clear neural tube defects caused by ZIC2 LOF *in vivo*, most mechanistic and genomic insights into ZIC2 function have been obtained in the context of pluripotency (14–16). Previous work in mESC indicates that ZIC2 binds to many enhancers and mediates their silencing through the recruitment of co-repressors, which in turn is important for proper mESC differentiation (14). A similar and widespread binding of ZIC2 to active enhancers was also reported in Epiblast Stem Cells (EpiSC), which led to the suggestion of ZIC2 being involved in the establishment of the primed pluripotency regulatory network (15). Interestingly, ZIC2 binds the EpiSC enhancers already in mESC when they are still inactive, suggesting that ZIC2 could prime these enhancers for their subsequent activation. More recently, work in human ESC (hESC) showed that ZIC2, together with ZIC3, regulate chromatin accessibility during the transition from naive to primed pluripotency through the recruitment of BRG1 (16), thus suggesting that ZIC2 could act as a pioneer TF rather than as a repressor. In fact, the role of ZIC proteins as pioneer TFs might be evolutionary conserved, as previous studies in *Drosophila melanogaster* indicate that Opa, the *Drosophila* homolog of ZIC TFs, modulates chromatin accessibility genome-wide during the establishment of the body plan segmentation upon cellularization (17, 18). The seemingly contradicting roles reported for ZIC2 in pluripotent cells could be attributed to the difficulties in assessing its function due to the partial redundancy between ZIC2 and ZIC3 (16, 19–21). Accordingly, these two TFs are the most expressed members of the ZIC family in ESC (22), reflecting their conserved upregulation during *in vivo* peri-implantation development in mouse, primate, and human embryos (23–26). Furthermore, the partial redundancy between ZIC2 and ZIC3 in pluripotent cells could also explain why ZIC2 LOF *in vivo* does not cause defects during pre- or peri-implantation developmental stages (10, 12). However, as development progresses, the redundancy between ZIC proteins diminishes and reduced ZIC2 levels lead to brain defects both in mouse (10, 12) and humans (5, 6). Nevertheless, there are limited insights into the Gene Regulatory Network (GRN) controlled by ZIC2 during brain development as well as into the mechanisms whereby ZIC2 controls its target genes. In this regard, it is currently unknown whether, as reported in pluripotent cells, ZIC2 might preferentially act as a pioneer TF during neural induction and brain patterning or, whether, alternatively, the role of ZIC2 might be more dynamic and cell context dependent.

To resolve ZIC2 function and its underlying molecular mechanisms, we used a mESC *in vitro* differentiation protocol towards neural progenitors as a genetically tractable system to model early developmental transitions (27–30). Following a stage-specific, multi-omic strategy that combines bulk and single-cell approaches, our work dissects the gene regulatory network and chromatin landscape dynamics regulated by ZIC2 during the differentiation of mESCs into anterior neural progenitors (AntNPCs) (28, 31). By combining both constitutive ZIC2 *knock-out* and conditional ZIC2 degron cell lines, our approach leverages the complementary strengths of bulk and single-cell genomic approaches to faithfully capture ZIC2-dependent regulatory events. As a result, we identified the genes and cis-regulatory elements (CREs) that ZIC2 regulates during the *in vitro* transition of mESCs into AntNPCs. Our findings revealed that ZIC2 drives the transition from pluripotency to neuroectodermal identity by first acting as a promiscuous pioneer factor, and then as a selective enhancer activator, canalizing cell fate towards neural progenitors with anterior-dorsal identities.

## Results

### Generation of *Zic2*^*-/-*^ and *Zic2*^*FLAG-HA*^ mESCs lines

As a first step to resolve ZIC2 function during neural induction, two mESC lines were genetically engineered using CRISPR-Cas9 for subsequent differentiation. First, *Zic2*^*-/-*^ mESC lines were generated to interrogate the transcriptional and molecular defects associated with the loss of ZIC2 upon neural differentiation (Figure 1 A; Figure S1 A-C). In addition, a *Zic2*^*FLAG-HA*^ mESC line was generated, in which the endogenous ZIC2 was homozygously tagged with a FLAG-HA epitope (Figure 1 A ; Figure S1 D-G). This cell line allowed us to interrogate the ZIC2 binding profiles during neural differentiation while overcoming possible antibody specificity issues associated with the high sequence similarity of ZIC family members (4).

**Fig. 1.**
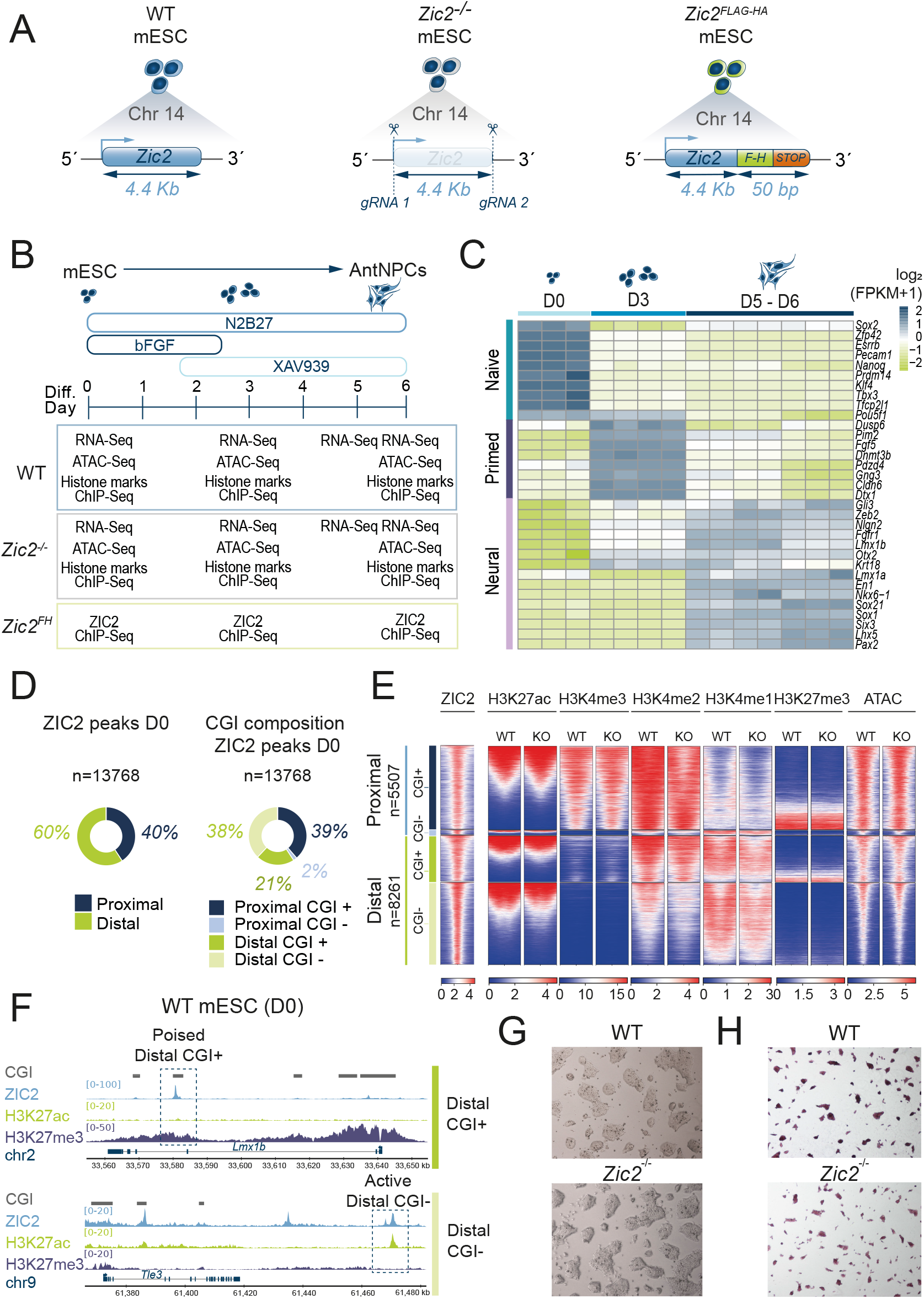
Generation and characterization of mESC lines to assess ZIC2 function in pluripotent cells and during neural induction. **(A)** Graphical overview of the mESCs used in this work to investigate ZIC2 function. From left to right: wild-type mESC (WT; E14Tg2a strain), representing our parental control cell line; *Zic2*^*-/-*^ mESC, containing a 4.4 Kb deletion spanning the whole *Zic2* gene; *Zic2*^*Flag-HA*^ mESCs, in which a Flag-HA epitope was homozygously inserted in-frame immediately upstream of the *Zic2* stop codon. **(B)** Graphical summary of the AntNPC *in vitro* differentiation protocol and the bulk genomic datasets obtained for each differentiation time-point and transgenic mESC lines described in (A). **(C)** Heatmap plot showing the temporal expression dynamics (as measured by bulk RNA-Seq) of major naive pluripotency, primed pluripotency and neural markers during differentiation of WT E14 mESC into AntNPCs at day 0 (D0), day 3 (D3) and day 5-day 6 (D5-D6). For each time point, the different heatmap columns represent biological replicates. **(D)** Left: Pie chart indicating the percentage of D0 ZIC2 peaks that are either proximal (<5 Kb away from a transcription start site (TSS)) or distal (>5kb from a TSS) to a TSS. Right: Pie chart subdividing the proximal and distal D0 ZIC2 peaks into CGI+ (peaks located <3 Kb from a CGI (see Methods)), and CGI- (peaks located >3 Kb from a CGI). **(E)** Heatmap plots of aggregated ChIP-Seq (for the indicated histone marks) and ATAC-seq signals in WT and *Zic2*^*-/-*^ mESC around (+/- 1Kb) ZIC2 peaks identified in mESC (D0). The ZIC2 peaks are separated in four groups based on their proximity to genes (proximal or distal) and CGI (CGI+ or CGI-). ChIP-seq signals are shown as RPKMs (see Methods for normalization details). **(F)** Genome browser snapshots illustrating ZIC2 binding at representative poised (top) and active (bottom) enhancers. CGIs, ZIC2, H3K27ac and H3K27me3 ChIP-Seq profiles in mESC, and NCBI RefSeq Curated gene annotations (GRCm38/mm10) are shown around the representative enhancers. **(G)** Brightfield microscopy images (4X) of WT and *Zic2*^*-/-*^ mESC cultured in S+LIF media. **(H)** Alkaline Phosphatase Staining images of WT mESC and *Zic2*^*-/-*^ mESC. Abbreviations: gRNA (guide RNA), Chr (Chromosome), AntNPCs (Anterior neural progenitor cells), Diff. (Differentiation), CGI (CpG island).

### Selection of neural differentiation time points for the genomic characterization of ZIC2 function

To interrogate ZIC2 function during neural induction, the generated mESC lines were combined with a differentiation protocol whereby mESC are differentiated into neural progenitors with anterior (i.e. forebrain/midbrain) identity (AntNPC) over a six-day period (28, 31–33) (Figure 1 B). Next, genomic data (i.e. RNA-seq, ChIP-seq and ATAC-seq) were generated at three different time points that, based on gene expression profiling, should capture major neural differentiation stages (Figure 1 C): (i) Day 0 (D0) corresponds to mESC cultured in S+L conditions (Serum + LIF) and is characterized by the expression of naive pluripotency markers such as *Sox2, Zfp42, Nanog* and *Klf4* (34–37) ; (ii) Day 3 (D3) corresponds to the exit from pluripotency during which cells express high levels of primed/formative pluripotency markers, such as *Fgf5, Dnmt3b* and *Pim2*, but low levels of somatic specifiers (38); (iii) Finally, Day 5 (D5) and Day 6 (D6) correspond to neural induction, capturing the expression of neural and midbrain/forebrain markers. For each differentiation time point, RNA-seq, ATAC-seq, and ChIPseq data for different histone marks associated to active and repressed/poised states (H3K27ac, H3K4me1, H3K4me2, H3K4me3 and H3K27me3) were generated in WT and *Zic2*^*-/-*^ cells. Moreover, ZIC2 ChIP-seq experiments were performed in *Zic2*^*FLAG-HA*^ cells using an anti-HA antibody (Figure 1 B).

### *Zic2*^*-/-*^ mESCs do not display any major phenotypic, transcriptional or chromatin defects despite extensive ZIC2 genomic binding

To investigate the role of ZIC2 in pluripotent cells, ChIP-seq experiments were performed in *Zic2*^*FLAG-HA*^ mESCs. A total of 13.768 ZIC2 peaks were identified, which were highly enriched for the canonical ZIC binding motif (Figure S2 A and Supplementary Data 1). The majority (60%) of ZIC2 peaks were distal (>5 Kb from a TSS) (Figure 1 D), thus in agreement with the previously reported role of ZIC2 as an enhancer-binding TF (14). However, it should be noted that a significant fraction (40%) of ZIC2 binding sites occurred within proximal promoter regions. Furthermore, the majority of ZIC2-bound proximal regions (96%) and a significant fraction of the distal ones (36%) overlapped with CpG islands (CGI). The frequent overlap of distal ZIC2 peaks with CGI is in agreement with the previously reported enrichment of ZIC2 binding motifs within poised/CG-rich enhancers (32), a group of distal CREs associated with CGI and that play essential roles during the induction of major developmental genes (31, 39). To get more insights into the epigenetic state of the different enhancer and promoter elements bound by ZIC2, ATAC-seq and ChIP-seq profiles for active and repressive histone marks (H3K27ac, H3K4me1, H3K4me2, H3K4me3 and H3K27me3) generated in WT and *Zic2*^*-/-*^ mESC were examined (Figure 1 E). This revealed that ZIC2 binds both active and poised/CG-rich enhancers (Figure 1 E, F, Figure S2 B-D). Among the 2939 active enhancers (ATAC+, H3K27ac+, H3K4me1+) identified in mESC, 56% were bound by ZIC2, of which almost 50% were CGI+ (Figure S2 B). Moreover, when considering poised enhancers (p300+, H3K27me3+, H3K4me1+) previously identified across various pluripotent cell types (39), 24% of them were bound by ZIC2 (Figure S2 B). On the other hand, ZIC2 was bound to numerous active (H3K27ac+, H3K4me3+) and bivalent (H3K27ac-, H3K27me3+, H3K4me3+) promoters that, as stated above, were mostly CGI+ (Figure 1 E, F).

Despite the widespread binding of ZIC2 to thousands of active promoters and enhancers, *Zic2*^*-/-*^ mESCs did not show any obvious morphological alterations (Figure 1 G). Alkaline phosphatase staining indicated that *Zic2*^*-/-*^ mESCs maintained an undifferentiated state (Figure 1 H). Additionally, bulk RNA-seq analysis revealed no major transcriptional changes in the absence of ZIC2 (n DEGs=8; Figure S3 A and Supplementary Data 2). This includes normal expression levels of major pluripotency markers, further supporting that pluripotency remains unaffected in *Zic2*^*-/-*^ mESCs (Figure S3 B). Lastly, and in agreement with the lack of major transcriptional defects, chromatin accessibility as well as active and repressive histone marks were largely unaffected in *Zic2*^*-/-*^ mESC across both proximal and distal ZIC2-bound regions (Figure 1 E, Figure S3 C and Supplementary Data 1 (n Diff. Acc. Regions = 86)). Note that a mild decrease in H3K4me1/2 was actually observed. However, in the absence of major transcriptional or chromatin accessibility defects and considering that the functional relevance of H3K4me1/2 remains questionable (40), the importance of these changes in H3K4me1/2 was not further investigated.

### ZIC3 plays a dominant role in the maintenance of pluripotency and compensates for the loss of ZIC2 in mESCs

One possible explanation for the lack of transcriptional and chromatin defects in *Zic2*^*-/-*^ mESCs could be that the role of ZIC2 is compensated by other ZIC family members with either redundant or dominant functions in pluripotent cells. In this regard, recent publications reported an important role for ZIC3 in pluripotency both in mouse and humans (16, 19, 22), making it the strongest candidate to compensate for the loss of ZIC2. To test this hypothesis, we generated hemizygous *Zic3*^*-/Y*^ mESC lines (Figure 2 A and Figure S4) and compared them to *Zic2*^*-/-*^ and WT mESCs. In our mESC *in vitro* system, *Zic3* expression levels were the highest of all the *Zic* members expressed (Figure 2 B), further supporting an important role for ZIC3 in S+L pluripotency conditions. *Zic3*^*-/Y*^ mESC lines showed evident morphological defects, including the presence of cells with a differentiated appearance that colonized the gaps between typical mESC colonies (Figure 2 C). Alkaline phosphatase staining confirmed that pluripotency was compromised in *Zic3*^*-/Y*^ lines (Figure 2 D). Furthermore, RNA-seq profiling revealed pronounced transcriptional differences between *Zic3*^*-/Y*^ mESC and either WT or *Zic2*^*-/-*^ mESC (Figure S5 A-C, Figure 2 E and Supplementary Data 2). These included the upregulation of genes involved in cell migration and morphogenesis, indicative of spontaneous differentiation, as well as the downregulation of genes associated with germline development and pluripotency (e.g., *Nanog, Klf4, Esrrb, Dazl, Ddx4*) (41–43) (Figure 2 E,F; Figure S5 B-C). A closer examination of the RNA-seq data revealed that the decreased expression of major pluripotency markers in *Zic3*^*-/Y*^ mESC was accompanied by an increase in the expression levels of endodermal genes (e.g., *Gata6, Sox17, Foxa2*) (Figure 2 E,F). Overall, our data are in agreement with previous studies indicating that *Zic3*^*-/Y*^ mESC can not maintain their pluripotency and show spontaneous differentiation, particularly towards the endodermal lineage (19).

**Fig. 2.**
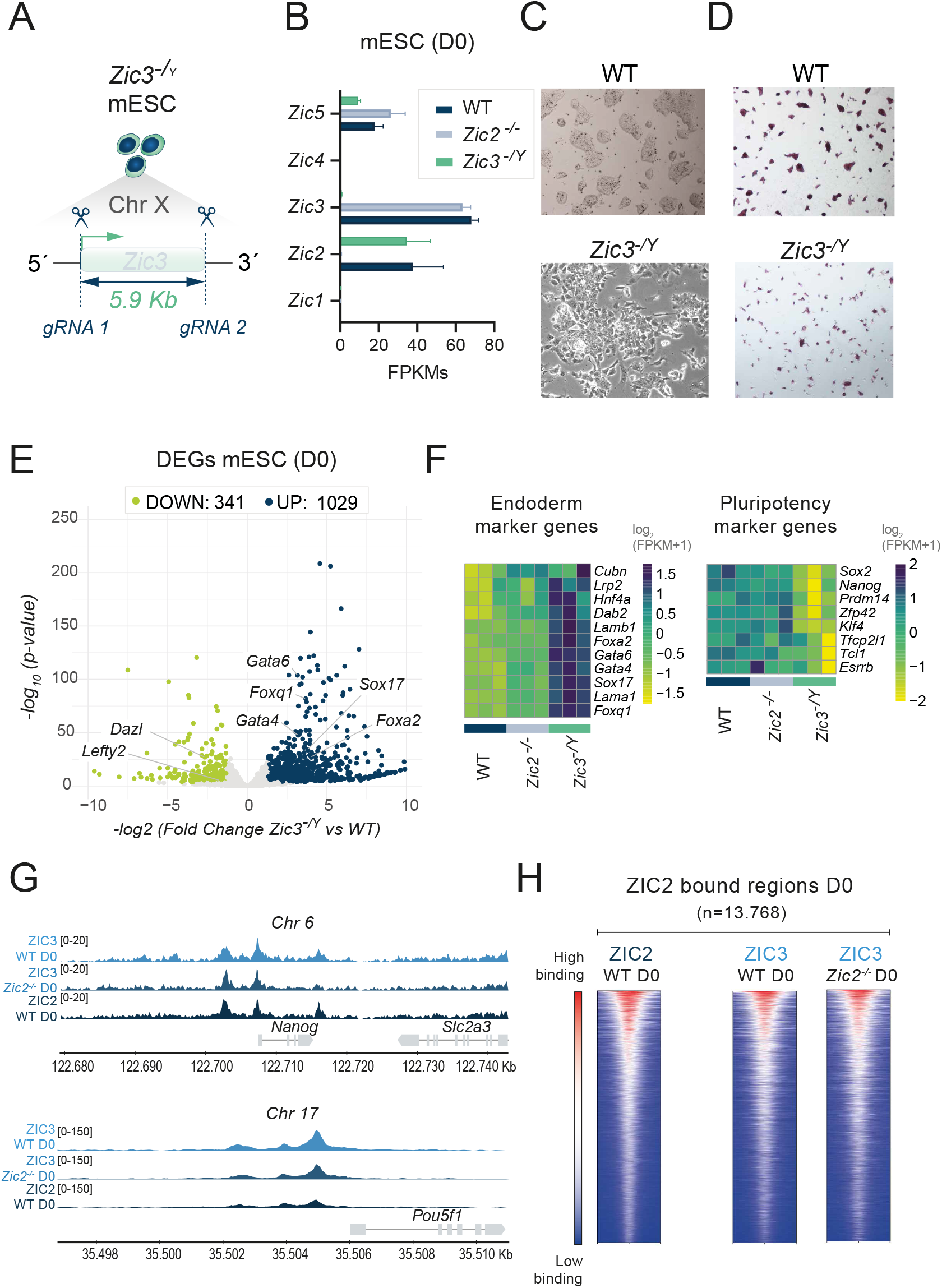
ZIC3 safeguards pluripotency and compensates for ZIC2 deficiency in mESCs. **(A)** Schematic view of the hemizygous *Zic3*^*-/Y*^ mESC line containing a 5.9 Kb deletion spanning the entire *Zic3* gene. **(B)** Barplot showing the expression levels, as measured by RNA-Seq expression (FPKM), of the *Zic* gene family members (i.e *Zic1-Zic5*) in WT, *Zic2*^*-/-*^ and *Zic3*^*-/Y*^ mESCs (D0). **(C)** Brightfield microscopy images (4X) of WT and *Zic3*^*-/Y*^ mESC cultured in S+LIF media. The WT image is the same as in Fig. 1G. **(D)** Alkaline Phosphatase Staining images of WT mESC and *Zic3*^*-/Y*^ mESC. The WT image is the same as in Fig. 1H. **(E)** Volcano plot of differentially expressed genes (DEGs) in *Zic3*^*-/Y*^ versus WT mESCs (D0). Significantly upregulated (blue) and downregulated (green) genes in *Zic3*^*-/Y*^ mESC are highlighted. **(F)** Heatmap plot showing the expression levels (as measured by RNA-seq; log_2_(FPKM + 1)) of selected lineage-specific markers in WT mESC (three biological replicates), *Zic2*^*-/-*^ mESC (three biological replicates for the *Zic2*^*-/-*^ mESC clonal line #1) and *Zic3*^*-/Y*^ mESC (one replicate for each of the three generated *Zic3*^*-/Y*^ clonal mESC lines (#4, #11, and #15)). Left: endoderm-associated genes; right: pluripotency-associated genes. **(G)** Genome Browser snapshots showing ZIC2 and ZIC3 ChIP-Seq profiles in WT and *Zic2*-/- mESCs around major representative genes (e.g., *Nanog, Pou5f1*). **(H)** Heatmap plots of aggregated ZIC2 ChIP-seq signals in WT mESC (left panel) and ZIC3 ChIP-seq signals in WT and *Zic2*^-/-^ mESCs (central and right panels) around all the ZIC2 peaks (n=13768) identified in WT mESCs (Fig. 1D). In all three heatmaps, the ZIC2 peaks are ranked by the ZIC2 ChIP-seq signals in WT mESC from high (red) to low (white/blue) signals. ChIP-seq signals are shown as RPKMs.

The phenotypic and transcriptional defects of the *Zic3*^*-/Y*^ mESC supports the notion of ZIC3 playing a dominant role among ZIC TFs in the maintenance of pluripotency, thus being able to compensate for the loss of ZIC2 in mESC. To better understand how these compensatory mechanisms might occur, we first evaluated *Zic3* RNA and protein levels in *Zic2*^*-/-*^ mESCs. These analyses revealed that the compensatory role of ZIC3 does not involve any major changes in ZIC3 expression levels upon loss of ZIC2 (Figure 2 B and Figure S5 D-E). Next, ChIP-seq experiments were performed in WT and *Zic2*^*-/-*^ mESC using a ZIC3 specific antibody (Figure S6 A-C). Comparison of the ZIC2 and ZIC3 ChIP-seq data showed that these two TFs bind very similar regions genome-wide, including regulatory elements linked to major pluripotency genes (e.g., *Nanog*), and that the binding profiles of ZIC3 do not change in the absence of ZIC2 (Figure 2 G,H). Therefore, the dominant function of ZIC3 in mESC does not involve a unique set of ZIC3-bound CREs, but rather a common set of CREs that are also bound by ZIC2.

Lastly, the strong similarity between the ZIC2 and ZIC3 binding profiles in mESC made us wonder whether ZIC2 could still play a regulatory function in pluripotent cells that, given the dominant role of ZIC3, might only become evident in the absence of ZIC3. Therefore, we aimed at deleting *Zic3* in the *Zic2*^*-/-*^ mESC in order to generate *Zic3*^*-/Y*^*/ Zic2*^*-/-*^ double *knock-out* cells, which according to our hypothesis, should display more severe defects than any of the single *knock-outs*. Although the *Zic3* deletion was readily detected in the transfected population of *Zic2*^*-/-*^ mESC, we were unable to derive stable *Zic3*^*-/Y*^*/Zic2*^*-/-*^ mESC clones (i.e. >40 clones were genotyped; Figure S7), indicating that self renewal and/or cell viability was compromised in the absence of both ZIC2 and ZIC3. These results suggest that, as recently reported in humans (16), ZIC2 and ZIC3 play partially redundant regulatory functions in mouse pluripotent cells, with ZIC3 being the most important ZIC family member at this developmental stage.

### ZIC2 is dispensable for both the exit from pluripotency and the induction of primed pluripotency genes

Having shown that ZIC2 plays a rather minor role in mESC, we next asked whether its regulatory function becomes more prominent during the exit from pluripotency, as cells acquire a primed pluripotent state. In agreement with this possibility, upon differentiation of mESC into AntNPCs, *Zic2* expression increased considerably by D3, while *Zic3* levels decreased (Figure 3 A). Moreover, the elevated expression of *Zic2* on D3 was accompanied by an increase in the number of ZIC2 binding sites (n=29.819) (Figure 3 A, B; Figure S8 A, B; Supplementary Data 1), which show a similar distribution with respect to gene TSS and CGI as the one observed in mESC (Figure S8 C, D). Interestingly, approximately two thirds (n=19.951) of the ZIC2 peaks on D3 appeared *de novo* (Figure 3 B), while the remaining ones were already bound in mESC. Furthermore, many of the distal ZIC2 peaks overlapped with putative CREs, as ZIC2 binds to 65% of the D3 active enhancers (Figure S8 E) and 53% of the poised enhancers previously mapped in mouse pluripotent cells (39) (Figure S8 F), further supporting that ZIC2 could play an important role as cells acquire a primed pluripotent state (15). However, despite the high expression and extensive genomic binding that ZIC2 displayed by D3, *Zic2*^*-/-*^ mESC differentiated successfully into D3 progenitors with no evident proliferation or morphological defects (Figure 3 C). Accordingly, RNA-seq profiling revealed rather moderate gene expression changes in D3 *Zic2*^*-/-*^ cells, with 195 upregulated and 155 downregulated genes in comparison to their WT counterparts (Figure 3 D; Supplementary Data 2). Importantly, the silencing of naive pluripotency markers (i.e: *Nanog, Klf4, Prdm14, Pou5f1*) and the induction of primed pluripotency genes (i.e: *Pou3f1, Otx2, Foxd3, Zfp281* (44, 45) proceeded normally in the absence of ZIC2 (Figure S9 A). Overall, our initial characterization of D3 *Zic2*^*-/-*^ cells indicates that ZIC2 is not necessary for either the exit from pluripotency or the establishment of the primed pluripotency regulatory network, thus contradicting previous studies based on shRNA-mediated *Zic2 knock-down* (14) and ZIC2 ChIP-seq profiling (15), respectively.

**Fig. 3.**
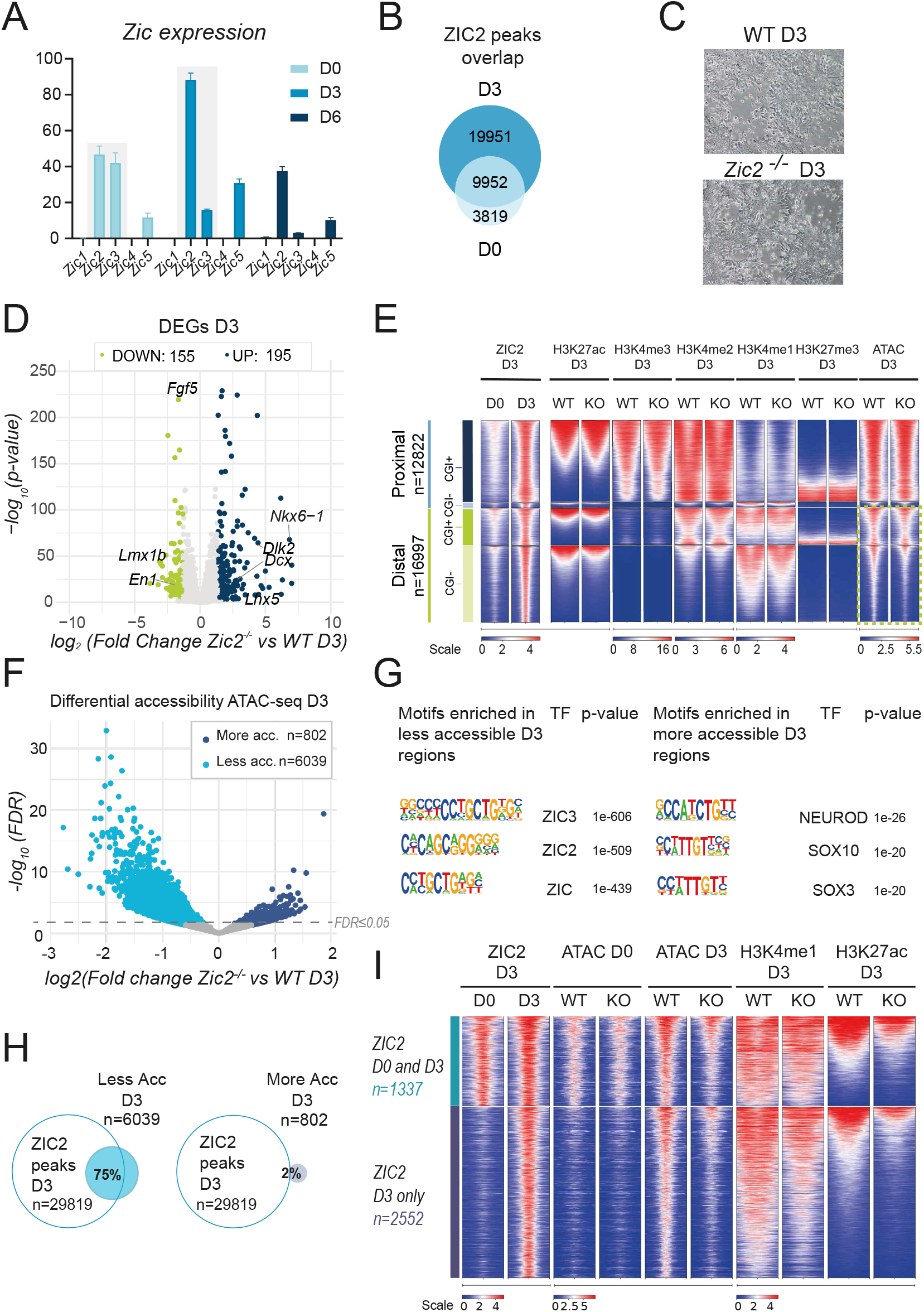
ZIC2 is dispensable for pluripotency exit but acts as pioneer TF promoting chromatin accessibility at putative neural enhancers. **(A)** Barplot showing the expression levels (as measured by RNA-Seq (FPKM)) of all the *Zic* gene family members (i.e *Zic1-Zic5*) in WT cells at the indicated differentiation time points: day 0 (D0), day 3 (D3) and day 6 (D6). **(B)** Venn diagram showing the overlap between the ZIC2 peaks identified at D0 and D3. **(C)** Brightfield images (10X) of WT and *Zic2*^*-/-*^ D3 cells. **(D)** Volcano plot of differentially expressed genes (DEGs) in *Zic2*^*-/-*^ versus WT D3 cells. Significantly upregulated (blue) and downregulated (green) genes in D3 *Zic2*^*-/-*^ cells are highlighted. **(E)** Heatmap plots of aggregated ChIP-seq (for the indicated histone marks) and ATAC-seq signals in WT and *Zic2*^*-/-*^ D3 cells around (+/- 1Kb) the ZIC2 peaks identified on D3. The ZIC2 peaks are separated into four groups based on their proximity to genes (proximal or distal) and CGI (CGI+ or CGI-). ChIP-seq and ATAC-seq signals are ranked by the H3K27ac signals in WT D3 cells and are shown as RPKMs (see Methods for normalization details). **(F)** Volcano plot showing genomic regions with differential accessibility in *Zic2*^*-/-*^ vs WT D3 cells. **(G)** Overrepresented motifs identified using HOMER (Heinz et al., 2010) in regions showing either less (left) or more (right) chromatin accessibility in D3 *Zic2*^*-/-*^ cells in comparison to D3 WT cells. **(H)** Venn diagram showing the overlaps between D3 ZIC2 peaks and regions with either less (left) or more (right) chromatin accessibility in D3 *Zic2*^*-/-*^cells in comparison to D3 WT cells (**I)** Heatmap plots of aggregated ATAC-seq and ChIP-seq signals for ZIC2, H3K4me1 and H3K27ac are shown in WT and *Zic2*^*-/-*^ cells at both D0 and D3 around (+/- 1 Kb) D3 ZIC2 peaks losing chromatin accessibility in D3 *Zic2*^*-/-*^ cells . The D3 ZIC2 peaks are classified into two groups: peaks exclusively present at D3 (ZIC2 D3 only; n=2552) and peaks already present at D0 (ZIC2 D0 and D3; n=1337). ChIP-seq and ATAC-seq signals are ranked by the H3K27ac signals in WT D3 cells and are shown as RPKMs (see Methods for normalization details).

### ZIC2 confers accessibility to putative neural enhancers upon pluripotency exit

Upon closer inspection of the D3 RNA-seq data, we noticed that the upregulated genes in *Zic2*^*-/-*^ cells were enriched in terms related to brain development and neural differentiation, while such enrichments were not detected in downregulated genes (Figure S9 B-C). Among the downregulated (e.g., *En1* (46–48), *Lmx1b* (49, 50)) and upregulated genes (e.g., *Nkx6-1* (51)) there were a few neural patterning regulators (Figure 3 D and Supplementary Data 2). However, the biological relevance of these gene expression changes on D3 is questionable, as these patterning genes showed a rather low and incipient expression in WT D3 cells before getting fully induced on D6 (Figure S9 D). Nevertheless, we questioned whether these subtle gene expression changes in D3 *Zic2*^*-/-*^ cells could be an indication of more severe chromatin defects whose effects on gene expression would only become strongly manifested at later differentiation stages. To explore this possibility, we examined the ATAC-seq and ChIP-seq profiles for active and repressive histone marks (H3K27ac, H3K4me1, H3K4me2, H3K4me3 and H3K27me3) around the D3 ZIC2 peaks in both D3 WT and *Zic2*^*-/-*^ cells. Although no major changes could be observed around ZIC2 proximal peaks, we noticed that a significant fraction of distal ZIC2 peaks displayed a loss of chromatin accessibility (as measured by ATAC-seq), and to a lesser extent of H3K4me1, in D3 *Zic2*^*-/-*^ cells (Figure 3 E). Next, to further explore the chromatin accessibility changes, differential accessibility analysis was performed between *Zic2*^*-/-*^ and WT D3 cells. Notably, there were 6039 genomic regions showing a loss of chromatin accessibility in *Zic2*^*-/-*^ cells, while only 802 regions were more accessible in comparison to WT cells (Figure 3 F and Supplementary Data 1). Furthermore, the majority of the differentially accessible regions were distal (87%), thus representing putative enhancers, and CGI- (72%) (Figure S10 A). To evaluate whether ZIC2 could be directly involved in controlling the accessibility of these regions, we first performed a motif enrichment analysis (Figure 3 G). Remarkably, the ZIC motif was strongly enriched among the regions losing accessibility in *Zic2*^*-/-*^ cells, but not in the regions gaining accessibility (Figure 3 G). Furthermore, and in full agreement with the motif analysis results, according to our ZIC2 ChIP-seq data in D3 WT cells, 75% of the less accessible regions in D3 *Zic2*^*-/-*^ cells were bound by ZIC2, while only 2% of the more accessible ones overlapped with ZIC2 peaks (Figure 3 H). These results strongly suggest a direct role for ZIC2 in promoting chromatin accessibility at distal regulatory elements as mESC transition to a primed pluripotent state.

To get additional insights into the potential function of the distal regions whose accessibility is directly controlled by ZIC2 in D3 cells, we next examined their chromatin state in both D0 and D3 (Figure 3 I). Interestingly, 65% of these distal regions were not initially bound by ZIC2 in D0 and, concomitantly with the binding of ZIC2 in WT D3 cells, these regions gained chromatin accessibility. Moreover, most of these regions were enriched in H3K4me1 in D3 cells, while only a small fraction was marked with H3K27ac, suggesting that they preferentially represent primed/poised enhancers whose accessibility and, to a lesser extent, H3K4me1 levels, are regulated by ZIC2 as mESC exit pluripotency (Figure 3 I). In agreement with this, functional annotation of the D3 ZIC2-bound regions losing accessibility in *Zic2*^*-/-*^ D3 cells revealed that they were preferentially associated with genes involved in brain development/patterning (e.g., *En1, En2, Lmx1a, Lmx1b, Msx1* (49, 50, 52–54)) and WNT signaling (e.g., *Wnt1, Wnt3, Wnt4, Wnt7* (55))(Figure S10 B, C) and that, in general, are either inactive or lowly expressed in both D0 and D3 cells (Figure S10 D). Overall, these findings suggest that ZIC2 binds to enhancers associated with genes involved in brain development/patterning and crucial signaling pathways (e.g., WNT) already on D3 (i.e. primed/formative pluripotency), acting as a pioneer TF that provides chromatin accessibility and primes these enhancers for their subsequent activation upon neural induction.

### The pioneering activity of ZIC2 at putative neural enhancers occurs widely across primed pluripotent cells

During developmental transitions, pioneer TFs can drive changes in chromatin accessibility that precede future changes in gene expression, creating poised/primed chromatin states that prepare genes for future activation (56–60). In principle, pioneer TFs can control chromatin accessibility during cell fate transitions according to two major modes of action: i) Differential binding: pioneer TFs broadly open cis-regulatory elements (mainly enhancers) in progenitor cells, which are then activated in a lineage-specific manner through the subsequent combinatorial binding of additional lineage-specific TFs; ii) Differential accessibility: pioneer TFs, alone or in a combinatorial manner (60–62), might promote CRE accessibility in a subset of progenitor cells in which those CREs will be subsequently activated upon binding of additional TFs (58, 63). These two alternative scenarios are difficult to discern using bulk genomic measurements. In addition, since our *in vitro* differentiation system leads to the induction of genes at D5-D6 that are known to define distinct, sometimes mutually exclusive neural identities *in vivo* (e.g., *Lmx1a* expressing cells do not express *Nkx6*.*1 in vivo*), we anticipated a certain degree of heterogeneity in the differentiated populations. Taking all this into account, we sought to determine how ZIC2 executes its pioneering role by performing a single-cell Multiome (ATAC + RNA) characterization of D3 progenitors (59).

After quality control filtering, Multiome data were obtained for 5599 WT and 2766 *Zic2*^*-/-*^ D3 cells (Figure S11 A, B). Firstly, we analyzed the scRNA-seq data, which after data integration and UMAP visualization (see Methods section), showed no specific clusters for either WT or *Zic2*^*-/-*^ D3 cells (Figure 4 A, B). This suggests that WT and *Zic2*^*-/-*^ D3 cells show rather similar identities, thus in agreement with the minor transcriptional differences detected in the previous bulk RNA-seq analyses (Figure 3). Furthermore, analyses of key naive and primed pluripotency markers showed that both WT and *Zic2*^*-/-*^ cells were able to silence the naive pluripotency program and acquire a primed state with similar dynamics (Figure 4 C). Our previous bulk RNA-seq data on D3 indicated that a few neural patterning genes showing an incipient, albeit low, expression in D3 WT cells were misregulated in the absence of ZIC2. Inspection of the scRNA-seq data revealed that the expression of most of these genes (i.e: *En1, Lmx1a, Nkx6-1, Dlk2, Dcx*) was barely detectable in either WT or *Zic2*^*-/-*^ D3 cells (Figure S11 C). One notable exception was *Lmx1b*, as this gene was expressed in a significantly larger fraction of WT than *Zic2*^*-/-*^ D3 cells (Figure S11 C). Overall, although these observations could be partly attributed to the sparsity of the scRNA-seq data (64), they further support that, up to D3, neural differentiation progresses properly in the absence of ZIC2.

**Fig. 4.**
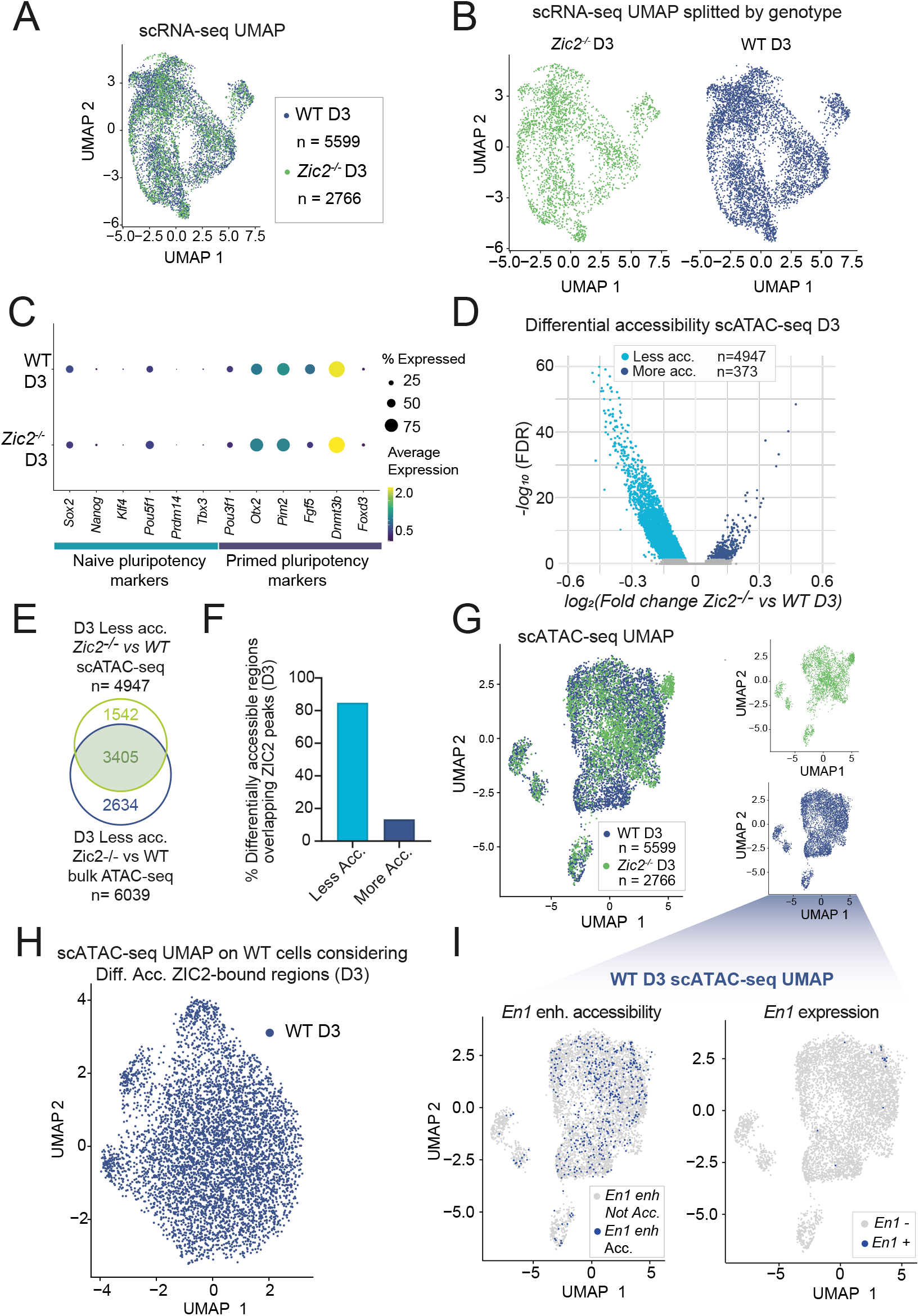
Single-cell Multiome (RNA-seq and ATAC-seq) profiling indicates that ZIC2 is a promiscuous TF that broadly pioneers neural enhancers in primed pluripotent cells. **(A)** UMAP plot of integrated scRNA-seq data generated for WT (blue, *n* = 5.599) and *Zic2*^*-/-*^ (green, *n* = 2.766) D3 cells. **(B)** UMAP plots from (A) split by genotype. **(C)** Dot plot showing the expression of selected naive and primed pluripotency markers in WT and *Zic2*^*-/-*^ D3 cells. The dot size indicates the percentage of expressing cells, and the dot color the average expression levels. **(D)** Volcano plot displaying differentially accessible chromatin regions between *Zic2*^*-/-*^ and WT D3 cells, based on single-cell ATAC-seq data. Regions with significantly reduced (blue, n = 4.947) or increased (light blue, n = 373) accessibility in *Zic2*^-/-^ cells are highlighted. **(E)** Venn diagram comparing regions displaying reduced accessibility in *Zic2*^*-/-*^ versus WT D3 cells according to either scATAC-seq (top) or bulk ATAC-seq (bottom) data. **(F)** Percentage of differentially accessible regions (based on scATAC data, Figure 4 D) overlapping ZIC2-bound regions identified in D3 cells by ChIP-seq (Figure 3 B). **(G)** UMAP plot of integrated scATAC-seq data generated for WT (blue, *n* = 5.639) and *Zic2*^*-/-*^ (green, *n* = 2.598) D3 cells. The right insets show UMAP plots split by genotype. **(H)** UMAP plot for D3 WT cells generated considering only the scATAC-seq data at the D3 ZIC2-bound regions with altered chromatin accessibility in *Zic2*^*-/-*^ D3 cells. **(I)** scATAC-Seq UMAP plots of D3 WT cells (WT inset in (G)) highlighting cells according to chromatin accessibility of an *En1* putative enhancer and expression levels of *En1*. Left: Cells in which the *En1* putative enhancer (chr1 120643940 120644846 mm10) is considered to be accessible (*≥* 1 read count from scATAC-Seq data) are marked in blue (*En1* enh Acc.). Right: Cells in which *En1* expression is detectable (*≥* 1 read count from scRNA-Seq data) are marked in blue (*En1*+). Abbreviations: enh (enhancer), Acc. (accessible).

Next, we examined the scATAC-seq data generated in the WT and *Zic2*^*-/-*^ D3 cells. Reassuringly, the strong bias towards accessibility losses detected in D3 *Zic2*^*-/-*^ cells according to bulk ATAC-seq was also observed in the single-cell data (Figure 4 D). Furthermore, there was a large overlap between the regions losing chromatin accessibility in *Zic2*^*-/-*^ cells according to either scATAC-seq or bulk ATAC-seq (Figure 4 E). Importantly, 84.7% of the less accessible regions in *Zic2*^*-/-*^ cells were bound by ZIC2 in WT D3 cells, while only 13.4% of the regions gaining accessibility in *Zic2*^*-/-*^ cells were bound by ZIC2 (Figure 4 F). These findings further support the notion of ZIC2 acting as a pioneer TF that provides accessibility to putative enhancers that might get active at subsequent stages of neural differentiation. To evaluate whether ZIC2 broadly influences chromatin accessibility within D3 progenitors or, alternatively, this preferentially occurs within specific D3 cell sub-populations, we performed integration and UMAP visualization of the scATAC-seq data. Similarly to the scRNA-seq findings, no major differences between WT and *Zic2*^*-/-*^ D3 cells were observed (Figure 4 G). As ZIC2 controls the accessibility of only a relatively small fraction of all the D3 ATAC-seq peaks (among the 250.203 peaks considered in scATAC-seq analyses only 4244 are differentially accessible and bound by ZIC2), we then explored whether those regions could be particularly important in defining distinct WT D3 sub-populations based on chromatin accessibility. However, UMAP visualization of WT D3 cells considering only these ZIC2-bound regions did not reveal any distinct cellular cluster (Figure 4 H). Accordingly, visualization of chromatin accessibility levels at representative ZIC2-bound enhancers (e.g., *En1*-associated enhancer) showed that these regulatory elements were broadly accessible across WT cells, rather than within specific subpopulations, regardless of the low % of cells expressing their putative target genes (e.g., *En1*) (Figure 4 I right UMAP). Overall, our Multiome analyses suggest that, rather than controlling the accessibility of putative neural enhancers within distinct sub-populations of D3 progenitor cells, ZIC2 broadly and similarly pioneers these enhancers across D3 cells.

### *Zic2*^*-/-*^ AntNPCs show defective induction of brain patterning genes and premature neuronal differentiation

The previous analyses on D3 cells suggest that ZIC2 primes enhancers for their future activation upon neural induction (i.e. D5-6 in our AntNPC differentiation), which in turn might be important for the proper expression of major brain developmental genes (see, e.g., Figure 1 C, Figure S9 D and S10 D). Furthermore, this possibility is supported by the fact that, by D5-6, *Zic2* is the only *Zic* gene that remains highly expressed (Figure 3 A) and, thus, other ZIC family members are unlikely to compensate for its regulatory function in *Zic2*^*-/-*^ AntNPC.

To start evaluating the previous hypothesis, bulk RNA-seq and scMultiome (ATAC + RNA) experiments were conducted in WT and *Zic2*^*-/-*^ AntNPCs. In agreement with ZIC2 having an important regulatory function during neural induction, differential expression analysis of the bulk RNA-seq data revealed that, by D5-6, the number of DEGs was considerably larger than at previous differentiation stages (Figure 5 A and Supplementary Data 2). Functional enrichment analysis (65) (see Methods and Figure S12 A, B) and detailed examination of the DEGs (Figure 5 A) revealed that neural ventral markers (e.g., *Nkx6-1, Nkx6-2*) (51, 66) as well as markers of post-mitotic neurons (e.g., *Nhlh1, Dcx* (67–70)) (Figure 5 A) were upregulated in D5-6 *Zic2*^*-/-*^ AntNPC. The premature neuronal differentiation of D5-6 *Zic2*^*-/-*^ AntNPC was further supported by the presence of neurite-like projections (Figure S13 A) and the abundance of cells staining positive for the pan-neuronal marker TUJ1^+^ (71, 72) in D6 *Zic2*^*-/-*^ AntNPC in comparison to their WT counterparts (Figure S13 B). Conversely, key midbrain and roof plate markers, including *Lmx1a/b* (49, 50, 52, 53) *Wnt3a* (73, 74), *En1, Pax2* and *Wnt1* (46–48, 50, 75–78), were significantly downregulated in D5-6 *Zic2*^*-/-*^ AntNPC (Figure 5 A).

**Fig. 5.**
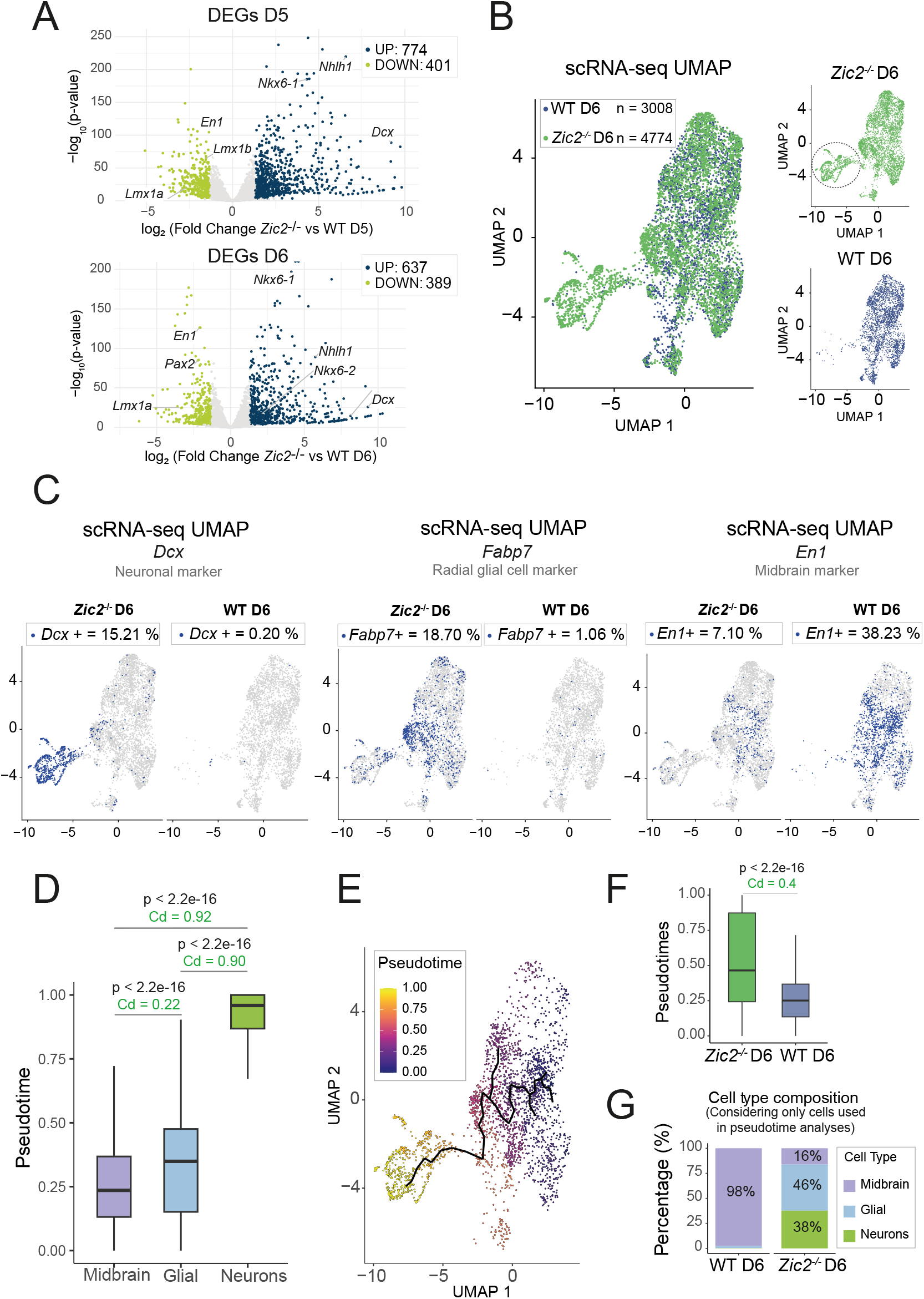
*Zic2*^-/-^ AntNPCs (D6) display impaired patterning and premature neuronal differentiation. **(A)** Volcano plots showing differentially expressed genes (DEGs) at day 5 (D5; top)) and day 6 (D6; bottom) in *Zic2*^-/-^ versus WT AntNPCs according to bulk RNA-seq data. Upregulated genes are shown in blue and downregulated genes in green. **(B)** UMAP plot of integrated scRNA-seq data generated in WT (n = 3008) and *Zic2*^*-/-*^ (n = 4774) D6 AntNPCs. The right insets show UMAP plots split by genotype. In the *Zic2*^*-/-*^ UMAP, the dotted circle marks a distinct cell cluster that is not present in WT AntNPCs. **(C)** Expression of selected genes (*Dcx, Fabp7* and *En1*) is shown within the D6 AntNPC scRNA-seq UMAPs split by genotype. For each selected gene, the expressing cells (*≥* 1 read count) are marked in blue. **(D)** Boxplot showing the distribution of pseudotime values for cells labelled as (i) midbrain progenitors (*En1*+), (ii) radial glial cells (*Fabp7* +), and (iii) post-mitotic neurons (*Dcx* +) (see Methods). **(E)** UMAP plot of integrated scRNA-seq data generated in WT and *Zic2*^-/-^ D6 AntNPC. Only cells considered for pseudotime analyses are depicted (midbrain progenitors, radial glial cells and post-mitotic neurons). Cells are coloured by pseudotimes and the inferred cellular trajectory is depicted with a tree. **(F)** Boxplot of pseudotime values, considering only the cells used for pseudotime analyses, splited by genotype (WT (right) and *Zic2*^*-/-*^ (left) D6 AntNPC). **(G)** Stacked bar plot showing the proportion of cells, considering only those used for pseudotime analyses, assigned in WT (left) and *Zic2*^*-/-*^ (right) D6 AntNPC to one of the following cell types: (i) midbrain progenitors (*En1*+), (ii) radial glial cells (*Fabp7* +), and (iii) post-mitotic neurons (*Dcx* +) (see Methods). In (D) and (F), the P-values were calculated using wilcox tests and the colored numbers beneath the p-values correspond to negligible (gray) and non-negligible (green) Cliff’s delta effect sizes.

To further refine how the previous gene expression changes translate into differences in cellular identities between WT and *Zic2*^*-/-*^ AntNPCs, the scRNA-seq data obtained as part of the scMultiome analyses performed in D6 AntNPCs were explored. Briefly, after quality control filtering, data for 3008 D6 WT and 4774 *D6 Zic2*^*-/-*^ cells were retrieved (Methods and Figure S14 A, B). Following data integration, UMAP visualization showed a major cluster containing both WT and *Zic2*^*-/-*^ cells and a smaller one with almost exclusively *Zic2*^*-/-*^ cells (Figure 5 B). This small cluster expressed high levels of post-mitotic neuronal markers (e.g., *Nhlh1, Dcx, Tubb3, Neurod1*) (Figure 5 C and Figure S14 D), further supporting that the loss of ZIC2 causes premature differentiation of AntNPC (Figure S13 B). Despite this premature differentiation, the majority of *Zic2*^*-/-*^ D6 cells still maintained an undifferentiated neural progenitor-like state (e.g., high levels of *Sox2* expression and no expression of post-mitotic neuronal markers (Figure S14 C, D)), as they were found, together with WT D6 cells, within the same large cluster. However, analysis of anterior-posterior (e.g., *En1, Lmx1b, Pax2*) and dorso-ventral (e.g., *Lmx1a, Lmx1b, Nkx6-1*) brain patterning genes that, according to the previous bulk RNA-seq data, were missregulated in *Zic2*^*-/-*^ AntNPCs, revealed that WT and *Zic2*^*-/-*^ neural progenitors significantly differed in their positional identity. Namely, in comparison to WT neural progenitor cells (NPCs), there was a strong reduction in the % of *Zic2*^*-/-*^ NPC expressing midbrain (e.g., *En1, Lmx1b, Pax2*) and roof plate (e.g., *Lmx1a, Wnt3a, Wnt1*) markers, while the % of *Zic2*^*-/-*^ NPC expressing ventral/basal markers increased (e.g., *Nkx6-1*) (Figure 5 C; Figure S14 E, F). Furthermore, the *Zic2*^*-/-*^ NPC acquiring more ventral identities also showed incipient signs of differentiation, as illustrated by the high expression of radial glial cell markers, such as *Fabp7* (79–82) (Figure 5 C).

Altogether, the bulk and scRNA-seq data suggest that, in the absence of ZIC2, a significant fraction of AntNPCs lose the expression of major midbrain and roof plate markers, including important WNT ligand genes (e.g., *Wnt3a, Wnt1*) and WNT signalling targets (e.g., *Axin2*) (Figure S14 F). WNT signaling in the neural tube promotes dorsal fates and prevents the premature differentiation of neural progenitors (83–90). Therefore, we hypothesize that, in the absence of ZIC2, the reduction in WNT signalling levels might in turn increase the % of AntNPCs displaying more ventral identities and undergoing premature neuronal differentiation (91). If this is correct, during our AntNPC differentiation, midbrain progenitors should sequentially differentiate into radial glia and post-mitotic neurons, and this differentiation trajectory should become accelerated in the absence of ZIC2. In order to test this hypothesis, we performed a pseudotime analysis with the D6 scRNA-seq data (Figure 5 D-F) (see Methods). First, we selected D6 cells that, based on the expression of *bona fide* markers, could be assigned to the following cell types: i) Postmitotic Neurons (*Dcx* +), ii) Radial Glial Cells (*Fabp7* +), and iii) Midbrain progenitors (*En1* +) (see Methods). In agreement with their developmental timing *in vivo*, our analysis correctly assigned lower pseudotimes to the midbrain progenitors than to the radial glial cells and post-mitotic neurons (Figure 5 D). More importantly, visualization of the differentiation trajectories according to the estimated pseudotimes supported the initial differentiation of midbrain progenitors into radial glial cells, which then gave rise to post-mitotic neurons (Figure 5 E). In addition, when considering the pseudotimes of the cells according to their genotype, *Zic2*^*-/-*^ cells showed significantly larger pseudo-time values (i.e. more differentiated) than WT cells (Figure 5 F). Accordingly, the proportion of radial glial cells and post-mitotic neurons was considerably larger for *Zic2*^*-/-*^ cells than for WT cells (Figure 5 G). Overall, these findings further support the important role of ZIC2 in controlling the patterning of AntNPC and preventing their premature differentiation.

### ZIC2 directly controls the activation of enhancers linked to brain patterning genes, while its effects on neuronal differentiation are largely indirect

The previous transcriptional characterization of D6 AntNPCs points to an important regulatory function of ZIC2 during the induction of major brain developmental genes, particularly those providing positional information and controlling neuronal differentiation. However, such analyses cannot address (i) which of those genes (and associated CREs) are directly regulated by ZIC2 and (ii) whether the observed transcriptional defects are connected with the enhancer pioneering function of ZIC2 in D3/primed pluripotent cells. To start addressing these questions, we performed ChIP-seq experiments in D6 WT AntNPC and identified 19.935 ZIC2 peaks (Figure 6 A, Figure S15 A and Supplementary Data 1). The genomic distribution of the D6 ZIC2 peaks as well as their motif content were similar to those observed at earlier differentiation time points (Figure S15 B-D). In agreement with the pioneering function of ZIC2, analysis of the ZIC2 binding dynamics throughout AntNPC differentiation (D0, D3 and D6) showed that only 14% of the D6 peaks were *de novo*, whereas 80% were already present from D3, among which more than a third (36%) were also bound at D0 (Figure 6 A). Notably, 51% of the active enhancers called in D6 AntNPC (H3K27ac+, H3K4me1+ and ATAC+ in D6 AntNPC) were bound by ZIC2 at this stage (Figure S15 E). In addition, when considering pluripotency-associated poised enhancers bound by ZIC2 on D3 (n=2209, Figure S8 F), 60% remained bound by ZIC2 on D6 (Figure S15 F), among which 20% became active (i.e. gained H3K27ac on D6) (Figure S15 F). These findings not only support ZIC2’s role as a pioneer TF that already pre-marks in pluripotent cells a significant fraction of the active enhancers present in neural progenitors, but also suggest that, following its pioneering function in D3/primed pluripotent cells, ZIC2 remains bound to many of those enhancers that become active in AntNPC.

**Fig. 6.**
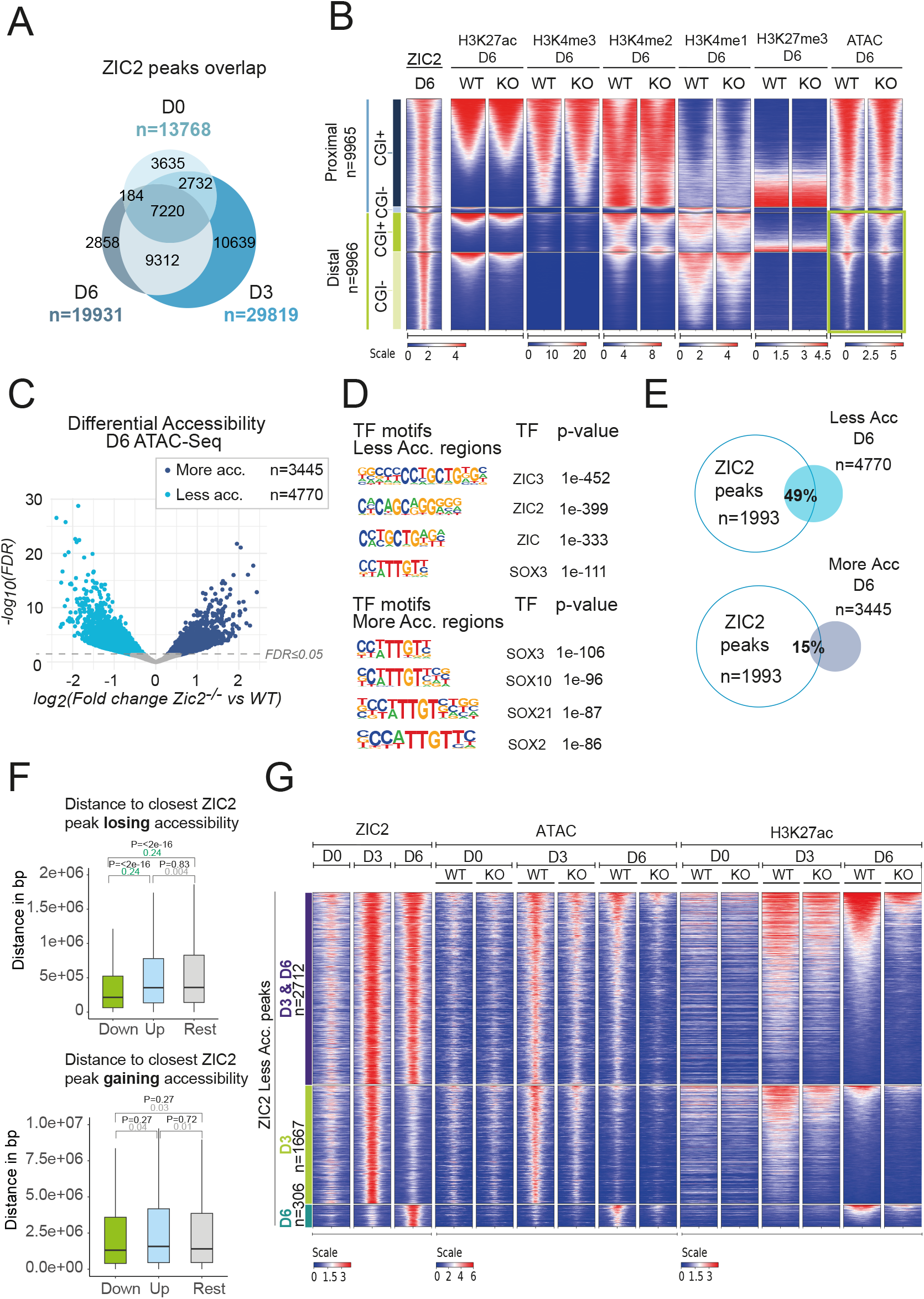
Previously primed enhancers linked to brain patterning genes are directly activated by ZIC2 in AntNPC (D6) . **(A)** Venn diagram showing the overlaps between ZIC2-bound regions identified by ChIP-seq at D0 (n = 13768), D3 (n = 29819), and D6 (n = 19331). **(B)** Heatmap plots of aggregated ChIP-seq (for the indicated histone marks) and ATAC-seq signals in WT and *Zic2*^*-/-*^ D6 AntNPC around (+/- 1Kb) the ZIC2 peaks identified on D6. The ZIC2 peaks are separated into four groups based on their proximity to genes (proximal if < 5kb from closest TSS, distal otherwise) and CGI (CGI+ or CGI-). ChIP-seq and ATAC-seq signals are ranked by the H3K27ac signals in WT D6 cells and are shown as RPKMs (see Methods for normalization details). **(C)** Volcano plot of differentially accessible chromatin regions (ATAC-seq) between *Zic2*^*-/-*^ and WT D6 AntNPC. Regions with decreased (n = 4770) or increased (n = 3445) accessibility in *Zic2*^*-/-*^ AntNPC are highlighted. **(D)** Overrepresented motifs identified using HOMER (Heinz et al., 2010) in regions showing either less (top) or more (bottom) chromatin accessibility in D6 *Zic2*^*-/-*^ AntNPC in comparison to D6 WT AntNPC. **(E)** Venn diagram showing the overlap between D6 ZIC2 peaks and the differentially accessible regions shown in (C). The percentage values indicate the fraction of regions displaying either more or less accessibility in D6 *Zic2*^*-/-*^ AntNPC that overlap with a D6 ZIC2 peak. **(F)** Boxplots generated with the distance from each of the genes downregulated (green), upregulated (blue) or unchanged (Rest; gray) in D6 and/or D5 *Zic2*^*-/-*^ AntNPCs with respect to their nearest ZIC2 peak either losing (top panel) or gaining (bottom panel) accessibility in D6 *Zic2*^*-/-*^ cells. P-values were calculated using wilcox test. The colored numbers beneath the p-values correspond to negligible (gray) and non-negligible (green) Cliff’s delta effect sizes. **(G)** Heatmaps displaying ZIC2 ChIP-seq, ATAC-seq, and H3K27ac ChIP-seq signals around (+/- 1Kb) D3 and D6 ZIC2 peaks that lose accessibility in *Zic2*^*-/-*^ D3 and/or D6 cells. The ZIC2 peaks are grouped according to the ZIC2 binding dynamics during neural differentiation:(i) regions bound both in D3 and D6 cells; (ii) regions bound exclusively in D3 cells; (iii) regions bound exclusively in D6 cells. Within each group, the ZIC2 peaks are ranked by the H3K27ac signal intensity in WT D6 cells. ChIP-seq and ATAC-seq signals are shown across the indicated time points (D0, D3, D6) and genotypes (WT and *Zic2*^-/-^).

Next, in order to evaluate whether ZIC2 was important for the proper activation of enhancers in D6 AntNPC, ChIP-seq and ATAC-seq profiles were generated in WT and *Zic2*^*-/-*^ D6 AntNPCs. As previously observed in D3 cells, distal ZIC2-bound regions exhibited a widespread loss of chromatin accessibility in *Zic2*^*-/-*^ cells, while proximal regions appeared largely unaffected (Figure 6 B). Interestingly, among the ZIC2-bound distal regions losing chromatin accessibility, those displaying an active state in WT D6 AntNPC (i.e. H3K27ac+ in WT D6 AntNPC) also showed decreased levels of H3K27ac and H3K4me2 in *Zic2*^*-/-*^ cells. In addition, the distal regions losing chromatin accessibility but displaying a primed rather than active state in WT cells (i.e. H3K4me1+ and H3K27ac- in WT D6 AntNPC) showed lower levels of H3K4me1 instead. Next, to directly identify those distal ZIC2-bound regions whose regulatory activity could be compromised in the absence of ZIC2, differential chromatin accessibility analysis was performed between *Zic2*^*-/-*^ and WT D6 cells. This analysis revealed that there were 4770 regions losing and 3445 regions gaining accessibility in *Zic2*^*-/-*^ D6 cells with respect to their WT counterparts, which, as expected, were mostly distal (Figure 6 C, Figure S16 A and Supplementary Data 1). Importantly, regions losing accessibility in *Zic2*^*-/-*^ D6 AntNPCs were highly enriched in ZIC motifs (Figure 6 D) and strongly overlapped with ZIC2 ChIP-seq peaks (49%) (Figure 6 E), supporting a direct role for ZIC2 in the establishment and/or maintainance of chromatin accessibility at these regions. In contrast, the regions gaining accessibility in *Zic2*^*-/-*^ AntNPCs were enriched for SOX rather than ZIC motifs (Figure 6 D) and, accordingly, only a small fraction of them (15%) were bound by ZIC2 (Figure 6 E). As SOX TF family members are involved in various processes related to neuronal and glial differentiation (92), the regions gaining chromatin accessibility in *Zic2*^*-/-*^ cells might represent a secondary effect (i.e. not directly mediated by ZIC2) associated to the premature differentiation of *Zic2*^*-/-*^ D6 AntNPCs. In agreement with this possibility, functional annotation of the differentially accessible regions identified in *Zic2*^*-/-*^ D6 AntNPCs (Figure S16 B, C) showed that while the genes linked to the regions losing accessibility in *Zic2*^*-/-*^ AntNPCs were associated with neural patterning (Figure S16 B), the regions gaining accessibility in *Zic2*^*-/-*^ AntNPCs were strongly associated with genes involved in neuronal differentiation and cell identity commitment, including several *Sox* genes (Figure S16 C). Most importantly, by evaluating the distance between genes and ZIC2-bound regions displaying changes in chromatin accessibility in *Zic2*^*-/-*^ AntNPCs, we observed that the genes downregulated in *Zic2*^*-/-*^ AntNPCs (D5 and/or D6) were significantly closer to ZIC2-bound regions losing accessibility in *Zic2*^*-/-*^ cells compared to either upregulated or non-differentially expressed genes (Figure 6 F). In contrast, when considering ZIC2-bound regions gaining accessibility in *Zic2*^*-/-*^ cells, no significant distance differences were appreciated between differentially and non-differentially expressed genes in *Zic2*^*-/-*^ AntNPCs (Figure 6 F). Overall, the previous ChIP-seq and ATAC-seq analyses indicate that ZIC2 directly binds to and promotes the accessibility and regulatory activity of enhancers controlling the expression of brain patterning genes. In contrast, the pre-mature neuronal differentiation observed in *Zic2*^*-/-*^ AntNPCs does not seem to be directly mediated by ZIC2 and might result from the downregulation of dorsal patterning and WNT signalling genes directly controlled by ZIC2.

### The majority of enhancers activated by ZIC2 in D6 AntNPC are already pioneered by ZIC2 on D3

To more directly explore to what extent the pioneering function of ZIC2 in D3/primed pluripotent cells could be important for enhancer activation in D6 AntNPC, all the ZIC2-bound regions losing accessibility in D3 and/or D6 *Zic2*^*-/-*^ AntNPCs were classified in three groups according to the ZIC2 binding dynamics during AntNPC differentiation: i) regions exclusively bound at D3 (n=1667), ii) regions exclusively bound at D6 (n=306) and iii) regions bound at both D3 and D6 (n=2712). Then, we evaluated chromatin accessibility and histone modifications at these three groups of regions in both WT and *Zic2*^*-/-*^ cells during AntNPC differentiation (Figure 6 G). This analysis revealed that strongly active enhancers in WT D6 AntNPC (i.e. high levels of H3K27ac in WT D6 AntNPC) displayed lower H3K27ac levels in D3 cells, while showing similar accessibility and ZIC2 binding levels in both D3 and D6 WT cells. Furthermore, these D6 active enhancers already showed reduced H3K27ac and ATAC-seq signals in D3 *Zic2*^*-/-*^ cells, and such reductions became even more pronounced in D6 Zic2^-/-^ cells. These results indicate that, by D3, ZIC2 binds to and controls the accessibility of the majority of enhancers whose subsequent activation in D6 AntNPC is ZIC2-dependent.

On the other hand, approximately half of the D3 & D6 ZIC2-bound regions as well as the majority of D3-only ZIC2-bound regions losing chromatin accessibility in *Zic2*^*-/-*^ cells, did not show high H3K27ac levels in either D3 or D6 WT cells. Moreover, most of these regions were more accessible in D3 than in D6 WT cells, especially those that were only bound by ZIC2 on D3 (Figure 6 G). Therefore, ZIC2 pioneers in D3/primed pluripotent cells a significant number of putative enhancers that do not get active under our differentiation conditions, but might do so in alternative developmental contexts. To evaluate this possibility, we intersected each of the three categories of ZIC2-bound regions losing accessibility in *Zic2*^*-/-*^cells with active enhancers mapped by the ENCODE consortium in multiple mouse embryonic tissues (93) (Figure S16 D; see Methods). Firstly, when considering active enhancers in E10.5-E11.5 forebrain and midbrain (i.e. Early brain enhancers) we found that the D3&D6 ZIC2 peaks showed a 3.6-fold higher overlap with those enhancers than the D3-only ZIC2 peaks (18% vs 5%). Nevertheless, these differences were substantially reduced when considering all active enhancers identified by the ENCODE consortium in additional mouse embryonic tissues *(i*.*e. All enhancers)*. Notably, the three groups of ZIC2-bound regions show high and similar overlaps with active embryonic enhancers currently annotated in the ENCODE database (>30%) (Figure S16 D). Overall, these observations support the notion that, in pluripotent cells, ZIC2 promiscuously primes enhancers that become subsequently active in multiple developmental stages and lineages.

### ZIC2 targeted degradation coupled with scMultiome profiling identifies the ZIC2-dependent regulatory network in neural progenitors

Our previous analyses suggest that, upon neural induction (i.e. D6 AntNPC), ZIC2 could directly activate a subset of enhancers required for the proper expression of major brain patterning and WNT signaling genes (e.g: *Lmx1a, Lmx1b, En1, Wnt1*). Notably, ZIC2 was bound to and promoted the accessibility of these enhancers already by D3, yet most of them only became fully active by D6 (Figure S17 A,B and Figure S18 A,B). Therefore, we wondered whether the activation of these enhancers was only dependent on the pioneering activity of ZIC2 in D3 cells or, whether, alternatively, its binding in D6 cells was still important. On the other hand, the results presented in previous sections also suggest that the genes upregulated in *Zic2*^*-/-*^ D6 AntNPC, especially those typical of post-mitotic neurons, do not represent direct ZIC2 targets. To address these questions and further refine the regulatory network directly controlled by ZIC2 during neural induction, we generated a ZIC2 degron mESC line using the d-TAG protein targeted degradation system (94), enabling us to deplete ZIC2 in an acute and inducible manner (Figure S19 A-C). Once we confirmed that the engineered 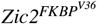 mESC line retained its capacity to differentiate into AntNPCs (Figure S19 D), we differentiated it under three different dTAG-13 treatment conditions (Figure 7 A and Figure S20 A, B): (1) continuous from D0 to D6, (2) during the final 48 hours (D4–D6), and (3) during the last 24 hours (D5–D6). The final 48 h and 24 h time points coincide with neural induction (Figure 1 A; Figure S19 D) and were selected to minimize indirect transcriptional defects associated with the prolonged absence of ZIC2. RNA-seq experiments were performed on D6 for the three ZIC2 depletion conditions (Figure 7 A). Principal component analysis (PCA) showed that the transcriptional differences between dTAG-13–treated and DMSO-treated (i.e. control) samples (Figure S20 A) increased with the duration of the ZIC2 depletion, reflecting a progressive accumulation of transcriptional changes. Differential expression analysis revealed 395 downregulated and 661 upregulated genes in the total depletion condition (D0–D6) (Figure S20 C). While this number of DEGs (Figure S20 C) was slightly lower than what was observed in *Zic2*^*-/-*^ cells (Figure 5 A), functional enrichment analysis (see Methods section) indicated that similar genes and gene categories were affected (i.e. downregulation of brain patterning genes (e.g., *En1, Lmx1a/b, Wnt3a*) and up-regulation of post-mitotic neuronal markers (e.g., *Neurog1, Dcx*)) (Figure S20 C, Figure S21 A vs Figures 5 A and S12 A,B). On the other hand, fewer DEGs were observed for the 48h (D4–D6) and 24h (D5–D6) depletion experiments (211 up/243 down and 101 up/187 down, respectively; Figure S20 D,E), something expected if indirect transcriptional defects are minimized under these late depletion conditions. Notably, the reduction in the number of DEGs in the short depletion experiments in comparison to the total depletion ones was considerably more pronounced for the upregulated genes (∼6.5-fold reduction) than for downregulated genes (∼2-fold reduction) (Figure 7 B). Furthermore, the genes downregulated in the short depletion experiments (i.e. down in 48h and/or 24h) were significantly closer to ZIC2 peaks losing chromatin accessibility in *Zic2*^*-/-*^ D6 AntNPCs than either the upregulated or non-differentially expressed genes, while such distance differences were not observed when considering ZIC2 peaks gaining accessibility in *Zic2*^*-/-*^ AntNPC (Figure 7 C). These results further support that ZIC2 directly activates brain patterning and WNT signaling genes, while indirectly repressing genes implicated in neuronal differentiation.

**Fig. 7.**
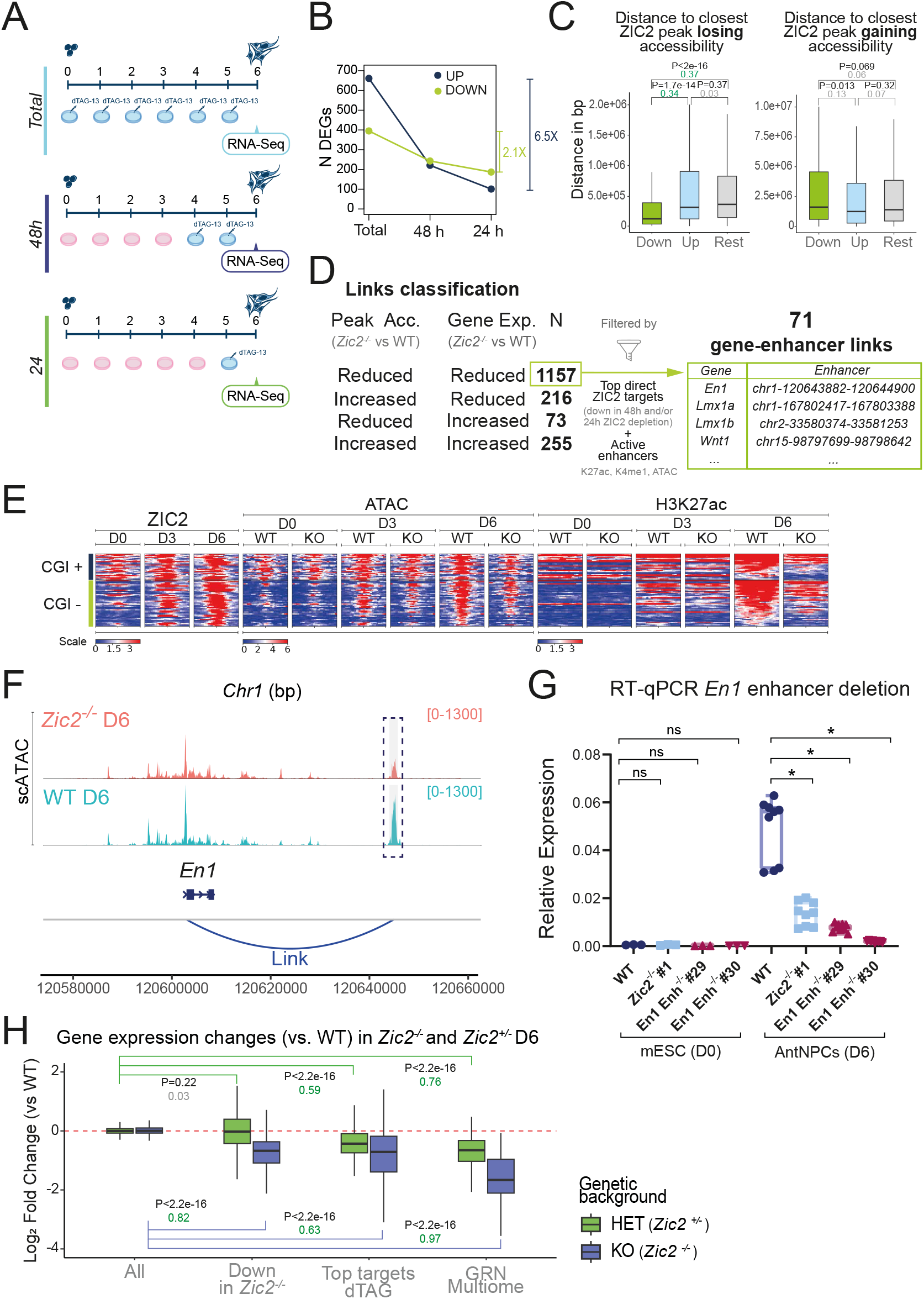
Identification of the ZIC2-dependent gene regulatory network in AntNPC using acute ZIC2 degradation and multiomic profiling. **(A)** Graphical summary of the experimental design used for the inducible degradation of ZIC2 using the dTAG system. 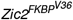 mESCs were differentiated into AntNPCs under three dTAG-13 treatment conditions: (i) Total: cells were treated from day 0 to day 6 (D0–D6), (ii) 48h: cells were treated for the final 48 h (D4–D6) (iii) 24h: cells were treated for the final 24 h (D5–D6). RNA-seq was performed on D6 for all three treatment conditions (i.e. Total, 48h and 24h). **(B)** Number of genes that are either upregulated (blue) or downregulated (green) upon ZIC2 depletion in comparison to DMSO-treated cells are shown for each of the dTAG-13 treatment conditions described in (A) (i.e. Total, 48h, 24h). **(C)** Boxplots generated with the distance from each of the genes downregulated (green), upregulated (blue) or unchanged (Rest; grey) in 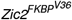 D6 AntNPCs treated with dTAG-13 (48h and/or 24h conditions) with respect to their nearest ZIC2 peak either losing (left panel) or gaining (right panel) accessibility in D6 *Zic2*^*-/-*^ cells. P-values were calculated using wilcox test. The colored numbers beneath the p-values correspond to negligible (gray) and non-negligible (green) Cliff’s delta effect sizes. **(D)** Summary of peak-to-gene linkage analysis performed using scMultiome data (scATAC + scRNA-seq) from WT and *Zic2*^*-/-*^ D6 AntNPCs. Differential chromatin accessibility and gene expression changes (*Zic2*^*-/-*^ vs WT) were integrated using Seurat’s LinkPeaks() function as described in the Methods section. The number of gene–enhancer links (N) classified according to the direction (increased or reduced) of the enhancer chromatin accessibility and gene expression changes observed in *Zic2*^*-/-*^ D6 AntNPCs are shown. The gene-enhancer links in which the genes showed reduced expression levels and the enhancers showed reduced chromatin accessibility in *Zic2*^*-/-*^ D6 AntNPCs were subject to additional filters: the enhancers were required to display active chromatin features in WT D6 AntNPC and the genes had to be part of the “ZIC2 top target genes”. This resulted in a curated set of 71 gene–enhancer links (ZIC2-dependent GRN), part of which are shown in the table to the right. **(E)** Heatmaps displaying ZIC2 ChIP-seq, ATAC-seq, and H3K27ac ChIP-seq signals around (+/- 1Kb) the ZIC2 peaks belonging to the ZIC2-dependent GRN defined in (D). The ZIC2 peaks are grouped based on their proximity to CGI (CGI+ or CGI-) and are ranked by the H3K27ac signal intensity in WT D6 cells. ChIP-seq and ATAC-seq signals are shown across the indicated time points (D0, D3, D6) and genotypes (WT and *Zic2*^-/-^). **(F)** Genome browser snapshot showing chromatin accessibility (scATAC-seq) at the *En1* locus in WT (coral) and *Zic2*^-/-^ (turquoise) D6 AntNPCs. The predicted ZIC2-regulated enhancer is highlighted and linked to *En1* based on the Multiome analysis described in (D). **(G)** *En1* expression levels were measured by RT-qPCR in mESC and D6 AntNPC that were either WT, *Zic2*^*-/-*^ (clone #1) or homozygous for a deletion spanning the *En1* enhancer highlighted in (F) (clones #29 and #30). For each cell line, *En1* expression was measured in the following number of replicates: D0 samples - one biological replicate with three technical replicates; D6 samples - two biological replicates with three technical replicates each . Expression values were normalized to two housekeeping genes (*Eef1a1* and *Hprt1*). The statistical significance of the expression differences among cell lines was calculated using two-sided unpaired t-tests with Welch’s correction (* fold-change > 2 and P < 0.0001; NS (not significant): fold-change < 2 or P > 0.05). Error bars represent the standard deviation. **(H)** Boxplots showing gene expression fold-changes in *Zic2*^*+/-*^ vs WT D6 AntNPC (HET) or *Zic2*^*-/-*^ vs WT D6 AntNPC (KO) for four gene groups: (1) all genes (control group), (2) genes downregulated in *Zic2*^*-/-*^ D5/D6 AntNPC (Fig. 5A), (3) genes downregulated in the dTAG 24 h/48 h experiments (i.e. “ZIC2 top target genes”) (B), and (4) genes included in the ZIC2-dependent GRN from panel (D). P-values were calculated using wilcox test. The colored numbers beneath the p-values correspond to negligible (gray) and non-negligible (green) Cliff’s delta effect sizes.

The genes downregulated in the short depletion experiments (48h and/or 24h), which, as stated above, are more likely to represent direct ZIC2 targets, were strongly associated with brain development and patterning (Figure S21 B). These genes, which we refer to as “top direct ZIC2 targets”, included key midbrain, roof plate and dorsal patterning regulators (e.g., *Lmx1a, Lmx1b, En1, Pax2*) as well as several WNT signalling genes (e.g.,*Wnt1, Wnt3a*). To identify the regulatory elements through which ZIC2 was likely to regulate its top target genes, we leveraged the Multiome (scRNA-seq + scATAC-seq) dataset generated in WT and *Zic2*^*-/-*^ D6 AntNPCs. Briefly, significant correlations between changes (KO vs WT) in chromatin accessibility at D6 ZIC2 peaks and the expression of nearby genes were assessed using the *LinkPeaks()* function available in Signac (95) (see Methods). This analysis identified a total of 1701 links (Figure 7 D), each corresponding to a gene-ZIC2 peak pair, where changes in the accessibility of the peak were significantly correlated with changes in the expression of the associated gene. Among the 1701 links, the vast majority (1157) included ZIC2 peaks that lost accessibility in D6 *Zic2*^*-/-*^ cells and were paired with genes downregulated in D6 *Zic2*^*-/-*^ cells (Figure 7 D). Next, the 1157 links indicated above were filtered by only keeping those in which the ZIC2 peaks overlapped with D6 active enhancers (H3K27ac+, ATAC+, H3K4me1+, see Methods) and the genes were included among the “top direct ZIC2 targets” (Figure 7 D). This resulted in a refined set of 71 links containing high-confidence ZIC2-regulated enhancers and their associated target genes, thus representing a high-confidence ZIC2-dependent gene regulatory network (GRN) in AntNPC (Supplementary Data 3). Importantly, visualization of the ZIC2 and histone modification ChIP-seq profiles generated in WT and *Zic2*^*-/-*^ cells throughout the AntNPCs differentiation showed that the enhancers that are part of the ZIC2-dependent GRN (i) were often bound by ZIC2 in both D3 and D6 cells; (ii) lost H3K27ac in *Zic2*^*-/-*^ D6 cells; (iii) often overlapped with CGI (i.e. CpG-rich enhancers) and were linked to key brain patterning genes (e.g: *Lmx1a, Lmx1b, En1*) (See Figure 7 E, Figure S17, Figure S18 and Supplementary Data 3). Overall, by combining the ZIC2 degron cell line with the scMultiome data generated in WT and *Zic2*^*-/-*^ AntNPC, we defined the ZIC2 GRN and demonstrated that ZIC2 is necessary in AntNPC for the proper activation of enhancers controlling the induction of major brain patterning genes.

To directly assess the functional relevance of the ZIC2-bound enhancers that are part of the ZIC2-dependent GRN, we used CRISPR-Cas9 to delete a few representative enhancers linked to *En1, Lmx1a* and *Lmx1b* (Figure 7 F, G and Figures S17, S18, S22, S23, S24). These enhancers were selected because they shared the following features: (i) they are associated with key brain patterning genes (*En1*: midbrain; *Lmx1a*: roof plate; *Lmx1b*: midbrain and roof plate); (ii) they are bound by ZIC2 in D3 and D6 cells (Figures S17 and S18); (iii) they harbor CGIs (i.e. CG-rich enhancers), which is typical of enhancers controlling major developmental genes (32); and (iv) they showed reduced H3K27ac and ATAC-seq signals in *Zic2*^*-/-*^ AntNPCs (Figure S17 and S18). Once we obtained several mESC clonal lines with homozygous deletions of each of the selected enhancers (Figures S22 to S24; note that *Lmx1b* enhancer deletion clone 5 harbors a partial deletion in one allele), we differentiated them into AntNPC and measured the expression of the predicted target genes by RT-qPCR. Chiefly, in all three cases, the enhancer deletions led to a strong reduction in the expression of the corresponding target genes, thus confirming their functional relevance (Figure 7 F,G and Figures S22 E, S23 E, S24 E).

Recent studies have demonstrated that CREs whose activity relies heavily on a specific transcription factor are especially susceptible to alterations in its expression levels (96). Therefore, we decided to finally assess whether the ZIC2-dependent GRN was particularly sensitive to ZIC2 dosage, as this could further highlight the genes representing the primary and direct ZIC2 targets. To this end, we generated *Zic2*^*+/-*^ mESC lines (Figure S25), which were then differentiated into D6 AntNPC, together with WT and *Zic2*^*-/-*^ mESC, and analyzed by RNA-seq (Figure 7 H). Then, the RNA-seq data was used to compare the expression of four different groups of genes: (i) all genes (as a control group); (ii) genes downregulated in D5/6 *Zic2*^*-/-*^ AntNPC according to previous analyses (Figure 5 A); (iii) genes downregulated in the 24h and/or 48h ZIC2-dTAG depletion experiments (Figure 7A-B; i,e “top direct ZIC2 targets”) and (iv) genes included in the ZIC2-dependent GRN (Figure 7 D). Interestingly, whereas no expression differences were observed between WT and *Zic2*^*+/-*^ AntNPC for the second group of genes (i.e. downregulated genes in *Zic2*^*-/-*^ AntNPC), such differences were progressively more pronounced for the third and fourth groups, indicating that the “top direct ZIC2 targets” and, especially, the genes included in the ZIC2-GRN were particularly sensitive to the ZIC2 dosage. This final set of results illustrates how our integrative approach allowed us to refine the primary ZIC2 target genes during neural induction, which we speculate might be particularly affected in HPE cases caused by ZIC2 haploinsufficiency.

## Discussion

Here we show that, during the *in vitro* differentiation of mESCs into neural progenitors, ZIC2 plays a dual and sequential regulatory role, first acting as a promiscuous pioneer and then as a selective lineage-specific activator. Initially, and as mESC exit pluripotency and acquire a primed pluripotent state (i.e. D3), ZIC2 binds to and facilitates the opening of a large set of enhancers, potentially preparing them for their subsequent activation in various cell lineages and/or developmental stages. Then, as neural differentiation proceeds, ZIC2 remains bound to and is necessary for the activation of a subset of enhancers controlling the induction of major brain patterning and WNT signaling genes, which in turn might prevent the premature differentiation of neural progenitors into post-mitotic neurons. Uncovering the dual stage-specific regulatory functions of ZIC2 during neural induction was only possible due to our integrative approach, in which we combined constitutive and inducible loss-of-function approaches with bulk and single-cell genomics. Previous studies on canonical pioneer TFs, such as FOXA, suggest that, during development, these factors can bind inactive CREs in progenitor cells and open up their chromatin. Subsequently and as progenitor cells become exposed to differentiation signals, the initial chromatin opening mediated by the pioneer TFs facilitates the recruitment of other TFs and co-factors, ultimately leading to CRE activation and target gene induction (60, 97). However, and in contrast to what we report here, these previous studies did not address whether, following the initial opening of the chromatin, the binding of the pioneer TF was still required once the CREs and their target genes became active. We posit that future studies on pioneer TFs could benefit from using conditional LOF approaches (e.g., degron cell lines) to untangle the distinct regulatory functions that these proteins might display as differentiation processes unroll (98). On the other hand, it has been shown that some pioneer TF (e.g., OCT4, FOXA) not only open chromatin to facilitate the lineage-specific activation of CREs and target genes, but can also have a complementary function whereby they prevent precocious differentiation and/or the establishment of alternative-lineage gene expression programs through the recruitment of co-repressor complexes to a subset of their binding sites (99, 100)). Our data argue that ZIC2 canalizes pluripotent cells towards AntNPCs and prevents their premature differentiation into neurons by mainly acting as an activator, with the repression of neuronal genes most likely stemming from the ZIC2-dependent induction of dorsal patterning and WNT signalling genes.

Overall, our work refines our understanding of ZIC2 function, identifies a ZIC2-dependent GRN during neural induction and sheds light on the molecular basis of the defects observed in mouse holoprosencephaly models (e.g: roof plate and midbrain midline defects, premature differentiation) (10, 12). Notably, our findings also raise new questions regarding the context-dependent factors that drive the functional switch of ZIC2 during neural differentiation as well as the changes in its binding profiles (e.g., why are only some of the D3 binding sites retained in D6?). In this regard, given the promiscuous multi-lineage pioneering activity that ZIC2 displays in primed pluripotent cells (Figure 6), future studies should also address whether the switch towards a selective enhancer activator that ZIC2 undergoes in AntNPC also occurs in other differentiation trajectories in which ZIC2 remains expressed (e.g., NPC in the dorsal spinal tube, retinal ganglion cells (RGC) in the embryonic retina, etc).

### ZIC2 and ZIC3 redundancy in pluripotent cells

Regarding the role of ZIC2 in pluripotency, our results using a mESC differentiation system indicate that ZIC2 does not play a major function in either maintaining naive pluripotency (D0) or establishing primed pluripotency (D3). Consistent with previous reports (14, 15), ZIC2 preferentially binds to distal regulatory elements, including both active and poised/primed enhancers, but also to many promoter regions. Despite its extensive binding, our constitutive *Zic2*^*-/-*^ and conditional 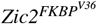 mESC degron lines showed no evident pluripotency or self-renewal defects. Instead, and in agreement with *Zic2 knock-out* mouse models, ZIC2 function seems to be more essential for neural induction and brain patterning. Several non-mutually exclusive explanations may account for the absence of major transcriptional defects in *Zic2*^*-/-*^ mESCs. First, the *in vitro* culture conditions followed in our study do not fully capture the biochemical and biomechanical complexity of pluripotency *in vivo* (101), where ZIC2 might be more functionally relevant. However, and in agreement with our findings, although *Zic2* expression *in vivo* is detected from E0.5 (102), *Zic2* defective mouse models remain unaffected before gastrulation (10, 12). Second, many ZIC2 binding events might be non-functional. This phenomena is common in TF regulatory networks, where the interactions between TFs and chromatin span a continuous range of binding events in functional terms, with only a small subset of the binding events having a functional outcome (103, 104). Third, biochemical, genomic and *in vivo* data (16, 19, 22) together with our own findings strongly suggest that ZIC3 is the most relevant ZIC protein in the pluripotent state, and its expression in our mESC could largely compensate the loss of ZIC2.

### ZIC2 as a promiscuous pioneer TF in primed pluripotent cells

Our findings support a role for ZIC2 as a pioneer transcription factor during the transition from pluripotency to neural induction. In contrast to Luo et al. (2015) (14), who reported that *Zic2 knock-down* mESCs die upon pluripotency exit, our *Zic2*^*-/-*^ cells generated via CRISPR-Cas9 transitioned to the primed pluripotent state without obvious defects. This discrepancy may be explained by technical differences, including the use of shRNA-mediated *Zic2 knockdown* by Luo et al. vs our constitutive *knock-out*, or the possibility of targeting both *Zic2* and *Zic3* when using shRNA, given the high sequence similarity between both genes. On the other hand, based on the ChIP-seq binding profiles of ZIC2 in EpiSC, Matsuda et al. (2007) (15) proposed that ZIC2 could be important for the establishment of the primed pluripotency network. However, using not only ChIP-seq but also RNA-seq and ATAC-seq data (bulk and single-cell) generated in WT and *Zic2*^*-/-*^ D3 cells, we showed that both the silencing of the naive pluripotency program as well as the induction of major primed pluripotency genes ocurred without notable defects in the absence of ZIC2. Instead, chromatin accessibility profiling in *Zic2*^*-/-*^ D3 progenitors revealed that ZIC2 widely promotes accessibility at thousands of distal regulatory elements, consistent with a pioneering function. Our data suggest that ZIC2 might prepare a large number of these enhancers in primed pluripotent cells to facilitate their subsequent activation during differentiation towards multiple cell fates. This role appears evolutionarily conserved, as Opa, the homolog of ZIC proteins in Drosophila, similarly pioneers regulatory regions during early embryogenesis (17, 18). Structurally, it was recently reported that, in hESCs, ZIC2 interacts with BRG1, one of the catalytic subunits of the SWI-SNF complex (16). While this interaction remains to be confirmed in mESCs, it aligns well with our observation that chromatin accessibility defects in *Zic2*^*-/-*^ mESC occur mainly at distal, CGI-poor or orphan CGI regions known to require SWI-SNF activity for chromatin opening (105). In contrast, the ZIC2-bound promoters are typically associated with CGI and remained accessible independently of ZIC2 (106), which is also in agreement with the SWI-SNF independent accessibility of CpG-rich promoters (105). Together, these findings highlight a pioneer role for ZIC2 in primed pluripotent cells, where this TF is necessary for the opening of a large set of enhancers that become subsequently activated during neural induction as well as in other cellular lineages (Figure S16 D). Interestingly, our single-cell analyses indicate that ZIC2 does not control the accessibility of functionally related enhancers (e.g., enhancers active in midbrain progenitors) within specific sub-populations of primed pluripotent cells. Instead, ZIC2 promiscuously controls enhancer accessibility across most pluripotent cells, which, we speculate, could contribute to their broad and largely unrestricted developmental competence.

### ZIC2 as an activator of brain patterning genes upon neural induction

As neural induction progresses, ZIC2 appears to transition from a pioneer TF to a transcriptional activator, directly controlling enhancers involved in establishing region-specific patterning programs (e.g., roof plate, midbrain) and preventing premature neuronal differentiation (e.g., WNT ligand genes). Several *in vivo* studies in mouse and other model organisms are consistent with our findings. With respect to the loss of roof plate identity in *Zic2*^*-/-*^ D6 AntNPCs, it is important to consider that in mouse embryos, *Zic2* expression is restricted to the dorsal part of the developing brain and neural tube from E9.0 to E11.5 (4, 107, 108). Therefore, the loss of roof plate markers (i.e: *Lmxa, Lmx1b, Wnt3a*) in *Zic2*^*-/-*^ AntNPCs points towards a direct and central role of ZIC2 in specifying the roof plate gene regulatory network. A study by Nagai et al. (10) demonstrated that *Zic2*^*Kd/Kd*^ mice, which show an 80% reduction in *Zic2* expression levels, presented HPE together with a wide range of malformations that caused its death soon after birth. Matching our results, these mutants showed a delay in the expression of the roof plate marker *Wnt3a*, which is also downregulated in our *Zic2*^*-/-*^ D6 AntNPCs together with *Lmx1a* and *Lmx1b*, the roof plate master regulators (49, 50, 52, 53). In addition, our work suggests that ZIC2 directly controls the expression of major midbrain regulators (i.e. *En1, Lmx1b, Pax2*). In vertebrates, including mice, ZIC2 expression is notably elevated in the midbrain-hindbrain border region during neural plate development (107, 109, 110). Previous studies have demonstrated that ZIC orthologs in *Drosophila melanogaster* and *Xenopus laevis* can induce the expression of Engrailed (*En*) and some WNT family members (111, 112). The defective expression of these genes leads to notable failures in processes such as cerebellum and caudal midbrain formation during mid/hindbrain development, as evidenced by severe abnormalities in *Wnt1* mutants (77). Collectively, these findings support a role for ZIC2 in defining the midbrain-hindbrain identity *in vivo*. Furthermore, the premature differentiation phenotype seen in *Zic2*^*-/-*^ AntNPCs is consistent with reports linking ZIC2 mutations to size reduction in organs and organisms, a consequence of precocious differentiation of progenitor populations (12, 107, 110, 113, 114). Our data suggest that this effect is likely indirect, resulting from impaired expression of ZIC2-activated genes like *Wnt1* and *Wnt3a*, which normally contribute to the maintenance of progenitor identity via canonical WNT signaling. Together, these *in vivo* studies support our proposed role for ZIC2 as a critical regulator of regional identity and progenitor maintenance during early brain development.

In summary, our integrative approach led to the identification of the key enhancers and genes directly regulated by ZIC2 during the early stages of neural/brain development, as modeled by our *in vitro* differentiation system. Recent work has shown that the regulatory elements whose activity is largely dependent on a given TF are particularly sensitive to changes in its dosage (96). In concordance with this, our experiments using heterozygous *Zic2*^*+/-*^ mESC differentiated into AntNPCs revealed that the most affected genes were part of the ZIC2 GRN that we defined thanks to the combination of our ZIC2 degron cell line with single-cell genomics. Importantly, as ZIC2 happloinsufficiency causes HPE in humans, the identified ZIC2-dependent GRN highlights enhancers and genes that might be particularly vulnerable during neural induction and brain patterning and, thus, potentially implicated in the aetiology of HPE.

### Limitations of the study

This study relies exclusively on a 2D *in vitro* differentiation system, which does not entirely recapitulate the spatial organization and the morphogen gradients that shape regional identities during *in vivo* development (115). While this *in vitro* model allowed us to characterize ZIC2 functionally and mechanistically, it lacks the complexity of more advanced model systems such as 3D organoids (101, 116, 117). Additionally, the use of XAV939 (118) to inhibit WNT signaling from differentiation D2 to D6, while beneficial to inhibit endoderm and mesoderm lineages and to promote anterior identity by reducing cellular heterogeneity, may have partially masked the full functional and molecular relationship between ZIC2 and the WNT signaling pathway. Thus, while our multiomic approach, including scRNA-seq and chromatin accessibility profiling, provided novel mechanistic insights into ZIC2 regulatory function and captured relevant gene regulatory networks downstream of ZIC2, the extent to which these findings reflect *in vivo* biology remains to be further validated in more physiologically relevant systems.

## Methods

### Experimental model details

#### ESC maintenance and differentiation protocol

E14Tg2a (E14) mouse ESC were cultured on gelatin-coated plates using Knock-out DMEM (Life Technologies, 10829018) supplemented with 15% FBS (Life Technologies, 10082147), leukemia inhibitory factor (LIF), antifungal and antibiotics (Sigma-Aldrich, A5955), β-mercaptoethanol (ThermoFisher Scientific, 21985023), Glutamax (ThermoFisher Scientific, 35050038) and MEM NEAA (ThermoFisher Scientific, 11140035). Cells were cultured at 37°C with 5% CO2.

For the NPC differentiation (32), mESCs were plated at 20.000 cells/cm^2^ on geltrex-coated plates (ThermoFisher, A1413302) and grown in N2B27 medium: Advanced Dulbecco’s Modified Eagle Medium F12 (Life Technologies, 21041025) and Neurobasal medium (Life Technologies, 12348017) (1:1), supplemented with 1 × N2 (R&D Systems, AR009), 1 × B27 (Life Technologies, 12587010), 2 mM l-glutamine (Life Technologies, 25030024) and 0.1 mM 2-mercaptoethanol (Life Technologies, 31350010)). During the six days of differentiation, the N2B27 medium was additionally supplemented with the following components: bFGF (ThermoFisher Scientific, PHG0368) 10 ng/ml from day 0 to day 2, Xav939 (Sigma-Aldrich, X3004-5MG) 5 µM from day 2 to day 6, BSA (ThermoFisher Scientific, 15260037) 1 mg/ml at day 0 and 40 µg/mL the remaining days.

### Molecular biology methods

#### Genomic DNA isolation

For routine genotyping of clonal mESC lines, genomic DNA (gDNA) was extracted using a rapid lysis protocol. Cells were resuspended in Cell Lysis Buffer (25 mM KCl (SigmaAldrich, 27810.295), 5 mM TRIS (Sigma-Aldrich, 0497-5KG) pH 8.3, 1.25 mM MgCl2 (VWR BDH7899-1), 0.225% IGEPAL (Sigma-Aldrich, I8896-50 ML) and 0.225% Tween20 (VWR)) supplemented with 0.4 µg/µl Proteinase K (ThermoFisher Scientific, EO0492), vortexed, and incubated sequentially at 65 °C for 6 min and 95 °C for 2 min. Lysates were cooled down, and 1–2 µl were used directly for PCR.

For applications requiring high-purity gDNA, including Sanger sequencing, gDNA was isolated using the NZY Tissue gDNA Isolation Kit (NZYTech, MB13503) according to the manufacturer’s instructions.

#### RNA isolation

Total RNA was isolated with the NZY Total RNA Isolation Kit (NZYTech, MB13402) following the manufacturer’s protocol. For RNA-seq applications, 7.5 µg of RNA were treated with the TURBO DNA-free Kit (Thermo Fisher Scientific, AM1907), and RNA integrity (RIN) was assessed on an Agilent 4200 TapeStation. Only samples with RIN > 8 were used for sequencing.

#### RNA sequencing

Total RNA from D0 (n=3), D3 (n=4), D5 (n=4) and D6 (n=3) WT and *Zic2*^*-/-*^ cells and 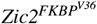 D6 DMSO-treated and dTAG13-treated D6 (n=3) AntNPCs was purified and DNAse-treated as described above. Libraries for WT and *Zic2*^*-/-*^ D3 and D5 samples were prepared using stranded mRNA-seq protocols and sequenced on an Illumina NovaSeq platform at CNAG (Barcelona, Spain) with 50 bp PE reads (>40M reads/sample), while the rest of the samples were processed (i.e. quality control and library preparation) and sequenced by Macrogen Inc. (Korea) using the TruSeq Stranded mRNA protocol (150 bp PE reads). For the analyses performed on the heterozygous *Zic2*^*+/-*^ AntNPCs shown in Figure 7 H, an independent dataset composed by WT (n=3), *Zic2*^*-/-*^ (n=3) and *Zic2*^*+/-*^ (n=3) belonging to an independent differentiation was used. These samples were also processed and sequenced by Macrogen Inc. (Korea) as previously described.

#### cDNA synthesis

For gene expression analysis by RT-qPCR, 1 µg of RNA was reverse-transcribed into cDNA using the ProtoScript II First Strand cDNA Synthesis Kit (New England Biolabs, E6560L) using oligo-dT primers to ensure selective reverse transcription of mRNA.

#### RT-qPCR, ChIP-qPCR and Colony PCR

Reverse transcription quantitative PCR (RT-qPCR) was performed on a Opus qPCR CFX384 detection system (Bio-Rad, 12011452) using the NZYSpeedy qPCR Green Master Mix (2x) (NZYtech, MB224). Reactions were performed in technical triplicates for each sample using the gene-specific primers listed in Supplementary Data 4. The number of biological replicates and independent clonal lines analyzed per condition is detailed in the corresponding figure legends. Relative expression levels of *Lmx1a, Lmx1b* and *En1* were calculated using the 2-ΔΔ CT method, with housekeeping genes *Eef1a1* and *Hprt1* serving as loading controls. In Figures 7, S22, S23 and S24 E, Log_2_-transformed fold changes are presented as box plots, where boxes represent the interquartile range (IQR), the central line indicates the median, and whiskers denote the minimum and maximum values.

ChIP-qPCR experiments for ZIC3 were also performed on the Opus CFX384 detection system (Bio-Rad, 12011452). The binding of ZIC3 to specific genomic target regions (80–120 bp long), including negative control regions (Chr2-neg and Chr6-neg; see Supplementary Data 4) was calculated as percentage of input. Technical triplicates were used for each reaction, and standard deviations were plotted as error bars.

Colony PCR was used to screen *E. coli* transformants for successful insertion of gRNA sequences into the pX330A-hCAS9-long-chimeric-gRNA-G2P (kind gift from Leo Kurian’s laboratory) plasmid. Reactions were performed with NZYTaq II 2x Green Master Mix (NZYTech, MB35803) according to the manufacturer’s protocol. Individual colonies were picked using sterile 10 µl pipette tips and resuspended in 30 µl of sterile deionized water. Cell lysis was carried out by heating 20 µl of the suspension at 95 °C. 3 µl of lysate were used as DNA template. Amplification was performed with a forward primer matching the inserted gRNA sequence and a reverse primer located in the vector backbone (see Supplementary Data 4). A 300 bp amplicon indicated successful integration of the target insert.

#### Genomic deletions using CRISPR-Cas9

Pairs of sgRNAs were designed to flank either coding regions (e.g., *Zic2, Zic3*) or putative enhancers (see Supplementary Data 4), using Benchling’s genome engineering toolbox (https://www.benchling.com/). Guides were selected for high on-target efficiency and minimal off-target potential. Complementary oligonucleotides containing BbsI (NEB, R0539L) overhangs were synthesized and annealed (see Supplementary Data 4) by heating to 95°C for 5 min, followed by gradual cooling down to 25°C at -5 °C/min. Annealed oligos were ligated into BbsI-digested pX330A-hCas9-long-chimeric-gRNA-G2P vector using T4 DNA ligase (NEB, M0202L) and transformed into chemically competent *E. coli* (TOP10) via heat shock. Colonies were plated on LB-ampicillin agar and screened by colony PCR to confirm the presence of the desired insert (see Colony PCR section). Plasmids were purified using the NZY Miniprep Kit (NZYTech, MB01001) and verified by Sanger sequencing with the pX330A_R primer (see Supplementary Data 4). Verified clones were stored as glycerol stocks at -140 °C. mESCs were plated on gelatin-coated 12- or 6-well plates and co-transfected with Lipofectamine 3000 transfection reagent (Thermo Fisher Scientific, L3000001) and one or two sgRNA-expressing pX330A vectors. For 12-well plates, 150.000 mESC were transfected with 1.5 µl of Lipofectamine 3000. Transfection efficiency was assessed via GFP fluorescence 24 h later. Cells with 70–90% transfection efficiency underwent puromycin (Sigma-Aldrich, P8833-25MG) selection (2 µg puromycin/ml during 48 h). Puromycin-resistant cells were serially diluted and plated in 96-well plates for clonal expansion. After 10–12 days, individual colonies were split into duplicate wells: one for gDNA extraction and PCR-based genotyping and another for expansion and maintenance of each clone. Positive clones for the desired genomic edit were expanded and cryopreserved at -140 °C for future use.

### Genomic insertions by CRISPR-Cas9-mediated homology directed repair

(HDR) To generate the DNA double strand break that would favour a genomic insertion through homology directed repair (HDR), a single gRNA was designed in order to cut as close as possible to the insertion site according to the instructions of the genome engineering toolbox from Benchling (https://www.benchling.com/). The steps followed for the design were the same as the ones described for the *Genomic Deletions* section. In order to introduce the desired insertion, a ssDNA repair template was designed and synthesized as an Ultramer™ (for fragments up to 200 bp; IDT) or as a Megamer™ (for fragments up to 500 bp; IDT). The repair template contained the sequence of the epitope (i.e. Flag-HA, FKBP^V36^-HA) flanked with two homology arms, complementary to the sequences upstream and downstream of the gRNA-Cas9 cutting site. mESC were transfected as described above, including the ssDNA sequence as an additional component of the transfection reactions. Subsequent steps to derive mESC clonal lines with the desired genomic insertion were as the ones described for genomic deletions.

### Chromatin Immunoprecipitation coupled to sequencing (ChIP-Seq)

ChIP assays were performed as described in (119) with modifications. For TF ChIPs (ZIC2, ZIC3), one confluent 10 cm plate (∼40–50 × 10^6^ cells) was used; for histone mark ChIPs, 1/5 of a 10 cm plate was used (∼10 × 10^6^ cells). Cells were cross-linked directly on the tissue culture plate in 1% formaldehyde (Sigma-Aldrich, 252549-100ML) for 10 min at room temperature, followed by quenching with 0.125M Glycine (Sigma-Aldrich, G8898-1KG) during 5–10 min. After two washes with 1X PBS supplemented with cOmplete^a^ Protease Inhibitor ULTRA Tablets (Sigma-Aldrich, 5892791001), cell pellets were snap-frozen and stored at –80°C prior to use. Chromatin was extracted by sequential re-suspension in three lysis buffers (Lysis Buffer 1:50 mM Hepes (pH 7.5), 140 mM NaCl, 1 mM EDTA, 10% glycerol, 0.5% NP-40, 0.2% TX-100, dH20 ;Lysis Buffer 2: 10 mM Tris pH 8, 200 mM NaCl, 1 mM EDTA, 0.5 mM EGTA, dH20; Lysis Buffer 3: 10 mM Tris pH 8, 100 mM NaCl, 1 mM EDTA, 0.5 mM EGTA, 0.1% Na-Deoxycholate, 0.5% N-lauroylsarcosine, dH20), followed by centrifugation (5 min, 1350 g, 4 °C) between steps. Chromatin was sonicated using an EpiShear Probe Sonicator (Active Motif, 53052) under the following conditions:

- Histone marks: 30% amplitude, 20 sec on / 30 sec off, 10 min total.
- Transcription factors: 35% amplitude, 20 sec on / 30 sec off, 4 min 40 sec total.

These conditions aimed to obtain chromatin with sizes 200-500bp for histone ChIPs and 500-1000 bp for TF ChIPs. After sonication, samples were cleared by centrifugation (10 min, 16000 g, 4 °C), and supernatants were aliquoted. Each aliquot was incubated overnight at 4 °C with 3–5 µg antibody (for histone marks) or 10 µg (for TFs). A 10% aliquot was reserved as input control. The next day, pre-blocked Dynabeads™ Protein G (Thermo Fisher Scientific, 10004D) were added (100 µl for TFs; 75 µl for histone marks) and incubated for 4 h at 4 °C. Beads were washed five times with RIPA buffer (50 mM Hepes pH 7.5, 500 mM LiCl, 1 mM EDTA, 1% NP-40, 0.7% Na-Deoxycholate, dH2O) and once with TE buffer containing 50 mM NaCl (Sigma-Aldrich, S9888). Chromatin was eluted from the beads by heating at 65°C with occasional vortex. After this, crosslinking was reversed by overnight incubation at 65°C. DNA was purified using DNA Clean & Concentrator kit (Zymo Research, D4034) for inputs and ChIP DNA Clean & Concentrator kit (Zymo Research, D5205) for ChIP samples. Purified DNA concentration was measured with a Qubit 4 Fluorimeter (Thermo Fisher Scientific, Q33238) using the Qubit™ dsDNA HS Assay Kit (Thermo Scientific, Q32854) for ChIP DNA, and the Qubit™ dsDNA BR Assay Kit (Thermo Scientific, Q32853) for input samples. Libraries were prepared for the ChIP and input samples from D0, D3, and D6 by Macrogen Inc. (Korea) using the Illumina TruSeq ChIP-seq protocol. Libraries were sequenced (paired-end 150 bp, 40M reads/sample) on a NovaSeq6000 platform. Please note that for the histone mark ChIP-seq experiments, the chromatin from the mouse cells was mixed with chromatin from human HEK293 cells in a 90%:10% proportion for normalization purposes (120, 121). However, this normalization strategy was discarded due to high variability in the number of human reads obtained for each sample and a quantile normalization strategy was used instead (see details below). The human reads were removed from each sample as described below (see “ChIP-seq data processing” section).

#### Western Blot

Western blot (WB) was used to quantify the expression levels of ZIC2, ZIC3, and loading controls in the different cell lines generated for this project.

Briefly, cells cultured on 10 cm plates were washed with cold 1X PBS (Sigma-Aldrich, D8537-24X500ML) and lysed in 500–1000 µl RIPA buffer supplemented with cOmplete^a^ Protease Inhibitor ULTRA Tablets (Sigma-Aldrich, 5892791001). For differentiating cells (D1–D6), lysis was assisted by sonication using an EpiShear Probe Sonicator (Active Motif, 53052) under the following conditions:

- D1–D2: 50% amplitude, 30 sec on / 30 sec off, 2 min on ice bath.
- D3–D6: 50% amplitude, 30 sec on / 30 sec off, 5 min (in two 2.5 min rounds, with cooling on ice bath).

Whole cell lysates were incubated at 4°C for 20–30 min with rotation and centrifuged (20 min, 13000 rpm, 4°C). Supernatants were collected, and protein concentrations were measured using the Pierce™ BCA Protein Assay Kit (Thermo Scientific, 23209) having a BSA standard curve as reference. Equal amounts of protein extract (30–40 µg) were mixed with 5X Laemmli buffer (0.5M DTT, 10% SDS (w/v), 0.4 M Tris-HCl pH 6.8, 50% Glycerol, 8.2 mM Bromophenol Blue), denatured at 95°C for 5 min, and resolved on 12% SDS-PAGE gels. Electrophoresis was performed using a Mini-PROTEAN system (Bio-Rad) at 90 V for 30 min, followed by 120 V for 1h. Proteins were transferred to nitrocellulose membranes using the Wide Mini-Sub Cell GT Horizontal Electrophoresis System (Bio-Rad, 1704468) in transfer buffer (4°C; overnight at 110 mA).

Membranes were blocked in Tris-Buffered Saline (20 mM Tris-HCl, 150 mM NaCl, pH 7.4–7.6) with 0.1% Tween-20 (VWR, 437082Q) (TBS-T) buffer containing 4% BSA (NZYTech, MB04602) for 1 h at room temperature. After three 5-min washes in TBS-T, membranes were incubated with primary antibodies (see Supplementary Data 4) diluted in blocking solution (4% BSA in TBS-T) for 1h at room temperature or overnight at 4°C. Following three additional TBS-T washes, membranes were incubated with HRP-conjugated anti-rabbit or anti-mouse secondary antibodies (see Supplementary Data 4) diluted in 2% non-fat milk in TBS-T or 4% BSA for 1 h at room temperature.

Membranes were washed twice for 10 min in TBS-T, and signal was visualized by chemiluminescence using an Enhanced Chemiluminiscence (ECL) detection system (Enhanced Chemiluminiscence Solution 1 (WB): 0.1 mM Tris-HCl pH 8.5, 0.4 mM Coumaric Acid, 2.5 mM, Luminol, dH2O; Enhanced Chemiluminiscence Solution 2 (WB): 0.1 mM Tris-HCl pH 8.5, 0.002% Hydrogen Peroxide, dH2O). Images were acquired using a CCD camera-based imaging system.

#### Immunofluorescence assays WT and Zic2^-/-^

D6 AntNPCs were differentiated on glass coverslips (VWR, MEN-ZCB00150RAC20) in 6-well plates as described in previous section. Cells were washed with PBS (Sigma-Aldrich, D8537-24X500ML), fixed for 10 min in 4% paraformaldehyde (VWR, 43368.9M) in PBS at 4°C, and rinsed three times with PBS (5 min each). Permeabilization was performed with 0.5% Triton X-100 in PBS for 5 min, followed by three PBS washes. Cells were then incubated in blocking solution (4% BSA in PBS) for 30 min at room temperature to reduce nonspecific binding. Primary antibodies (see Supplementary Data 4) were diluted 1:100 in blocking solution and applied to the coverslips, which were incubated overnight at 4°C in a humidified chamber. After three PBS washes (5 min each), cells were incubated for 1 h at room temperature with fluorophore-conjugated secondary antibodies (see Supplementary Data 4) diluted 1:500. Slides were washed again (3× PBS), mounted in DAPI-containing medium, and sealed with clear nail polish. Fluorescence images were acquired using a Zeiss Axio Imager M1 microscope and processed with Fiji/ImageJ software.

#### Assay for Transposase Accessible Chromatin (ATAC-Seq)

Tn5 transposase (produced *in house* at EMBL Protein Expression and Purification Core Facility according to a previously described protocol (122)) was pre-loaded with sequencing adaptors (see Supplementary Data 4) by mixing one volume of Tn5 enzyme with half volumes of annealed R1N:ME and R2N:ME oligos (e.g., 2µl Tn5, 1µl R1N:ME, 1µl R2N:ME). The assembly reaction was incubated at room temperature for 30–60 min. ATAC-seq was performed using 20.000 nuclei per reaction. The transposition mix (containing 16 µl Transposition buffer-38.8 mM Tris-Acetate, 77.6 mM Na-Acetate, 11.8 mM Mg-Acetate, 18.8% DMF 0.12% NP40, 0.005X Protease Inhibitor Cocktail, Nuclease-free H2O, 2 µl Assembled Tn5 + oligos and 2 µl of Nuclei suspension) was incubated at 37°C for 60 min with shaking at 500 rpm. DNA was purified using the MinElute Reaction Cleanup Kit (Qiagen, 28204), eluting in 22 µl of elution buffer. Samples were stored at -20°C prior to library preparation. To enable multiplexed sequencing, each sample (WT and *Zic2*^*-/-*^ at D0, D3, and D6) was uniquely barcoded with non-overlapping index combinations (see Supplementary Data 4). For each sample, 20 µl of tagmented DNA were mixed with 30 µl of amplification mix and preamplified in a Veriti™ thermocycler (Applied Biosystems). To determine the optimal number of cycles, 2.5 µl of the preamplified product were subjected to RT-qPCR using 12.5 µl of qPCR mix. Based on amplification curves, the remaining PCR was completed with the minimal number of additional cycles to avoid overamplification. Amplified libraries underwent double-sided size selection using SPRIselect reagent (Beckman Coulter, B23317) at a 1.4× bead ratio, eluting in 22 µl of Elution Buffer. DNA concentrations were measured using the Qubit™ dsDNA High Sensitivity Assay Kit (Thermo Scientific, Q32854), and fragment size distribution was assessed on a 2100 Bioanalyzer using the High Sensitivity DNA Kit (Agilent, 5067-4626). Barcoded libraries were pooled and sequenced on an Illumina NextSeq2000 platform using paired-end reads and ∼40 million reads per sample.

#### Multiome 10X Genomics protocol (coupled scRNA-Seq and scATAC-Seq)

Nuclei isolation from D3 progenitor cells and D6 AntNPCs was performed by washing cells twice with PBS before detaching them from the tissue culture plate with Stempro™ Accutase™ Cell Dissociation Reagent (Thermo Scientific, A1110501). The Accutase incubation times were adjusted to the cell type (6 min for D3 progenitors and 10 min for D6 AntNPCs). Next, Accutase was inactivated with BSA 0.04% in PBS. Cell clumps were disrupted by gently pipetting up and down the cell suspension in the collection tubes. Cell suspension was then centrifuged for 5 min at 300 rcf. The supernatant was aspirated without disrupting the cell pellet and cells were washed twice in BSA 0.04% in PBS. Then, cells were diluted 1/10 in BSA 0.04% in PBS and counted. Next, 10^6^cells were transferred to a DNA LoBind® 1.5 ml microtube (Eppendorf, 0030108051). Cells were centrifuged again at 300 rcf for 5 min at 4ºC. Next, the supernatant was carefully aspirated and cells were lysed by adding 175 µl of pre-chilled Lysis Buffer (10X Genomics) to the pellet while smoothly pipetting up and down 10 times. After incubation for 4 min on ice, suspension was centrifuged at 500 rcf for 5 min at 4º C. Then, supernatant was completely removed without disrupting the nuclei pellet. Nuclei suspension was cleaned up by adding Wash Buffer (10X Genomics) followed by centrifugation (500 rcf, 5 min at 4ºC). Assuming ∼50% nuclei loss during cell lysis, cells were resuspended in chilled Diluted Nuclei Buffer (10X Genomics) to achieve the desired final nuclei concentration. The rest of the steps of the scATAC-seq and scRNA-seq coupled protocol were performed exactly as indicated in the Chromium Next GEM Single Cell Multiome ATAC + Gene Expression User Guide CG000338 Rev F.

### Computational and statistical analyses

#### CGI data gathering

In order to classify the ZIC2 peaks as CGI+ (overlapping a CGI), or CGI- (not overlapping a CGI), three different sets of CGIs were retrieved and combined to be used as a unified reference CGI dataset. Two of them were obtained attending to biochemical properties: (i) CGIs identified by CXXC affinity purification and deep sequencing (CAP–seq), obtained from (123) and referred to as CAP-CGIs, (ii) nonmethylated islands (NMIs) obtained from (124). The third one corresponds to the classical CGI definition based on sequence composition features (length > 200bp, CG% ≥ 50% and CpG observed to expected ratio > 0.6) which was retrieved from the UCSC table browser (125).

#### Bulk RNA-Seq data processing

Reads quality control and removal of low quality regions and/or adapters was performed with *fastqc* (https://www.bioinformatics.babraham.ac.uk/projects/fastqc/), *MultiQC* (126) and *Trimmomatic* (127). Next, reads were mapped to the mouse reference assembly (mm10) using *Hisat2* (128). Those reads with a mapping quality below 10 were discarded with *Samtools* (129). Subsequently, the number of read counts per gene was calculated with *HTseq− count* (130) taking as reference gencode.vM25.annotation.gtf. Differential expression analyses were done with DESeq2. Prior to performing differential expression analyses, counts per million (CPM) were obtained for each gene using the function cpm() available in the *edgeR* package (131). Only genes minimally expressed in at least one of the compared conditions were considered (CPM > 1 for all replicates). Overall, differential expression analyses were done between:

- WT D0 mESC and *Zic2*^*-/-*^ D0 mESC (three replicates each).
- WT D3 mESC and Zic2^-/-^ D3 mESC (four replicates each).
- WT D5 AntNPCs and Zic2^-/-^ D5 AntNPCs (four replicates each).
- WT D6 AntNPCs and Zic2^-/-^ D6 AntNPCs (three replicates each).
- WT D0 mESC and *Zic3*^*-/Y*^ D0 mESC (three replicates each).
- 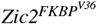 +DMSO D6 and 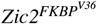 +dTAG-13 D6 (three replicates for each depletion condition i.e. total, 48h, 24h).

The criteria to consider a gene as differentially expressed were:

- Downregulated: adjusted p-value < 0.001 and log_2_FC < *−*1.32.
- Upregulated:adjusted p-value < 0.001 and log_2_FC > 1.32.

In addition, gene expression values in the form of FPKMs were obtained with *Cufflinks* (132).

Regarding the details of the analysis performed with the *Zic2*^*+/-*^ cell-line presented in Figure 7 H, the following steps were applied. First, three different replicates for WT, *Zic2*^*+/-*^ and *Zic2*^*-/-*^ were differentiated to D6 AntNPCs and RNA was extracted at D6. Library preparation and computational pre-processing was performed as described for all samples processed by Macrogen Inc. (Korea). Upon read counts matrix generation, counts per million (cpm) were calculated and the average cpm considering the three replicates was taken as reference. Next, genes genes with an average cpm of 0 in all conditions were discarded. Finally a pseudocount (+1) was added to the remaining gene expression data prior to compute the fold changes described in Figure 7 H.

#### Bulk ATAC-Seq data processing

ATAC-Seq experiments were performed in *Zic2*^*-/-*^ and WT cell lines for three representative differentiation timepoints (D0, D3 and D6). Two biological replicates were analysed in each case. Reads quality control and removal of low quality regions and/or adapters was performed with *fastqc* (https://www.bioinformatics.babraham.ac.uk/projects/fastqc/), *MultiQC* (126) and *Trimmomatic* (127). Next, reads were mapped to the mouse reference assembly (mm10) using Bowtie2 (133). Subsequently, reads with a mapping quality below 10 and duplicated reads were discarded with *Samtools*(129). Differential accessibility analyses between WT and *Zic2*^*-/-*^ cells at each differentiation timepoint were performed with *DiffBind* (134), taking as input the ATAC peaks called for each sample with *MACS2* (135) (narrow peak calling mode and -q 0.05). *DiffBind* was run with default settings. Chromatin accessibility changes were considered significant when the associated adjusted p-value < 0.05. ATAC-seq consensus peaks were also used to predict enhancers (more details in section “Enhancer calling”). Consensus peaks were defined as those regions covered by peaks in the two replicates generated for each sample. Lastly, ATAC-seq BigWigs (genome-wide coverage tacks) were generated applying a quantile normalization between samples of the same differentiation timepoint (more details in section “Quantile normalized BigWigs”).

#### ChIP-Seq data processing

ChIP-Seq data was generated for (i) transcription factors (ZIC3) and (ii) different histone marks (H3K27me3, H3K27ac, H3K4me1, H3K4me2, H3K4me3) in WT and *Zic2*^*-/-*^ cells. In addition, ChIP-seq data for ZIC2 was generated in *Zic2*^*FLAG-HA*^ cells.

For all ChIP-seq samples, reads quality control and removal of low quality regions and/or adapters was performed with fastqc (https://www.bioinformatics.babraham.ac.uk/projects/fastqc/), *MultiQC* (126) and *Trimmomatic* (127).

Regarding the ChIP-seq samples concerning TFs, reads were mapped to the mouse reference assembly (mm10) using *Bowtie2* (133). Subsequently, reads with a mapping quality below 10 and duplicated reads were discarded with *Samtools* (129). Peak calling was performed for each sample with *MACS2* (135) with the narrow peak calling mode. Only those peaks with a q value < 0.01 and fold-change > 4 were considered. When data was available for multiple replicates (i.e ZIC2 ChIPs), consensus peaks were defined as those regions covered by peaks in all replicates. Lastly, those peaks closer than 1 kb were merged. In addition, for each sample, a BigWig (genome-wide coverage track) based on RPKM normalization was generated. Finally, for conditions in which multiple replicates were available (i.e. ZIC2 ChIPs), an average BigWig file was created using *Wiggletools* mean (136) and bedGraphToBigWig (137) tools, to be used as consensus/reference in the different visualizations/quantifications in which BigWig files were used.

Regarding the ChIP-seq samples for histone marks, in order to handle the exogenous DNA (human) added to the samples during their preparation, and following published guidelines (138), reads were mapped to a mm10-hg19 concatenated genome using *Bowtie2* (133). Subsequently, reads with a mapping quality below 10 and duplicated reads were discarded with *Samtools* (129). Next, reads mapping to each of the genomes (mm10 and hg19) were identified using split_bam.py from *Spiker* (https://spiker.readthedocs.io/en/latest/index.html). The bam file with the reads mapping exclusively to mm10 was the only one considered for downstream analyses. ChIP BigWigs (genome-wide coverage tracks) were generated applying a quantile normalization between samples of the same histone mark (e.g., H3K27ac) and differentiation time point (e.g., D3) (for more details see section “Quantile normalized BigWigs”). For calling enhancers (more details in section “Enhancer calling”) in WT cells at various differentiation time points, (peaks for H3K27ac and H3K4me1 were called with *MACS2* (135) with the broad peak calling mode), only peaks with a q value < 0.1 were kept. Next, consensus peaks were defined as those regions covered by peaks in all replicates.

#### Quantile normalized BigWigs

Quantile normalization corresponds with a multi-sample global normalization methodology which transforms data distributions to be identical in their statistical properties (e.g., min, max, median, quartile values) (139). Quantile normalization has been successfully applied to remove technical unwanted variation when processing genomic data, such as, ATAC-Seq, RNA-Seq or ChIP-Seq (139, 140). Nevertheless, it is important to note that quantile normalization assumes that global changes (i.e. a cell line presenting higher levels of H3K27ac genome-wide with respect to another) are not expected due to biological reasons among the normalized samples, because the normalization strategy will eliminate this global pattern. In this regard, quantile normalization between certain samples was applied because global changes due to biological reasons were not expected. This assumption was also supported by the exploration of BigWigs generated with just sequencing depth correction strategies (e.g., RPKM-based BigWigs).

BigWig files for the histone marks (ChIP-seq) and bulk ATAC-seq samples were generated applying a quantile normalization between replicates and samples of the same type and differentiation timepoint (e.g., quantile normalization applied between all H3K27ac D3 ChIP-seq samples, including WT and *Zic2* ^*-/-*^). Firstly, the mm10 genome was split in bins of 100 bps with bedtools makeWindows (141) and the resulting coordinates were converted into saf (simplified annotation format) format, to be accepted as an annotation file in featureCounts (142). Next, for each sample, considering their bam file and the saf file, the number of reads in each of the 100 bp genome windows was counted with featureCounts. The resulting counts matrix was loaded in R where two main steps were applied. In the first one, read counts per bin were converted to RPKMs. Next, the resulting values were quantile normalized among samples using the function normalize.quantiles() from the preprocessCore package https://www.bioconductor.org/packages/release/bioc/html/preprocessCore.html.

Then, bedgraphs with the quantile normalized values per 100 bp window were generated and subsequently converted to BigWigs with the usage of UCSC bedGraphToBigWig tool. Finally, for conditions where multiple replicates were available (e.g., two for H3K27ac WT D3), an average Bigwig file was created using *Wiggletools* mean (136) and bedGraphToBigWig (137) tools.

#### Enhancer calling

Active enhancers were called as described in (39). Briefly, for each differentiation time point (D0, D3 and D6), active enhancers were defined as ATAC-seq peaks overlapping H3K27ac and H3K4me1 peaks. Subsequently, enhancers closer than 5 Kb to TSSs were discarded.

With respect to Poised enhancers, the published dataset in (39) was used, since it consists of a wide set of poised enhancers identified across three different pluripotency states (2i, S+L and EpiLCs).

#### ENCODE enhancers retrieval

Different groups of enhancers from the ENCODE database (93), available through SCREEN portal https://screen.wenglab.org/(143), were downloaded to evaluate the extent of ZIC2 binding to regulatory regions in a wider developmental context.

On the one hand, all proximal and distal enhancers annotated for mm10 were downloaded. This corresponded with the files mm10-cCREs.pELS.bed and mm10-cCREs.dELS.bed respectively. Next, both files were combined, creating the “AllEnhancers” group.

On the other hand, enhancer maps (mm10) for embryonic midbrain and forebrain at 10.5 and 11.5 days were downloaded. This corresponded with the files:

C57BL-6_forebrain_tissue_embryo_10.5_days.enhancers.bed, C57BL-6_forebrain_tissue_embryo_11.5_days.enhancers.bed, C57BL-6_midbrain_tissue_embryo_10.5_days.enhancers.bed, C57BL-6_midbrain_tissue_embryo_11.5_days.enhancers.bed. Next, these 4 files were combined, creating the “Early brain enhancers” group.

#### Transcription factor motif enrichment analyses

TF motif enrichment analyses were performed either with *HOMER* (144) or *MEME-ChIP* (145).

Regarding *HOMER*, the enrichment of known motifs was evaluated in differentially accessible regions (more and less accessible regions in *Zic2*^*-/-*^ were analyzed per separate) with findMotifsGenome.pl (the search for *de novo* motif enrichments was disabled with the option -nomotif).

With respect to MEME-ChIP, motif discovery analysis in ZIC2 and ZIC3 ChIP-Seq peaks was performed using the online version of the MEME Suite 5.5.7 (https://meme-suite.org/meme/doc/meme-chip.html): MEME Suite > Motif Discovery > MEME ChIP. This option is particularly recommended for sequences where the motif sites tend to be centrally located, such as ChIP-Seq peaks. Motif discovery and enrichment mode was run using the classic default mode. Input motifs were “Vertebrates *in vivo* and *in silico*”.

#### Functional enrichment analyses

Functional enrichment analyses were performed either with GREAT (146) or WebGestalt (65).

GREAT was used to perform functional enrichment analyses of the more and less accessible regions identified in *Zic2*^*-/-*^ vs WT cells. First, the BED files containing the more or less differentially accessible regions were loaded into GREAT, setting as background reference genome the mouse GRCm38 (UCSC mm10, Dec. 2011) assembly. For the gene-region associations, the Basal Plus extension default mode was applied. Then, the top GO terms for the “Biological Process” enriched in the resulting gene set were presented as bar plots. WebGestalt (65) was used to perform functional enrichment analyses of differentially expressed gene lists identified by RNA-seq. The analyses were performed upon selecting the following parameters: Over-representation analysis, gene ontology and “Biological Process non redundant”.

#### Heatmaps of aggregated signals

Heatmaps of aggregated signals (e.g., Figure 1 E or Figure 2 H) were generated with computeMatrix and plotHeatmap tools from *deepTools* (147).

#### Cliff Delta effect size estimator

The standardized difference (standardized effect size) for various comparisons was estimated and interpreted using Cliff’s delta (148, 149). It was computed with the cliff.delta() function from the effsize R package (https://cran.r-project.org/web/packages/effsize/index.html). Differences between groups can be interpreted as negligible when | Cliff Delta | < 0.147, small when | Cliff Delta | < 0.33, medium when | Cliff Delta | < 0.474 and large when | Cliff Delta | ≥ 0.474.

#### Single-cell Multiome data processing

Multiome scRNA-Seq and scATAC-Seq derived sequencing files for D3 WT, D3 *Zic2*^*-/-*^, D6 WT and D6 *Zic2*^*-/-*^ were processed independently with the Cell Ranger ARC (2.0.2) software (150). More specifically, cellranger-arc count was employed taking as reference genome the 10X Genomics pre-built version for mm10 (refdata-cellranger-arc-mm10-2020-A-2.0.0). Next, data from WT and *Zic2*^*-/-*^ cells, considering each differentiation timepoint independenly (D3 and D6), were loaded into R with the help of *Seurat* and *Signac* packages (151). Regarding the scATAC-seq data, a unified set of consensus peaks per differentiation time-point was defined by combining the peaks called in WT and *Zic2*^*-/-*^ by Cell Ranger ARC, using reduce() from GenomicRanges followed by the selection of peaks with a width between 20 and 10000 bps and the re-calculation of read counts in this new set of peaks with the help of Signac FeatureMatrix(). Subsequently, only cells meeting the following requirements were kept (i) RNA metrics: 200 < nFeature_RNA < 8000, 500 < nCount_RNA < 50000 and percent.mt < 40 and (ii) ATAC metrics: 200 < nFeature_ATAC < 60000, TSS.enrichment > 1 and nucleosome_signal < 2.

With respect to the scRNA-Seq data, count matrices were normalised with the “LogNormalize” option (available in the Seurat NormalizeData() method) and scaled. Next, PCAs were performed based on the top 2000 most variable features. To mitigate potential batch effects, gene expression data from WT and *Zic2*^*-/-*^ cells were integrated applying Seurat’s Canonical Correlation Analysis (CCA) integration method (152) (available in the IntegrateLayers function). The integration was performed on the PCA embeddings. Then, UMAPs were generated using the first 40 dimensions of the integrated embeddings.

With respect to the scATAC-Seq data, the peak count matrix was normalized through Latent Semantic Indexing (LSI). This involved applying the term frequency-inverse document frequency (TF-IDF) normalization followed by singular value decomposition (SVD) (50 singular values were computed). The first LSI component was not considered in downstream analyses given its expected high correlation with sequencing depth. Next, following an integration strategy analogous to the CCA for scRNA-Seq data, scATAC-Seq data from WT and *Zic2*^*-/-*^ cells was integrated applying Reciprocal Latent Semantic Indexing (RLSI). First, integration anchors between the datasets were calculated with the FindIntegrationAnchors() Seurat function, by specifying reduction =“rlsi” and using dimensions 2-30. Next, a batch-corrected low-dimensional representation was obtained with the IntegrateEmbeddings() function, taking profit of the calculated anchors and the previously computed LSI embeddings. Then, UMAPs were generated using dimensions 2-30 of the integrated LSI embeddings.

#### Pseudotime analyses

Pseudotime analysis was performed with D6 WT and D6 *Zic2*^*-/-*^ scRNA-Seq data. Firstly, cells were classified as *En1*+, *Dcx*+ and *Fabp7*+ if they had at least one read count for the corresponding gene. Next, cells present in more than one group (e.g., *En1*+ and *Dcx*+) were discarded. The Seurat object with the remaining cells was converted to a cell_data_set object thanks to as.cell_data_set() function from SeuratWrappers https://github.com/satijalab/seurat-wrappers. The resulting cell_data_set object was used as input to Monocle 3 (153). Afterwards, cells were clustered with cluster_cells() function, taking as reference dimensionality reduction the one obtained through CCA integration. Next, the principal graph of the analysis was generated with learn_graph() function, setting close_loop to FALSE as no cyclical processes were expected. At this point, taking into account (i) the structure of the principal graph obtained, (ii) the need to choose a node with respect to which the pseudo-times have to be established and (iii) that the selected node can represent either the starting or termination point of the biological process (154), the final pseudotimes were calculated in three steps. In the first step, the center of the *Dcx*+ cell cluster (Figure 5 C) was defined as the root node for pseudotimes calculation (done with order_cells() function). Secondly, the pseudotimes obtained in step 1 were scaled to 0-1 applying a Min-Max normalization. Thirdly, final pseudotimes were calculated by taking their complement (1 - pseudotime obtained in step 2).

#### Single-cell differential expression and differential accessibility analyses

In order to minimize the number of false discoveries (false positives and negatives) and given the absence of true biological replicates, an approach based on pseudoreplication combined with pseudobulking was applied to perform differential expression and differential accessibility analyses (155, 156). First, three pseudoreplicates for WT and *Zic2*^*-/-*^ were created by randomly assigning each cell a group from 1 to 3. The pseudobulk data were then generated by aggregating the information of all cells of the same replicate and genetic background (WT and *Zic2*^*-/-*^). This was done with Seurat AggregateExpression(). Next, differential expression and differential accessibility analyses between WT and *Zic2*^*-/-*^ were performed using FindMarkers() Seurat function, with test.use = “DESeq2”. An adjusted p-value of 0.05 was used as statistical significance cut-off.

#### ZIC2 Gene Regulatory Network (GRN)

A ZIC2-dependent GRN was constructed based on: (i) D6 AntNPCs single-cell Multiome data, (ii) ZIC2 D6 ChIP-Seq peaks, (iii) D6 active enhancer maps and (iv) “top direct ZIC2 targets” (genes downregulated in the short ZIC2 depletion experiments (48h and/or 24h) performed with the 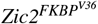 cell line (Figure 7 D)). First, the LinkPeaks() function, available in Signac (95) (defined based on the approach presented in SHARE-seq (59)), was applied to the D6 Multiome data. This function allows identifying significant correlations between changes in the accessibility of ATAC-seq derived peaks and the expression of proximal (<0.5 Mb, default) genes. The execution of LinkPeaks() function returns a set of significant (adj. p-value < 0.05) links, each consisting of an ATAC-seq peak and a matched gene. Only links involving genes differentially expressed and peaks differentially accessible between WT and *Zic2*^*-/-*^ were kept. In addition, links involving peaks that did not overlap with D6 ZIC2 peaks were discarded. After this, the total number of links was 1701. Among those 1701 links, the majority (1157) corresponded to links connecting less accessible scATAC peaks in *Zic2*^*-/-*^ D6 cells and downregulated genes in *Zic2*^*-/-*^ D6 cells (Figure 7 D). Finally, those 1157 links were also filtered to keep only those in which the ATAC peaks overlapped active enhancers in D6 cells and the genes were among the “top direct ZIC2 targets”.

#### Aditional tools

In addition to the tools already mentioned, other softwares were used for different purposes. Figeno (157) was used for the creation of all figures presenting BigWig coverage tracks (e.g., Figure 1 F). In order to share and explore BigWig files among collaborators, the hosting capabilities of Galaxy (158) and the visualization power of the UCSC Genome Browser (159) and IGV (160) were employed.

### Data availability

All the genomic datasets generated in this project have been deposited at GEO. They will be publicly available upon peer-reviewed publication of this work.

## ACKNOWLEDGEMENTS

We would like to thank all members of the Rada-Iglesias laboratory for their insightful comments and suggestions throughout the development of this work. We are grateful to the members of the Kyung-Min Noh and Judith Zaugg laboratories for their valuable input and the sharing of critical resources. We also thank the Protein Expression and Purification Core Facility and Genomics Core Facility teams at EMBL Heidelberg, in particular Laura Villacorta and Vladimir Benes, for their support and guidance with single-cell Multiome data generation. We are especially indebted to Cristina Mayor Ruiz (IRB, Barcelona) for her continuous willingness to help and her expert technical advice on the dTAG-mediated protein degradation system, which was instrumental for the successful establishment of the system.

M.M.F. acknowledges the support of the EMBO Scientific Exchange Grant, which funded her internship at EMBL Heidelberg during which the single-cell multiomic data were generated. M.M.F. PhD was supported by a Grant for Pre-doctoral Contracts for the Training of Researchers – FPI National Subprogram for Training (Ref: PRE2019-089037).

Work in the Rada-Iglesias laboratory is supported by the following grants: PID2021-123030NB-I00 (AR-I), funded by MCIN/AEI/10.13039/501100011033 and by ERDF “A way of making Europe”; RED2022-134100-T (AR-I) (REDEVNEURAL 3.0), funded by MCIN/AEI/10.13039/501100011033; ERC Consolidator Grant Poised-Logic (862022) (AR-I), funded by the European Research Council; ENHPATHY H2020-MSCA-ITN-2019-860002 (AR-I), funded by the European Commission; and Chromrare HORIZON-MSCA-2021-DN-01-101073334 (AR-I), also funded by the European Commission.

## Supplementary Figures

**Fig. S1.**
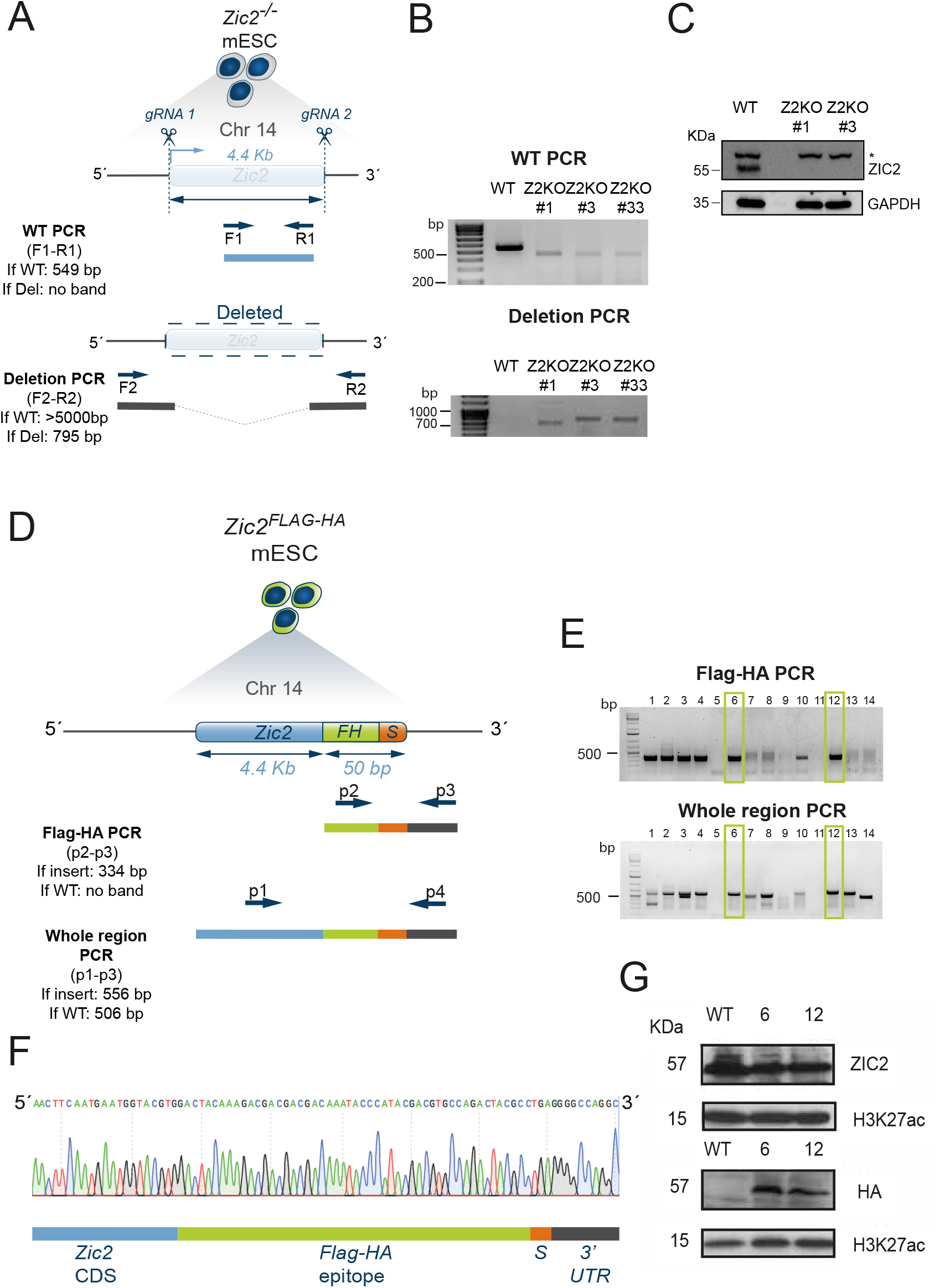
Generation of *Zic2*^*-/-*^ and *Zic2*^*Flag-HA*^ mESCs lines. **(A)** PCR genotyping strategy to identify *Zic2*^*-/-*^ mESC lines. Scissors represent the cut sites of the upstream 5’ and downstream 3’ guide RNAs (gRNAs). Primers used for genotyping are represented as arrows named F1, R1, F2 and R2. **(B)** Agarose gel images showing the genotyping results (WT and Deletion PCR) of three different *Zic2*^*-/-*^ homozygous mESC clones (Z2KO clones #1, #3 and #33). **(C)** Western Blot analysis showing protein expression levels of ZIC2 in WT and *Zic2*^*-/-*^ mESCs clones #1 and #3. (*) ZIC2 Abcam antibody 150404 shows an unspecific band, as reported in (14, 161). **(D)** PCR genotyping strategy to detect mESC clones positive for the endogenous insertion of the FLAG-HA epitope sequence at the C-terminus of ZIC2. Primers used for genotyping are represented as arrows named as p1, p2 and p3. The different PCRs employed (Flag-HA PCR and Whole region PCR) with their PCR products and expected sizes are depicted below. **(E)** Agarose gel images showing the PCR genotyping results for the screened clones. mESC clones #6 and #12, which were positive for the insertion, were selected for further characterization by Western Blot. **(F)** Representative example of the Sanger sequencing chromatogram of one of the selected *Zic2*^*Flag-HA*^ clonal mESC lines (#6) showing that the FLAG-HA epitope sequence is inserted in frame. **(G)** Western Blot analysis of ZIC2 protein expression levels measured with anti-ZIC2 and anti-HA antibodies in WT and *Zic2*^*FLAG-HA*^ mESC lines (clones #6 and #12). H3K27ac is shown as a loading control. Abbreviations: Chr (chromosome); If Del (If deletion); TSS (Transcription Start Site); UTR (Untranslated Region; CDS (Coding Sequence); S (Stop Codon).

**Fig. S2.**
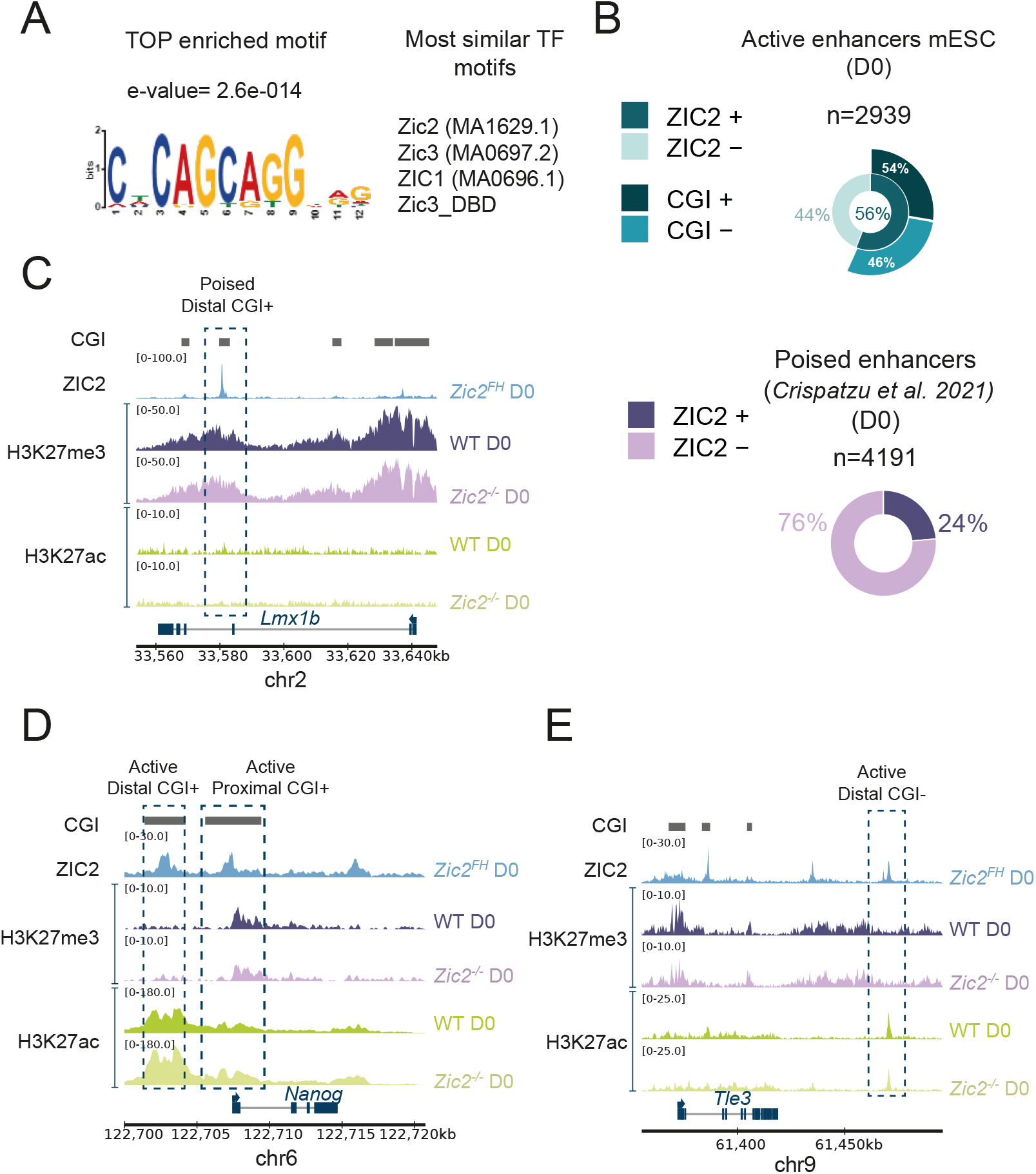
ZIC2 binds to the canonical ZIC motif at both active and poised enhancers in mESCs (D0). **(A)** Top enriched motif identified when performing motif enrichment analysis of D0 ZIC2 ChIP-Seq peaks using MEME-ChIP (145). **(B)** Pie charts summarizing the overlap between D0 ZIC2 ChIP-seq peaks at D0 and either (top) mESCs active enhancers (i.e. ATAC-seq peaks enriched in H3K27ac and H3K4me1) or (bottom) pluripotency-associated poised enhancers as reported by (39). **(C-E)** Genome Browser snapshots showing ZIC2, H3K27me3 and H3K27ac ChIP-seq profiles in WT and *Zic2*^*-/-*^ mESC around (C) a ZIC2-bound poised enhancer located at the *Lmx1b* locus, **(D)** a ZIC2-bound active CGI^+^ enhancer located at the *Nanog* locus and **(E)** a ZIC2-bound distal CGI-active enhancer located at the *Tle3* locus. CGIs are shown as gray bars. ZIC2 ChIP-seq profiles were generated in *Zic2*^*Flag-HA*^ mESCs using an anti-HA antibody.

**Fig. S3.**
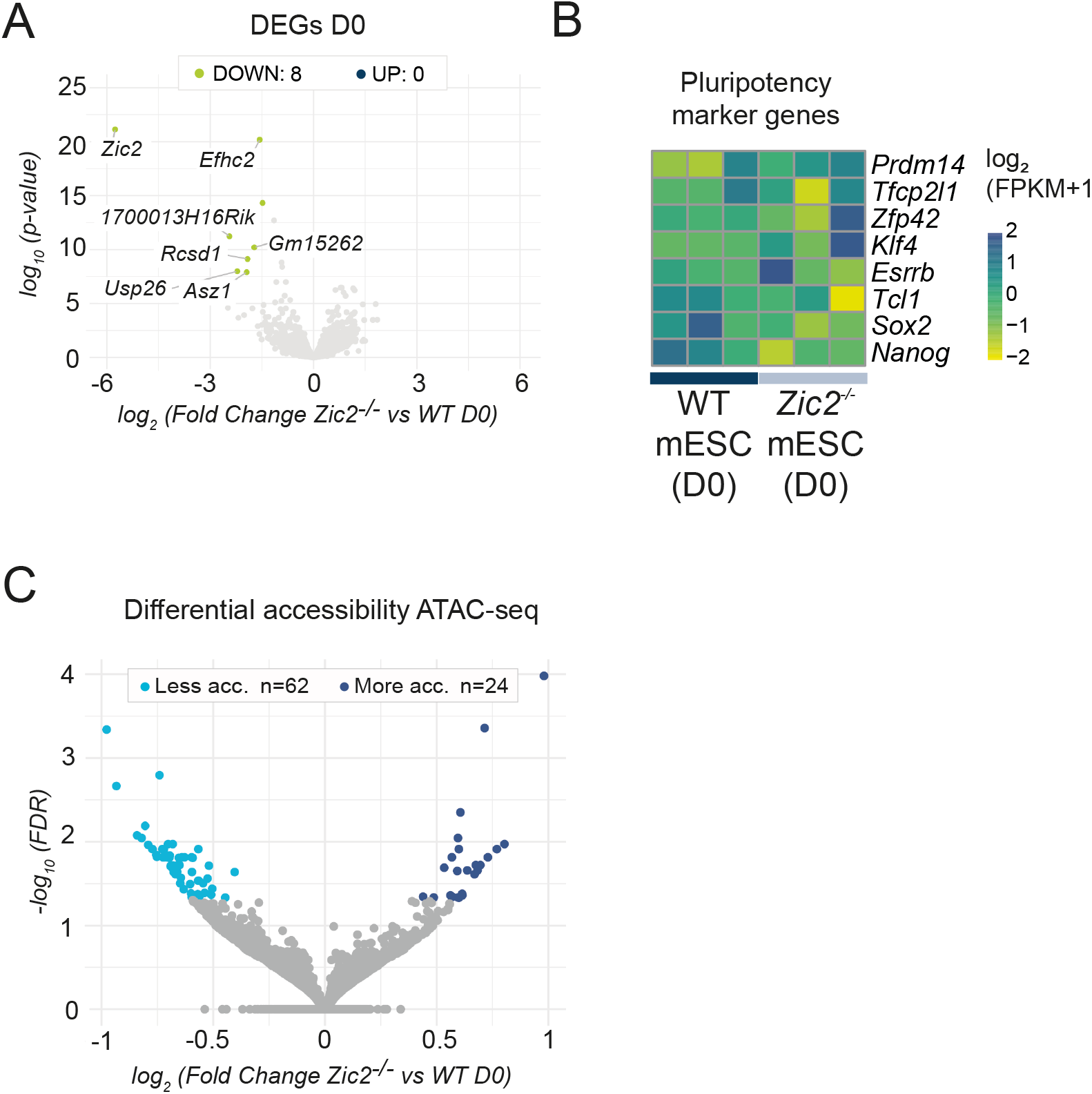
*Zic2*^*-/-*^ mESCs show minimal transcriptional and chromatin accessibility changes. **(A)** Volcano plot of differentially expressed genes (DEGs) in *Zic2*^*-/-*^ versus WT mESC (D0). Significantly upregulated (blue) and downregulated (green) genes in *Zic2*^*-/-*^ mESC are highlighted. **(B)** Heat map showing the expression levels of selected pluripotency genes in *Zic2*^*-/-*^ and WT mESC (D0). The heatmap columns represent biological replicates: WT (three replicates), Z2KO (*Zic2*^*-/-*^): three replicates for clone #1. **(C)** Volcano plot of differential accessible regions between *Zic2*^*-/-*^ and WT mESCs based on ATAC-seq. Genomic regions with significantly decreased (cyan blue) or increased (dark blue) accessibility in *Zic2*^*-/-*^ mESCs are shown.

**Fig. S4.**
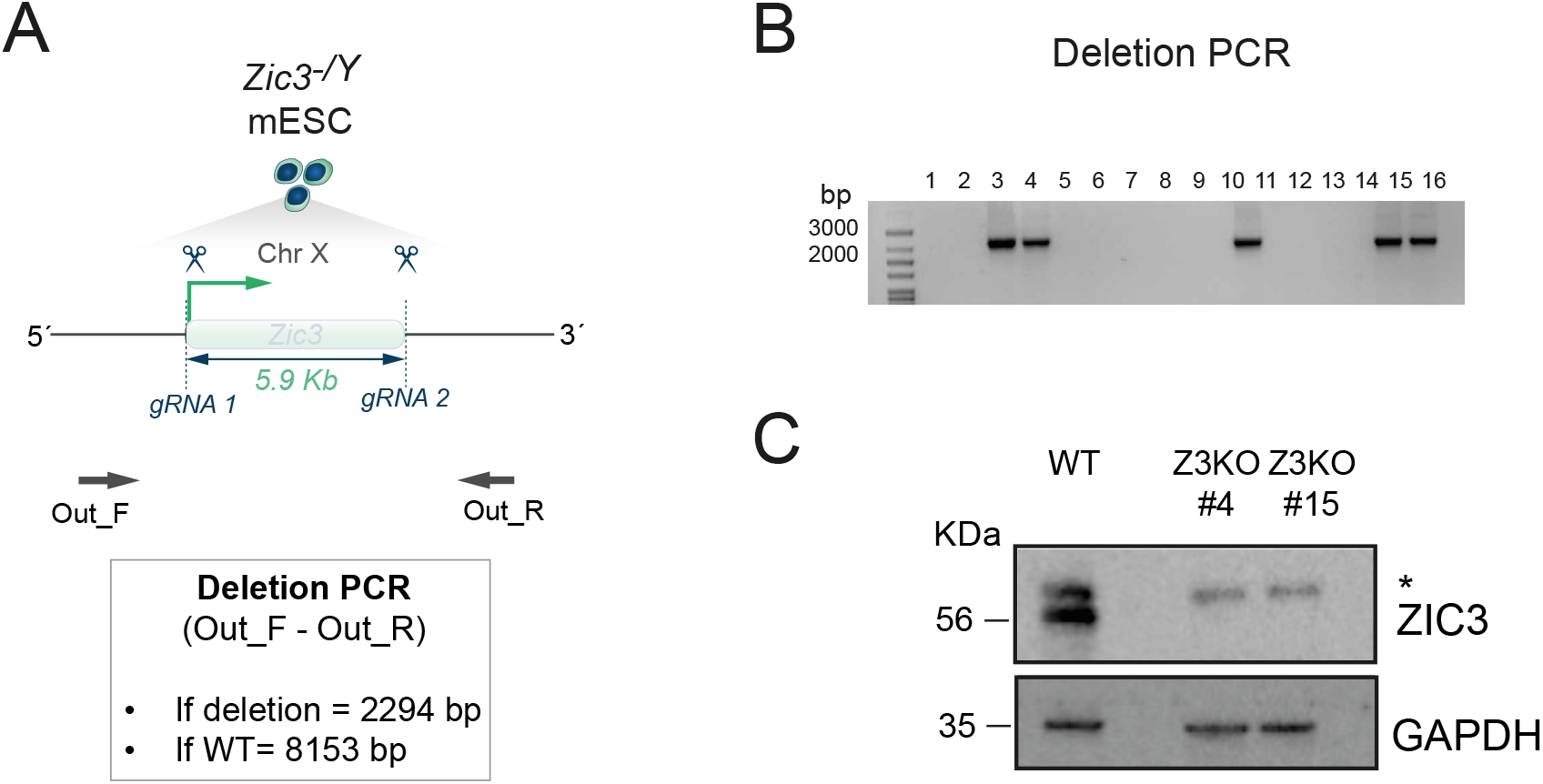
Generation of *Zic3*^*-/Y*^ mESCs. **(A)** PCR genotyping strategy to identify mESC clonal lines with a 5.9 Kb deletion spanning the whole *Zic3* gene. In the diagram, scissors represent the two gRNA cut sites and the arrows represent primers (F2, R2) used to detect the deletion together with the sizes of the expected PCR products. **(B)** Representative example of the PCR genotyping results obtained for several mESC clonal lines, with clones #3, #4, #10, #14 and #15 presenting the *Zic3* deletion. **(C)** Western Blot analysis showing ZIC3 protein expression levels in WT and Zic3^-/Y^ mESCs (clones #4 and #15). (*) ZIC3 Abcam antibody ab222124 shows an unspecific band.

**Fig. S5.**
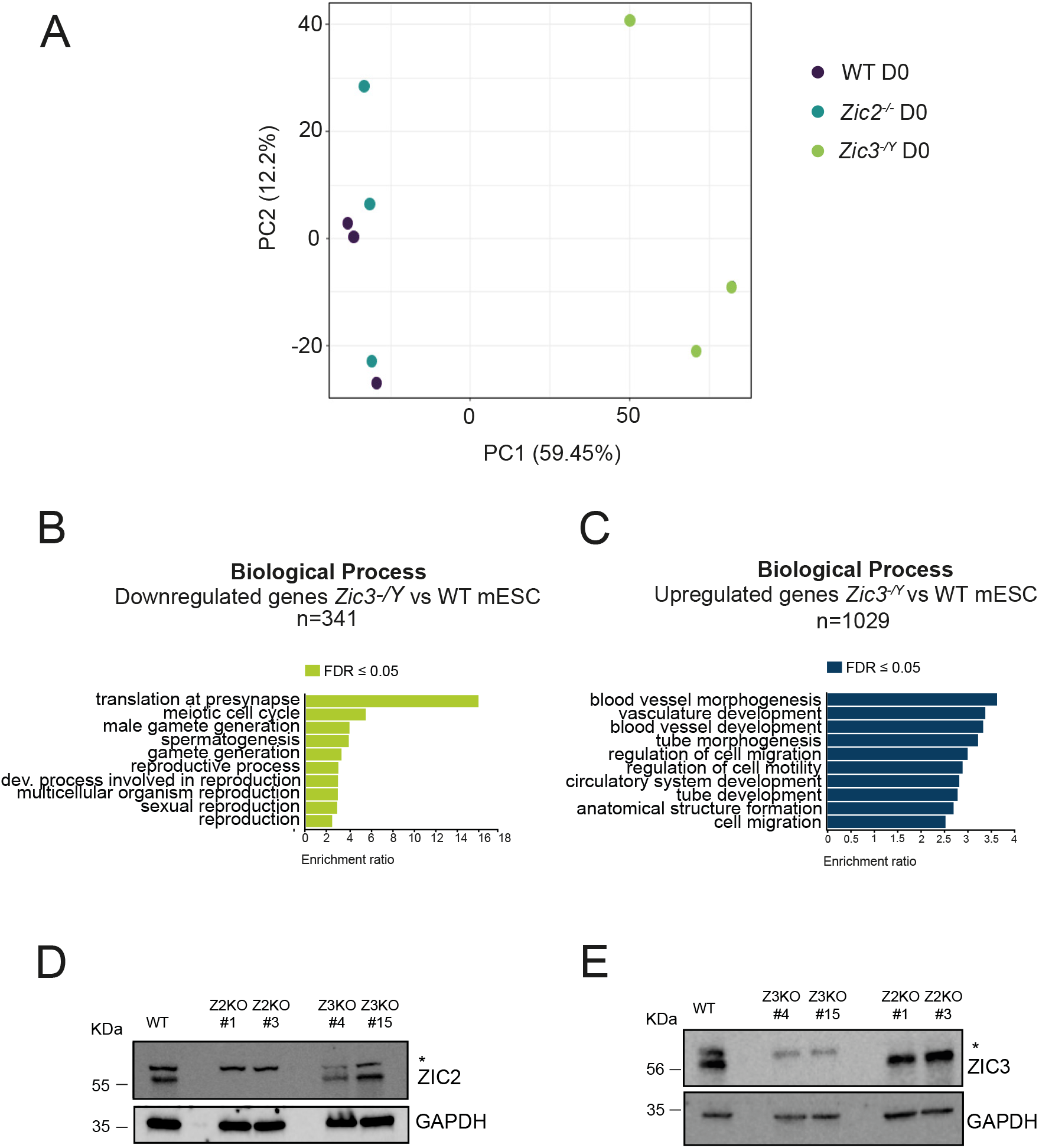
Transcriptomic and protein-level characterization of *Zic2*^*-/-*^ and *Zic3*^*-/Y*^ mESCs. **(A)** Principal Component Analysis of the RNA-Seq samples obtained for WT (three replicates), *Zic2*^*-/-*^ clone #1 (three replicates) and *Zic3*^*-/Y*^ (clones #4, #11, #15; one replicate for each clone) mESC. **(B-C)** Functional enrichment analysis (WebGestalt (65), GO terms for Biological Process non-redundant) of genes that were either (B) downregulated or (C) upregulated in *Zic3*^*-/Y*^ mESC in comparison to WT mESCs. **(D)** Western Blot analysis showing ZIC2 protein levels in WT, *Zic2*^*-/-*^ (Z2KO) and *Zic3*^*-/Y*^ (Z3KO) mESC. GAPDH is shown as a loading control. **(E)** Western Blot analysis showing ZIC3 protein levels in WT, *Zic3*^*-/Y*^ (Z3KO) and *Zic2*^*-/-*^ (Z2KO) mESC. GAPDH is shown as a loading control.

**Fig. S6.**
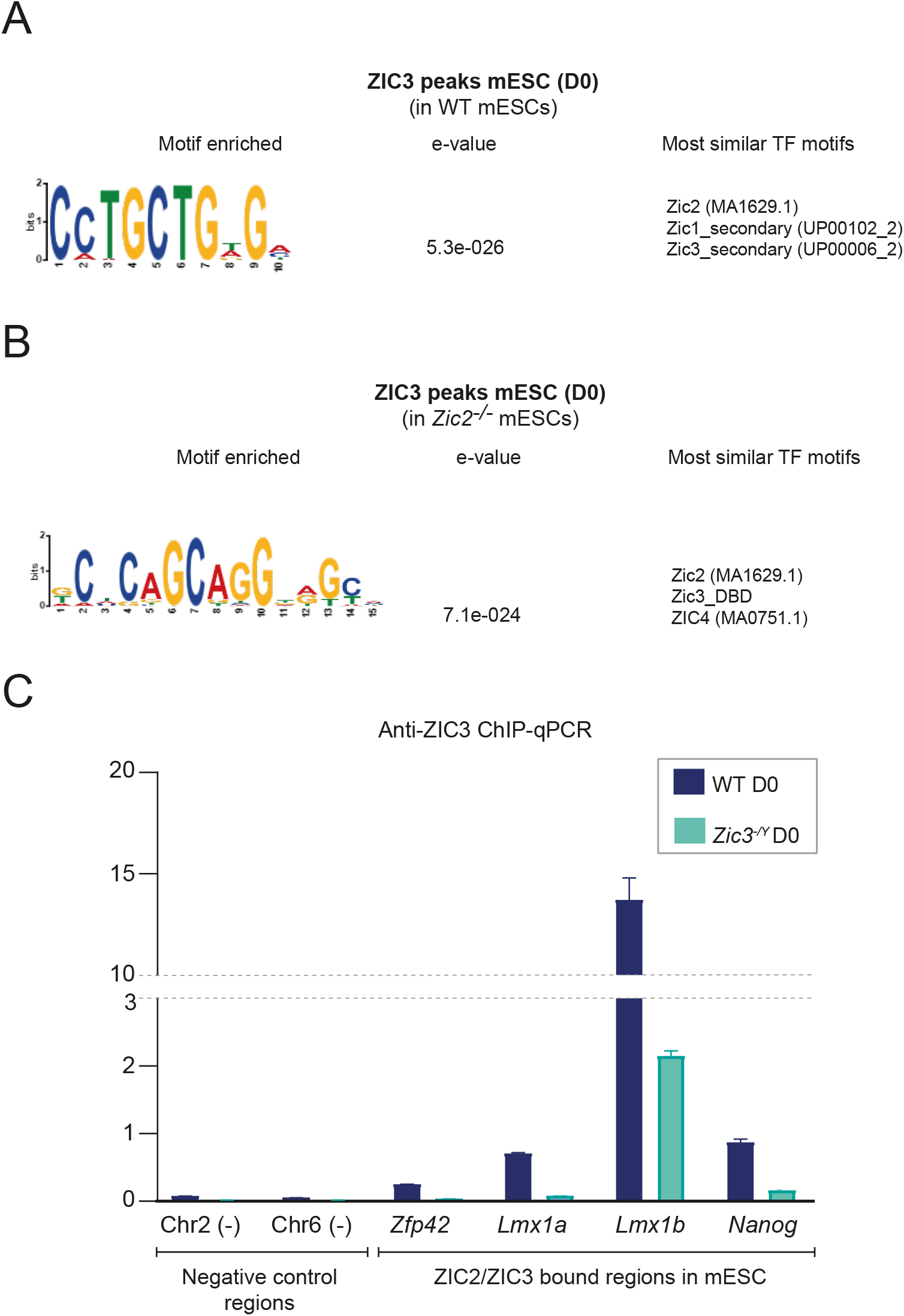
Validation of the anti-ZIC3 antibody (Abcam, ab222124) as suitable for ChIP-seq analysis. **(A-B)** Top enriched motifs identified by MEME-ChIP (145) when analysing the ZIC3 ChIP-Seq peaks identified in (A) WT mESC or (B) *Zic2*^*-/-*^ mESC. **(C)** ZIC3 binding levels were measaured by ChIP-qPCR (as % of input) in WT and *Zic3*^*-/Y*^ mESC at negative control regions (Chr2(-), Chr6(-); Supplementary Data 4) as well as at several regions considered to be bound by both ZIC2 and ZIC3 according to our ChIP-seq data and previous studies (14, 19). The strong reduction in ZIC3 binding levels observed for the positive control regions in *Zic3*^*-/Y*^ mESC supports the specificity of the anti-ZIC3 antibody (ab222124).

**Fig. S7.**
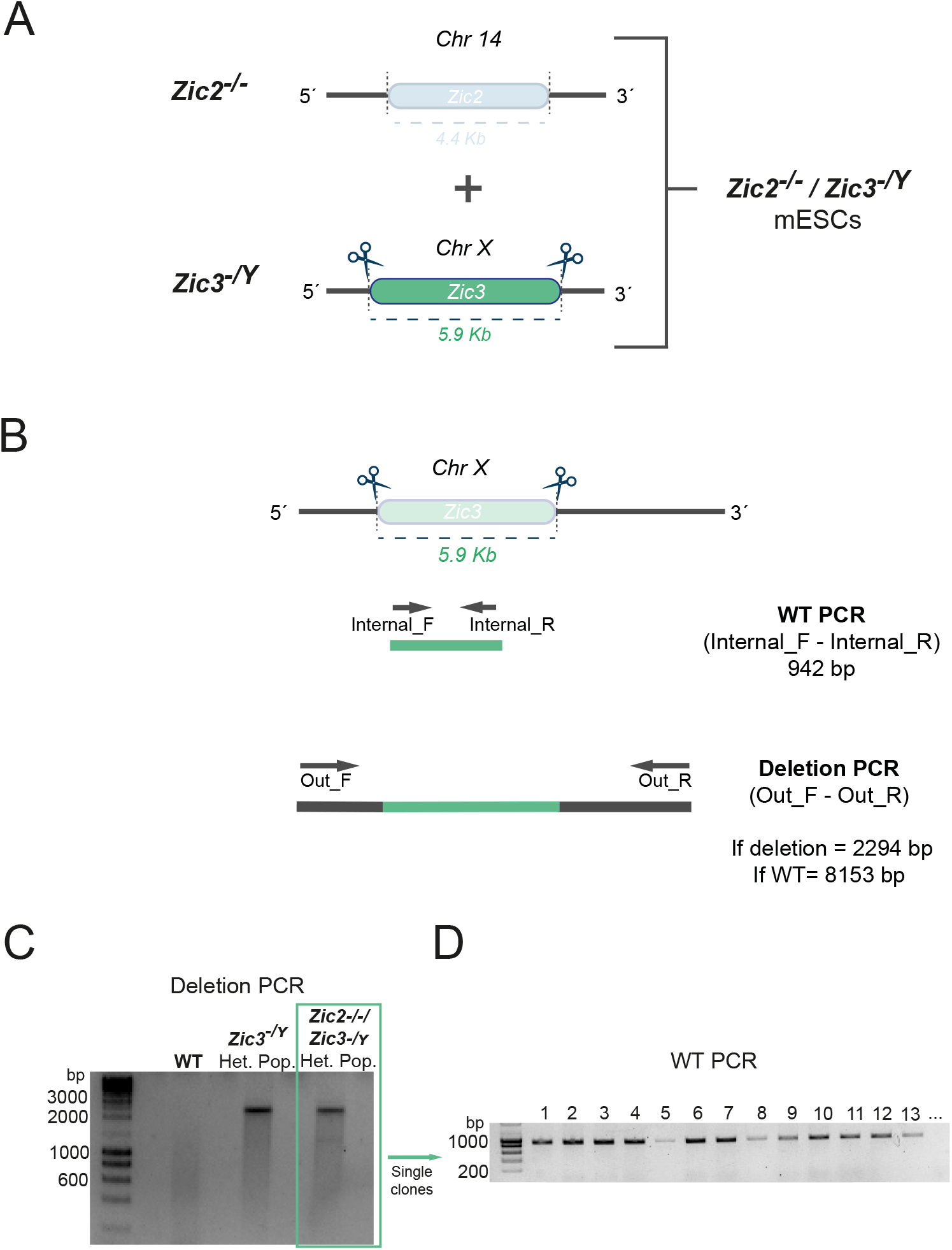
Attempted generation of *Zic2*^*-/-*^*/Zic3*^*-/Y*^ mESCs by CRISPR-Cas9. **(A)** Schematic diagram depicting the strategy followed to generate *Zic2*^*-/-*^/*Zic3*^*-/Y*^ mESC lines. Scissors represent the cut sites of the two gRNAs employed to generate the deletion by CRISPR-Cas9 in a *Zic2*^*-/-*^ clone as parental cell line. **(B)** PCR genotyping strategy to isolate positive clones for the *Zic3* deletion in *Zic2*^*-/-*^ mESCs. In the diagram, the scissors represent the two gRNA cut sites, arrows represent the primers (named Internal F, Internal R, Out F and Out R) used in the two genotyping PCRs showed below (WT PCR and Deletion PCR) together with the sizes of the expected PCR products.**(C)** PCR genotyping experiments performed in untransfected WT mESC, heterogeneous population (Het. Pop.) of WT mESC transfected with gRNAs designed to delete *Zic3* (*Zic3*^*-/Y*^ Het. Pop.) and heterogeneous population (Het. Pop.) of *Zic2*^*-/-*^ mESCs transfected with gRNAs designed to delete *Zic3* (*Zic2*^*-/-*^ *Zic3*^*-/Y*^ Het. Pop.). **(D)** Representative example of the WT PCRs performed for clonal mESC lines isolated from the *Zic2*^*-/-*^ *Zic3*^*-/Y*^ heterogenous population of transfected mESC shown in (C). The presence of WT products indicates that the *Zic3* deletion is not present in any of the analyzed clonal mESC lines.

**Fig. S8.**
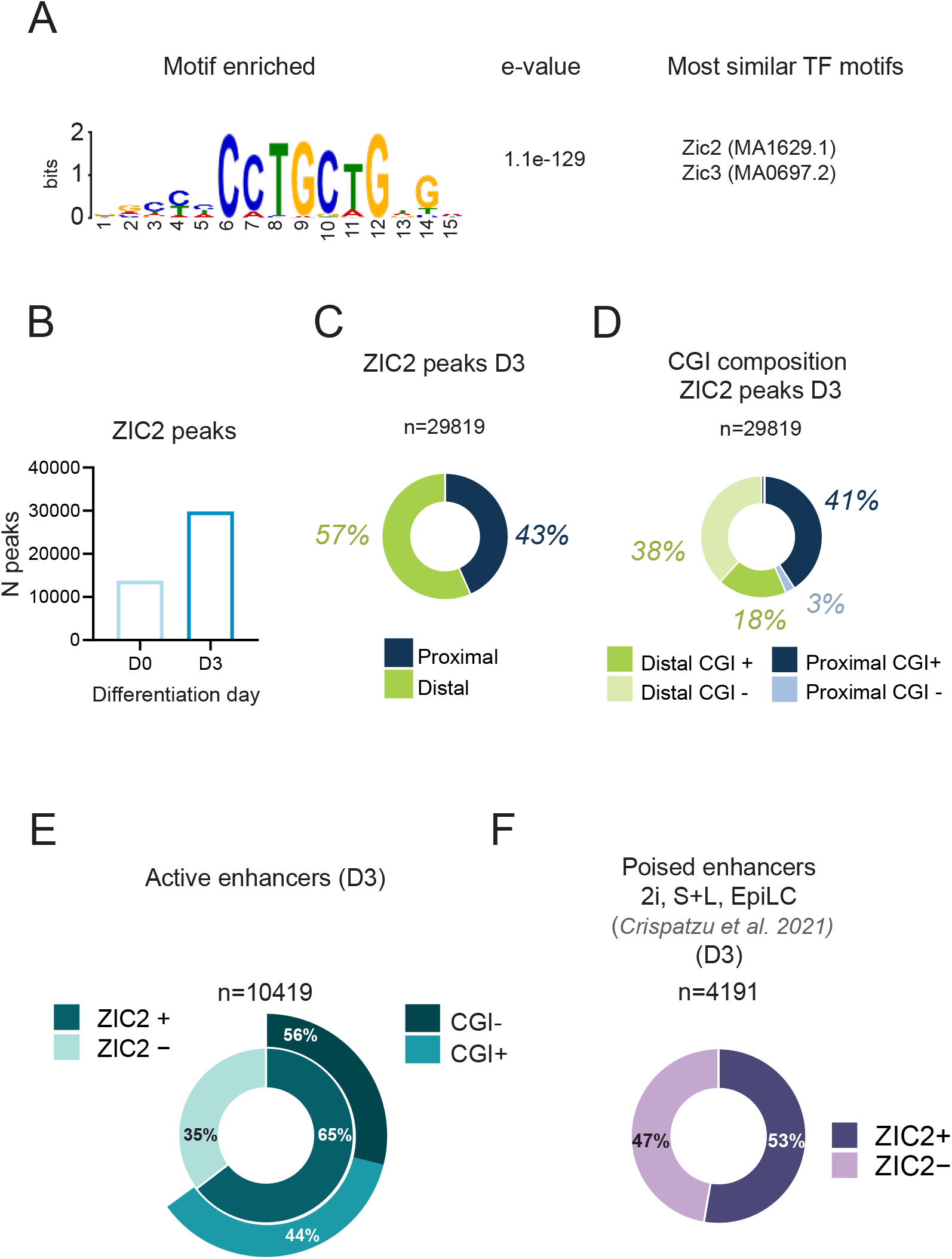
Genomic and epigenetic features of the ZIC2 peaks identified in D3 cells. **(A)** Top enriched motif identified by MEME-ChIP (145) when analysing the ZIC2 ChIP-Seq peaks identified D3 in WT cells. **(B)** Bar plot indicating the total number (N) of ZIC2 ChIP-seq peaks identified in D0 and D3 WT cells. **(C)** Pie chart indicating the percentage of D3 ZIC2 peaks that are either proximal (<5 Kb from a TSS) or distal (>5kb from a TSS) to a TSS. **(D)** Pie chart subdividing the proximal and distal ZIC2 peaks shown in (C) according to their distance to CGIs: CGI+: ZIC2 peaks <3 Kb from a CGI; CGI-: ZIC2 peaks >3 Kb from a CGI (see Methods). **(E)** Pie chart representing the percentage of D3 active enhancers (n=10419) bound by ZIC2 in D3 cells. D3 active enhancers (ATAC-seq peak enriched in H3K27ac and H3K4me1) were called using the ChIP-Seq and ATAC-Seq datasets generated in WT D3 cells (see Methods). **(F)** Pie chart representing the percentage of pluripotency-associated poised enhancers reported by (39) (n= 4191) that are bound by ZIC2 in D3 cells.

**Fig. S9.**
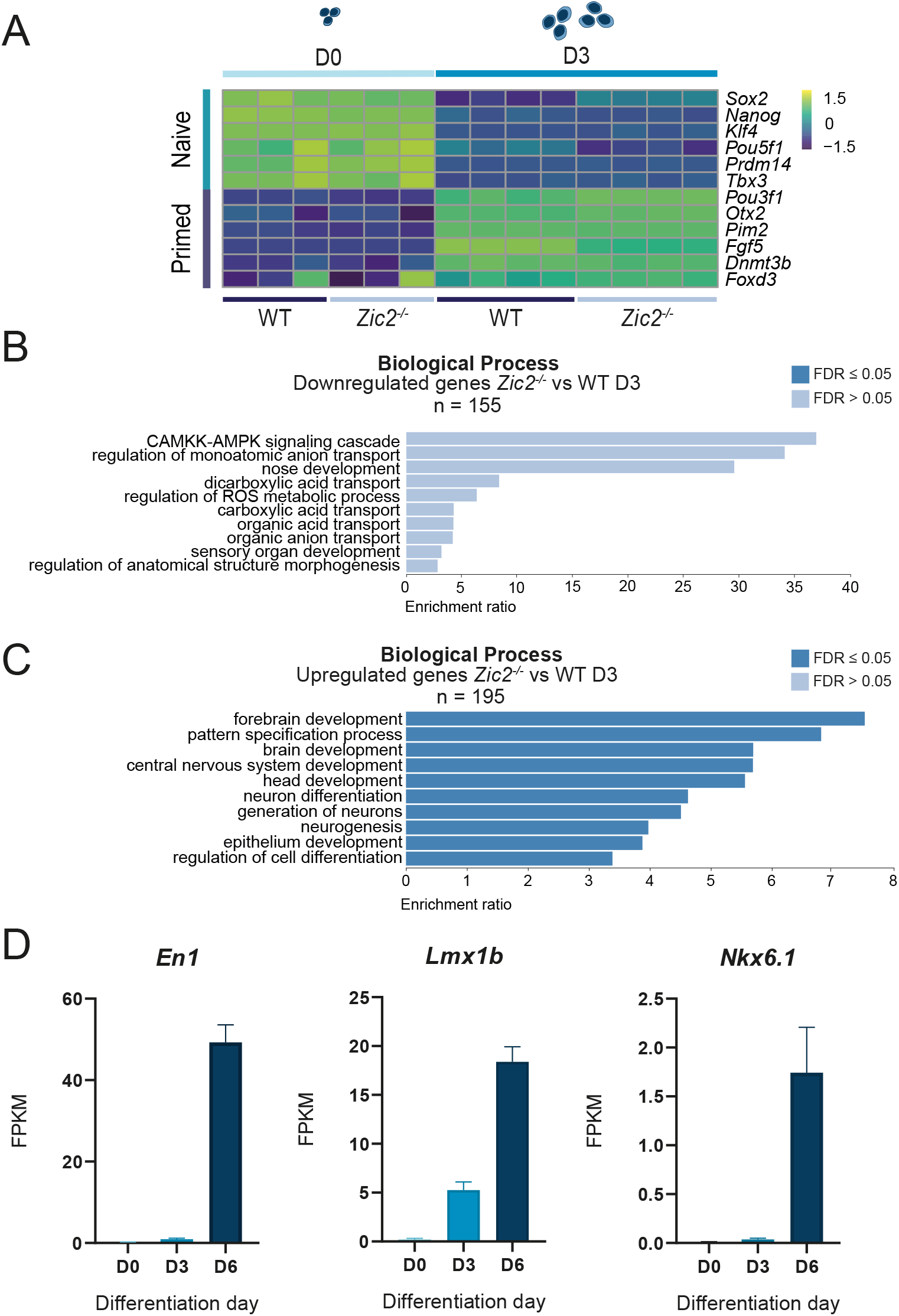
Transcriptional changes in D3 *Zic2*^*-/-*^ cells. **(A)** Heat map showing the temporal expression dynamics (as measured by bulk RNA-Seq data) of selected naive and primed pluripotency markers in WT and *Zic2*^*-/-*^ (KO) cells at both D0 and D3. Heatmap columns represent biological replicates. Scale expression units: log_2_ (FPKM+1). **(B-C)** Functional enrichment analysis results (WebGestalt (65), GO terms for Biological Process non-redundant) for the genes that are either (B) downregulated or (C) upregulated in D3 *Zic2*^*-/-*^ cells in comparison to D3 WT cells. **(D)** Bar plots showing the expression levels (as measured by RNA-seq; FPKM) of representative brain patterning genes in D0, D3 and D6 WT cells. Values represent the average of three biological replicates; error bars indicate the standard deviation.

**Fig. S10.**
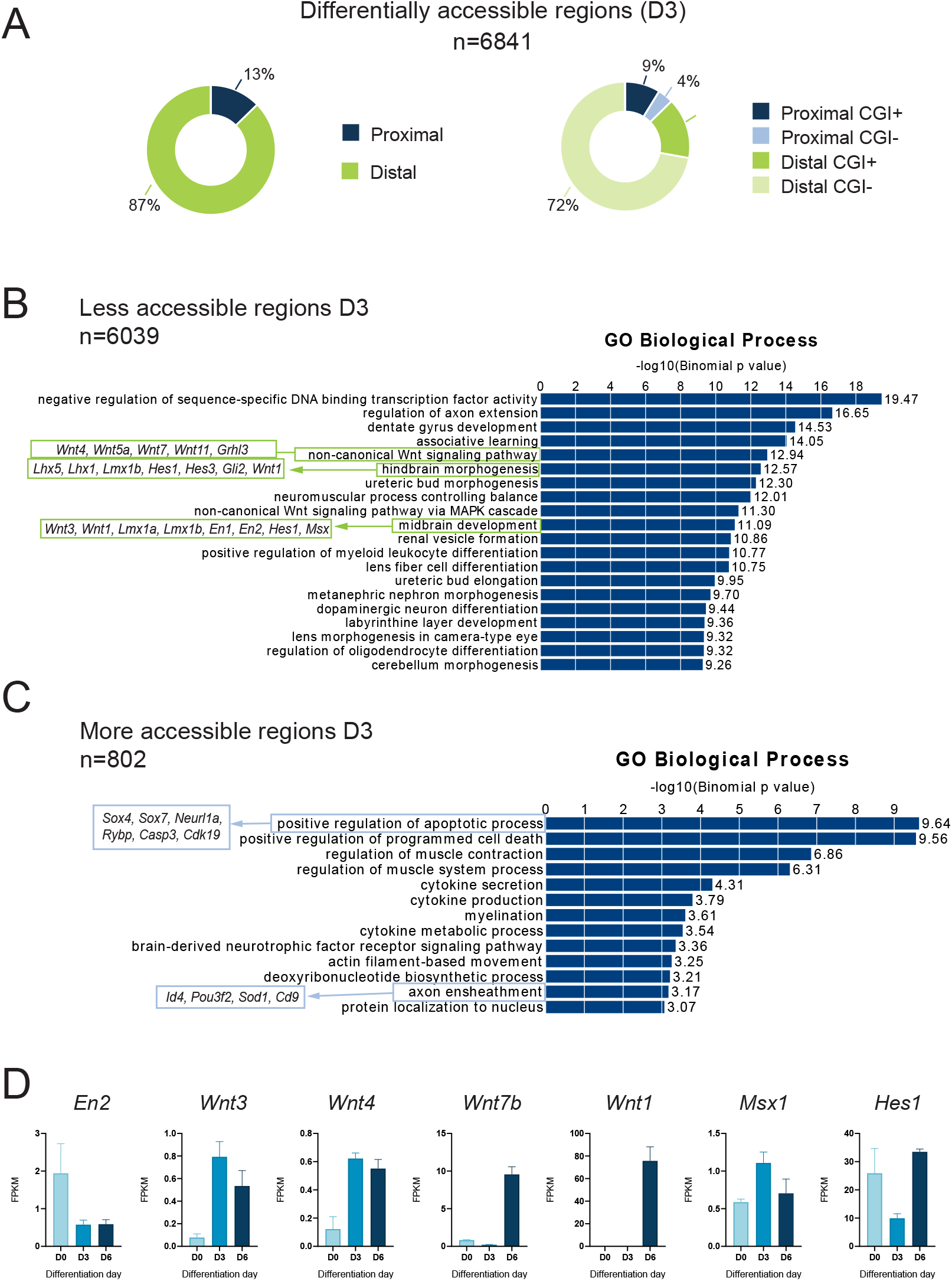
Genomic and transcriptional features associated with the regions displaying chromatin accessibility changes in D3 *Zic2*^*-/-*^ cells. **(A)** Pie charts showing the classification of the differentially accessible regions identified in D3 cells according to the ATAC-seq data generated in *Zic2*^*-/-*^ and WT D3 cells. The differentially accessible regions are classified based on distance to TSS (left) and CGI (right) as described in Figure 1 D. **(B)** Functional enrichment analysis (performed using GREAT (146)) results showing the main GO Biological processes enriched in regions that are either less **(B)** or more **(C)** accessible in *Zic2*^*-/-*^ D3 cells in comparison to WT D3 cells. Green and blue boxes highlight some specific GO terms and include some of the responsible genes for the enrichment of those terms. **(D)** Bar plots showing the expression levels (as measured by RNA-seq, FPKM) of a subset of genes (i.e. *En2, Wnt3, Wnt4, Wnt7b, Wnt1, Msx1, Hes1*) associated with regions losing chromatin accessibility in *Zic2*^*-/-*^ D3 cells. Expression levels are shown for D0, D3 and D6 WT cells. Bars represent the mean of three biological replicates; error bars indicate the standard deviation.

**Fig. S11.**
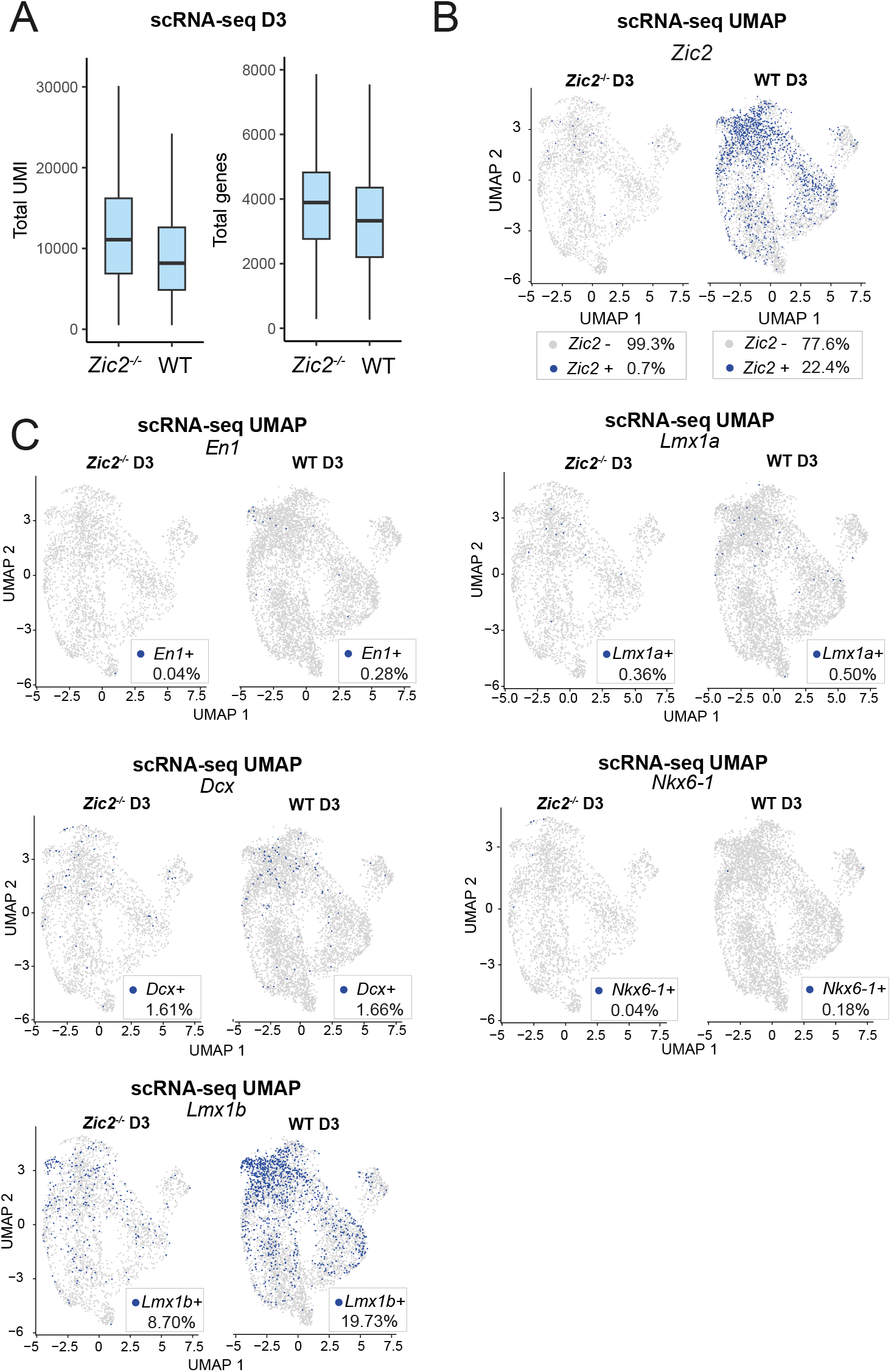
Single-cell RNA-seq analysis of WT and *Zic2*^-/-^ D3 cells. **(A)** Boxplots showing the distribution of total UMI counts (left) and number of detected genes (right) per cell in WT and *Zic2*^*-/-*^ D3 samples after quality filtering. **(B)** UMAP plots of the scRNA-seq generated in WT (right) and *Zic2*^*-/-*^ (left) D3 cells. Cells are labelled according the presence (blue, if ≥ 1 read count) or absence (gray) of *Zic2* expression . The percentage of ZIC2-positive (*Zic2 +*) and ZIC2-negative (*Zic2 -*) cells for each genotype is also indicated. **(C)** UMAP plots showing the expression of neural patterning (*En1, Lmx1a, Lmx1b, Nkx6-1*) and neuronal differentiation (*Dcx*) genes in WT and *Zic2*^*-/-*^ D3 cells. Cells with detectable expression (*≥* 1 read count) of each gene are labelled in blue. The percentage of WT and *Zic2*^*-/-*^ D3 cells positive for each gene is shown in the bottom right corner of each panel.

**Fig. S12.**
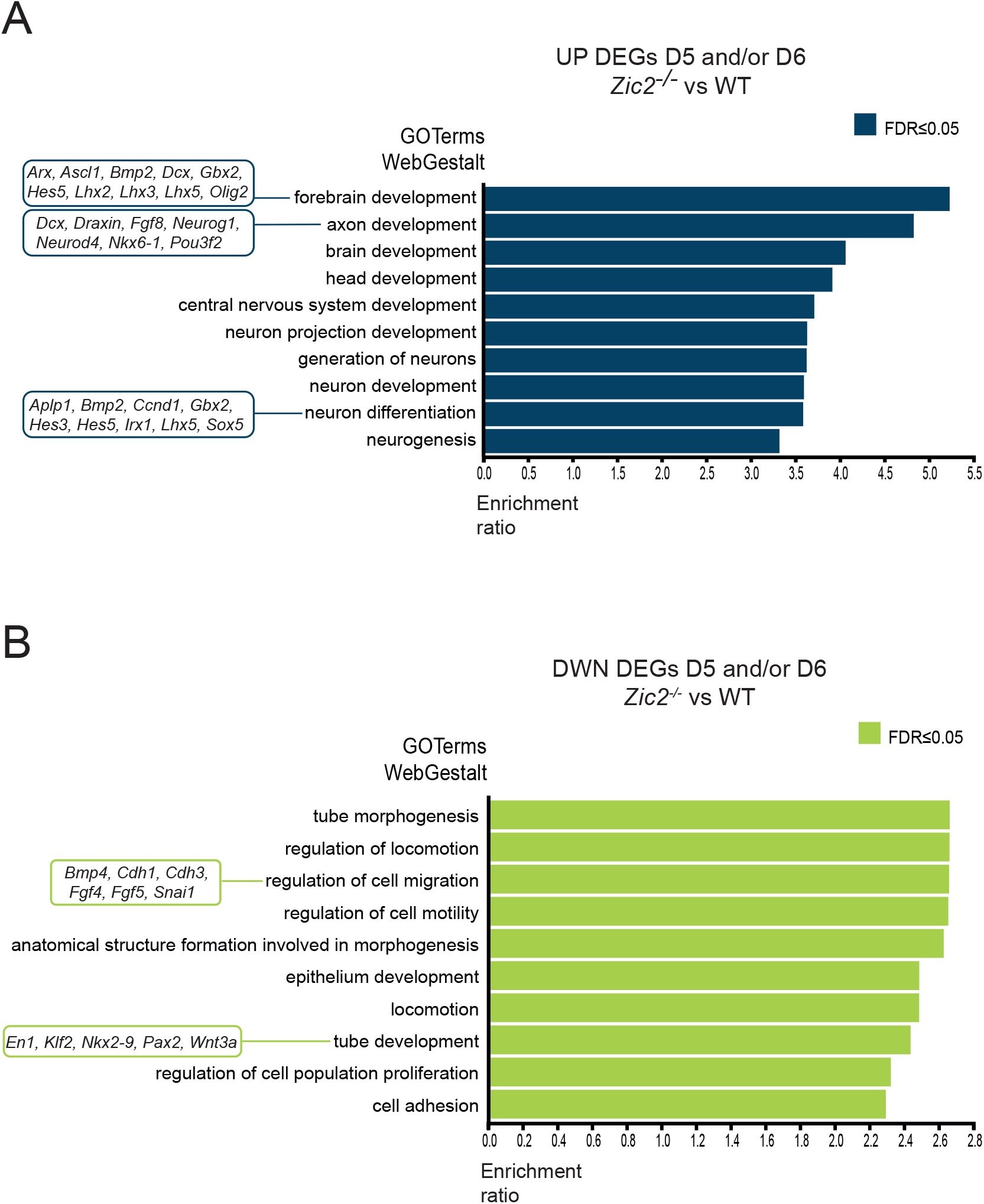
GO enrichment analysis of differentially expressed genes in D5/D6 *Zic2*^*-/-*^ AntNPCs. **(A-B)** Functional enrichment analysis (WebGestalt (65), GO terms for Biological Process non-redundant) of genes that were either (A) upregulated (UP) or (B) downregulated (DWN) in D5/6 *Zic2*^*-/-*^ AntNPCs in comparison to D5/6 WT AntNPCs. Representative genes associated with selected GO terms are indicated in boxes.

**Fig. S13.**
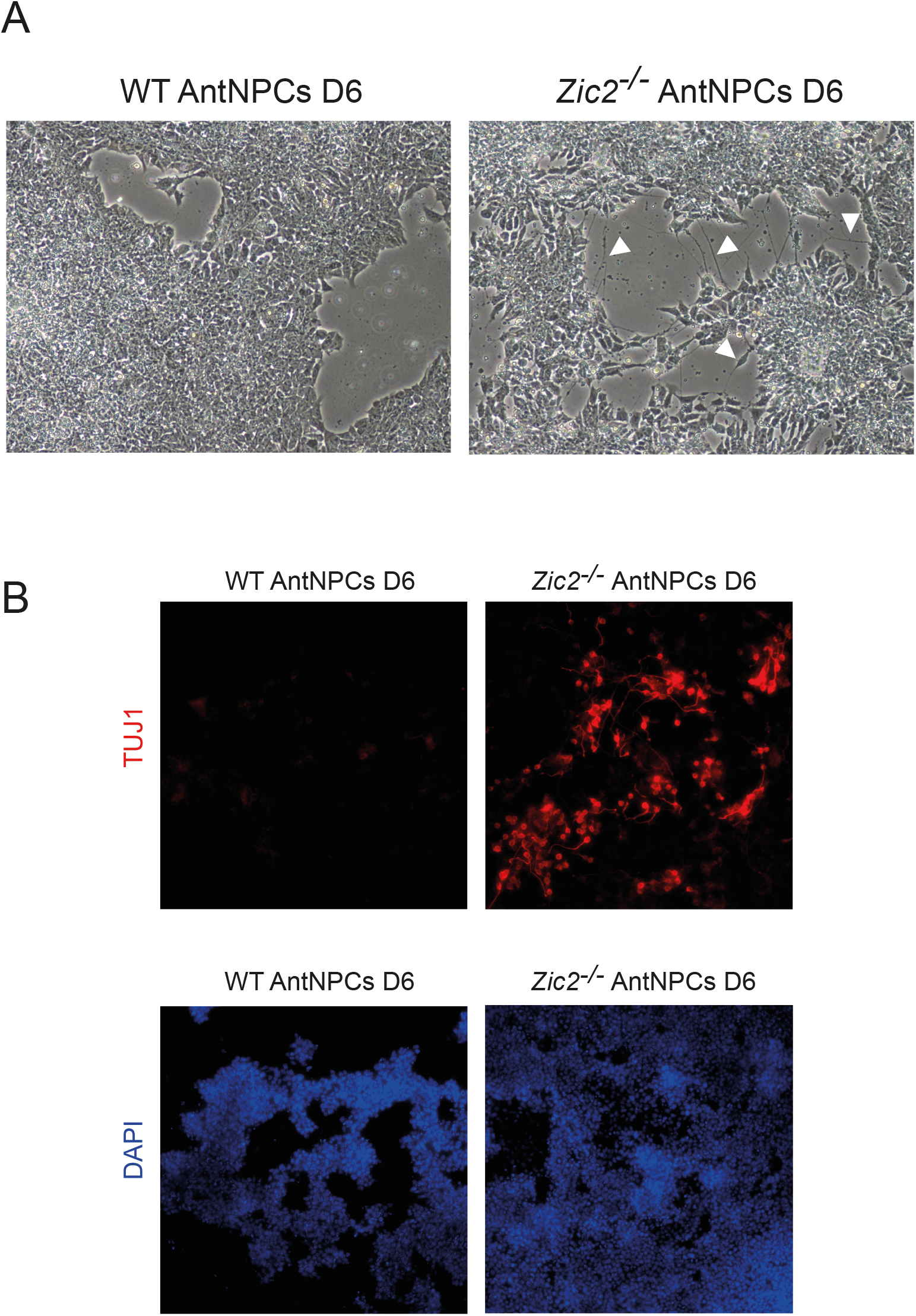
Premature neuronal differentiation of *Zic2*^*-/-*^ D6 AntNPCs. **(A)** Brightfield microscopy images (10X) of WT and *Zic2*^*-/-*^ D6 AntNPCs. White arrows highlight neurites present only in *Zic2*^*-/-*^ AntNPCs. **(B)** Representative images from immunofluorescence staining of TUJ1 protein (up) and DAPI staining (down) in WT and *Zic2*^*-/-*^ D6 AntNPCs.

**Fig. S14.**
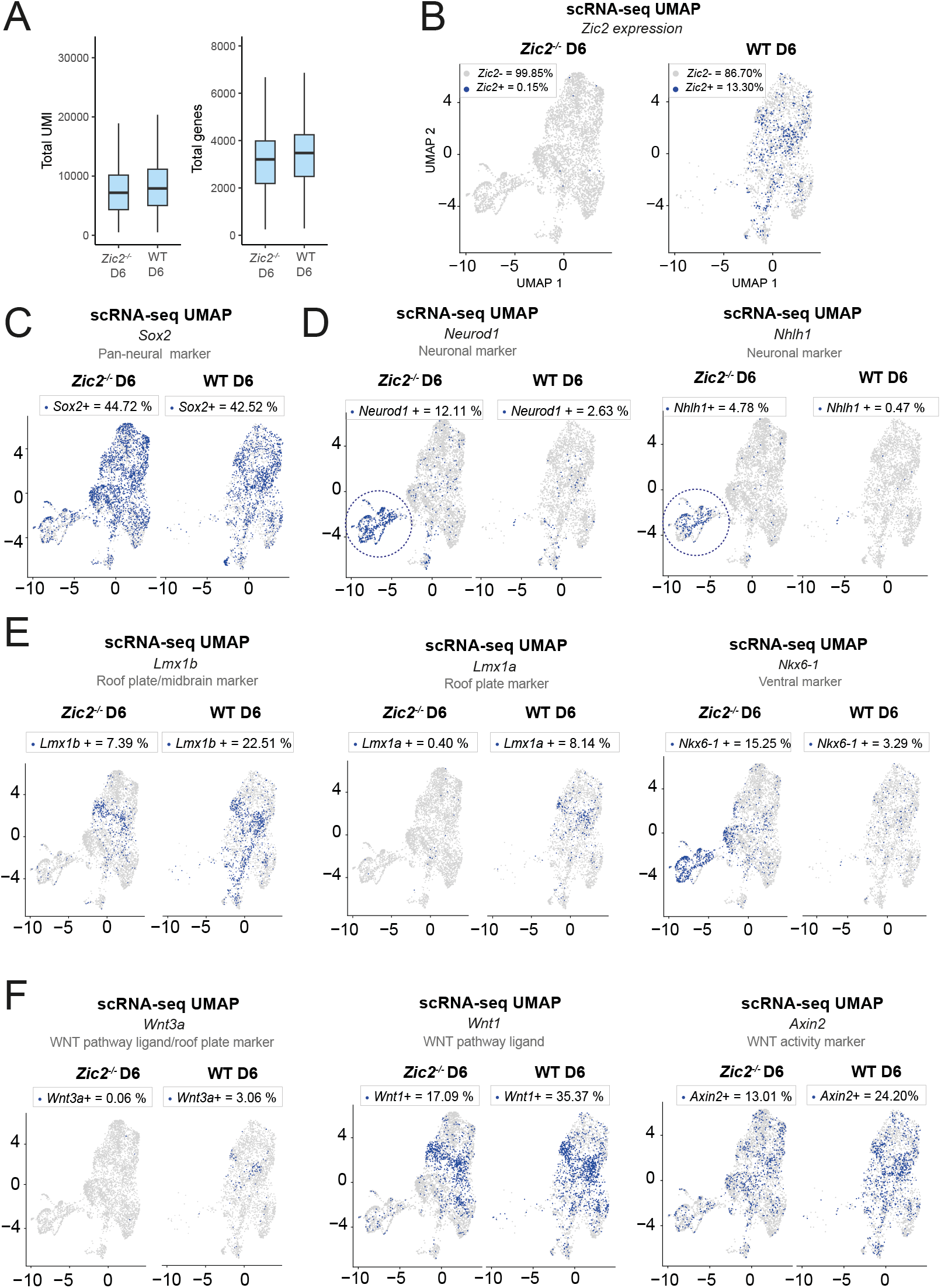
Single-cell RNA-seq analysis of WT and *Zic2*^*-/-*^ D6 AntNPCs. **(A)** Boxplots showing the distribution of total UMI counts (left) and number of detected genes (right) per cell in WT and *Zic2*^*-/-*^ D6 samples after quality filtering. **(B)** UMAP plots of the scRNA-seq data generated in *Zic2*^*-/-*^ and WT D6 AntNPC split by genotype. Cells expressing *Zic2* (≥ 1 read count) are labelled in blue. The percentage of *Zic2*-positive (*Zic2+*, in blue) and *Zic2*-negative (*Zic2*-, in grey) cells detected for each genotype are also shown. **(C-D)** UMAP plots showing the expression levels in WT and *Zic2*^*-/-*^ D6 AntNPC of the following genes: (C) the pan-neural marker *Sox2*, (D) post-mitotic neuronal markers (i.e. *Neurod1* and *Nhlh1*), (E) neural patterning genes (i.e. *Lmx1b, Lmx1a and Nkx6-1*) and (F) Wnt signalling ligands (i.e. *Wnt3a* and *Wnt1*) and *Axin2*. From **C** to **F**, the cells with detectable expression (*≥* 1 read count) of the corresponding genes are labelled in blue. The percentage of WT and *Zic2*^*-/-*^ D6 cells positive for each gene is also shown.

**Fig. S15.**
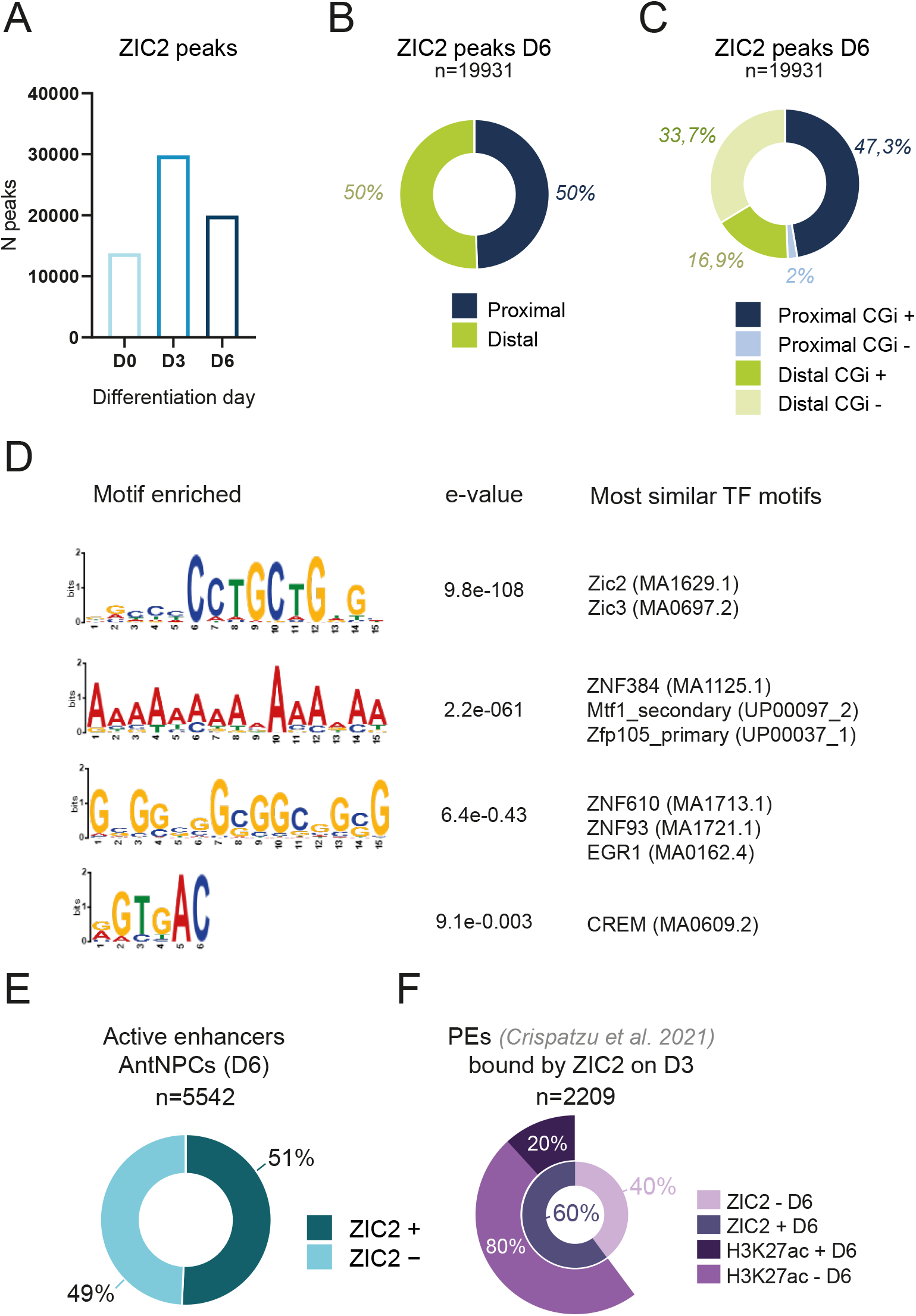
Genomic and epigenomic features of D6 ZIC2 peaks. **(A)** Bar plot indicating the total number (N) of ZIC2 ChIP-seq peaks identified in D0, D3 and D6 WT cells. **(B-C)** Classification of the D6 ZIC2 peaks according to their distance to (B) TSS and (C) CGI as described in Figure 1 D. **(D)**Top four most enriched motifs identified by MEME-ChIP when analysing D6 ZIC2 ChIP-Seq peaks.**(E)** Pie chart representing the percentage of active enhancers in D6 AntNPC that are bound by ZIC2 in D6 AntNPC. Active enhancers (ATAC-seq peak enriched in H3K27ac and H3K4me1) were called using the ChIP-Seq and ATAC-Seq datasets generated in WT D6 AntNPC (see Methods). **(F)** Pie chart representing the percentage of poised enhancers (39) bound by ZIC2 on D3 (n=2209) that remain bound by ZIC2 on D6 (60%) and further classified according to whether they become active (H3K27ac +) or stay inactive (H3K27ac -) by D6.

**Fig. S16.**
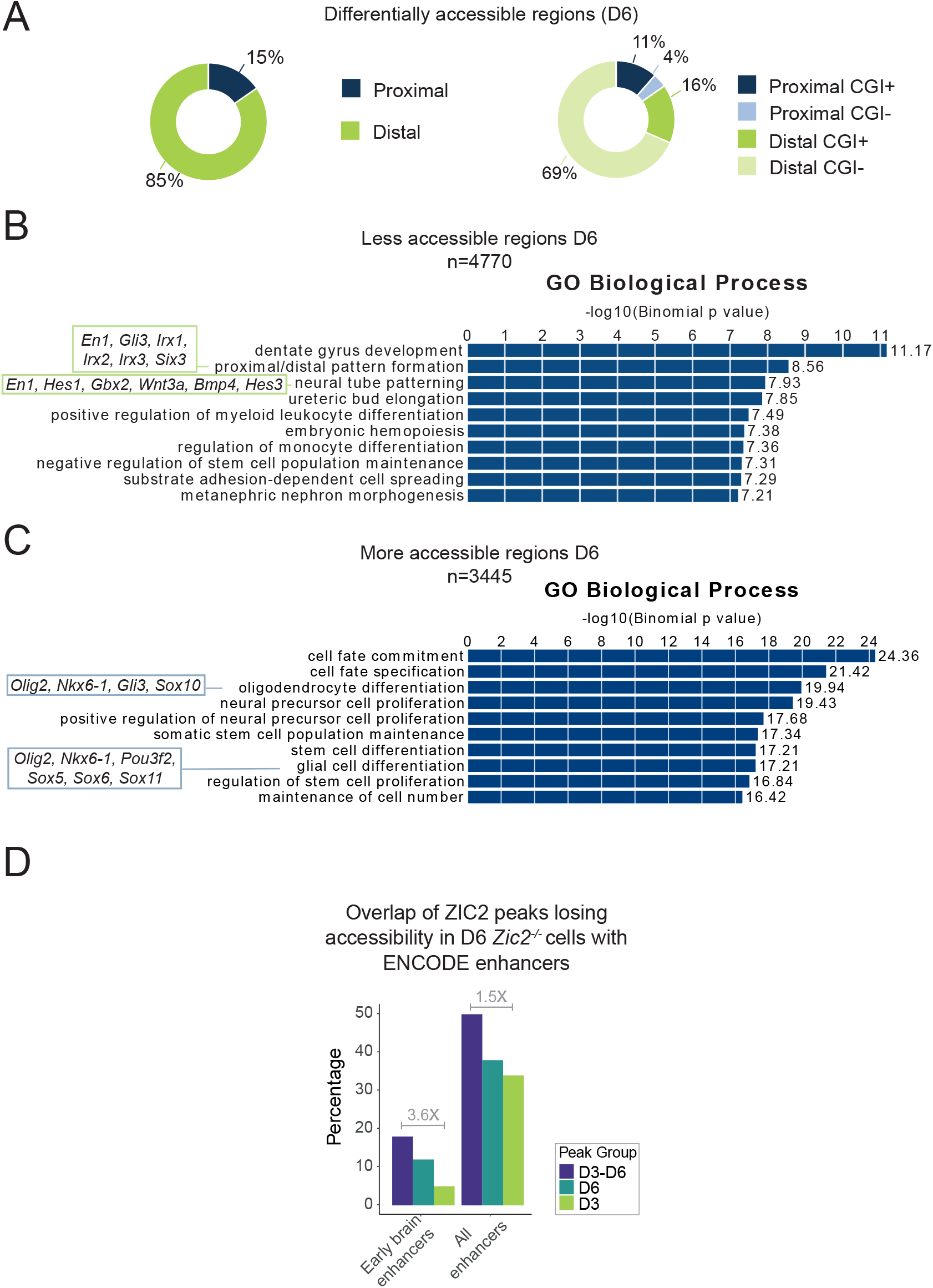
Genomic and transcriptional features associated with the regions displaying chromatin accessibility changes in *Zic2*^*-/-*^ AntNPCs D6. **(A)** Pie charts showing the classification of the differentially accessible regions identified in D6 AntNPC according to the ATAC-seq data generated in Zic2^*-/-*^ and WT D6 AntNPC. The differentially accessible regions are classified based on distance to TSS (left) and CGI (right) as described in Figure 1 D. **(B-C)** Functional enrichment analysis (performed using GREAT (146)) results showing the main GO Biological processes enriched in regions that are either less **(B)** or more **(C)** accessible in *Zic2*^*-/-*^ D6 AntNPC in comparison to WT D6 AntNPC. Green and blue boxes highlight some specific GO terms and include some of the responsible genes for the enrichment of those terms. **(D)** Bar plot showing the percentage of ZIC2-bound regions losing chromatin accessibility in *Zic2*^*-/-*^ cells that overlap with active enhancers annotated by the ENCODE consortium according to two broad categories i.e. early brain enhancers (forebrain and midbrain, E10.5–E11.5) and active enhancers across all embryonic tissues (see Methods). Three different groups of ZIC2 peaks were considered: ZIC2 peaks present only in D3 cells (D3), ZIC2 peaks present in both D3 and D6 cells (D3–D6), and ZIC2 peaks present only in D6 cells (D6). The fold-change differences in the overlap between the D3-6 and D3 ZIC2 peaks with respect to each ENCODE enhancer category (see Methods) are also indicated.

**Fig. S17.**
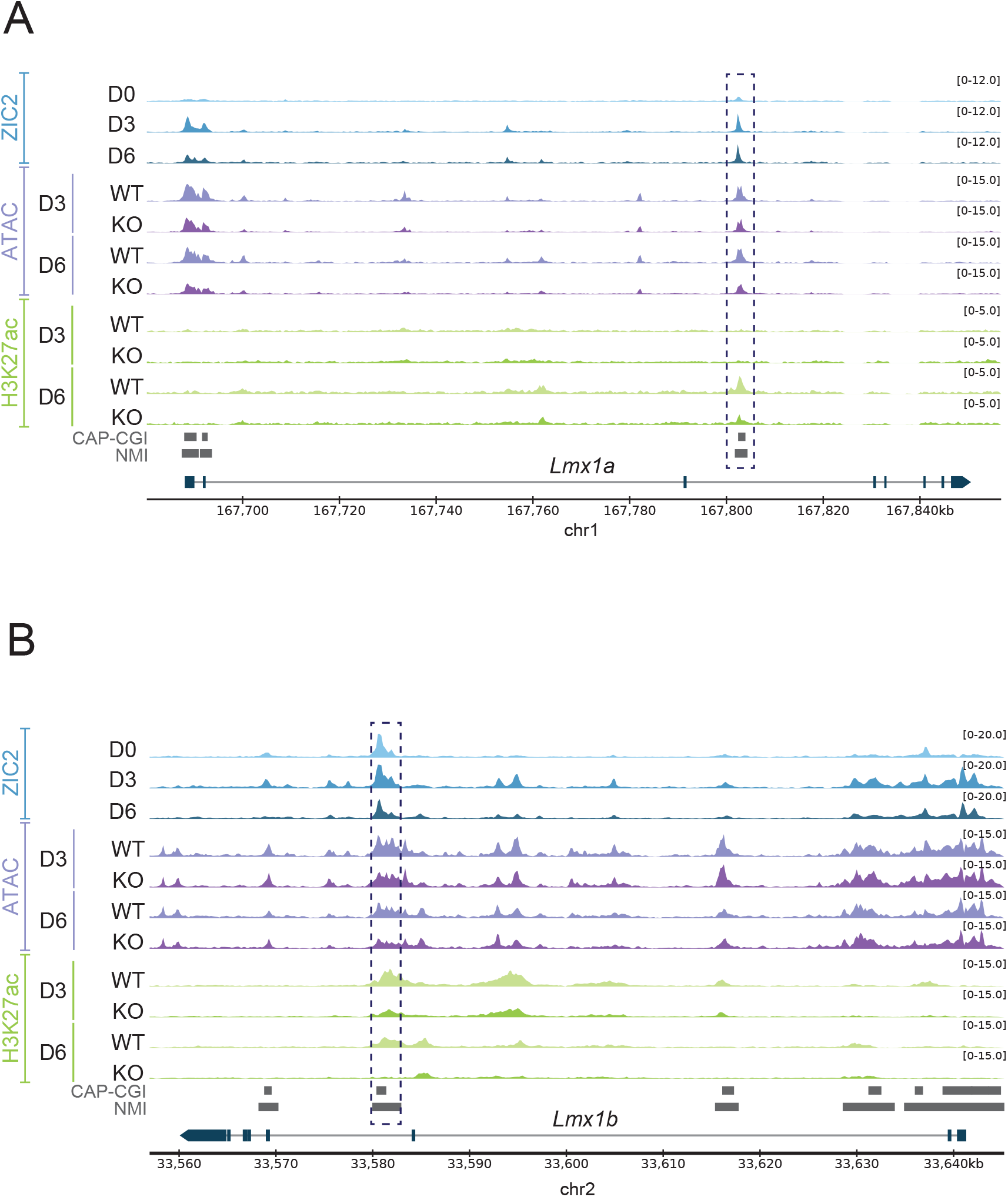
Chromatin dynamics of the ZIC2-bound enhancers located near *Lmx1a* and *Lmx1b* during neural differentiation. **(A–B)** Genome browser tracks showing the epigenetic landscape of the *Lmx1a* **(A)** and *Lmx1b* **(B)** loci during the differentiation of WT and *Zic2*^*-/-*^ (KO) mESC towards AntNPC. Tracks display ZIC2 ChIP-seq (blue), chromatin accessibility (ATAC-seq, purple) and H3K27ac ChIP-seq (green) profiles at the indicated differentiation time points (D0, D3 or D6). Two putative ZIC2-bound enhancers are highlighted with dashed boxes. CAP-CGI and NMI are shown as dark gray rectangles.

**Fig. S18.**
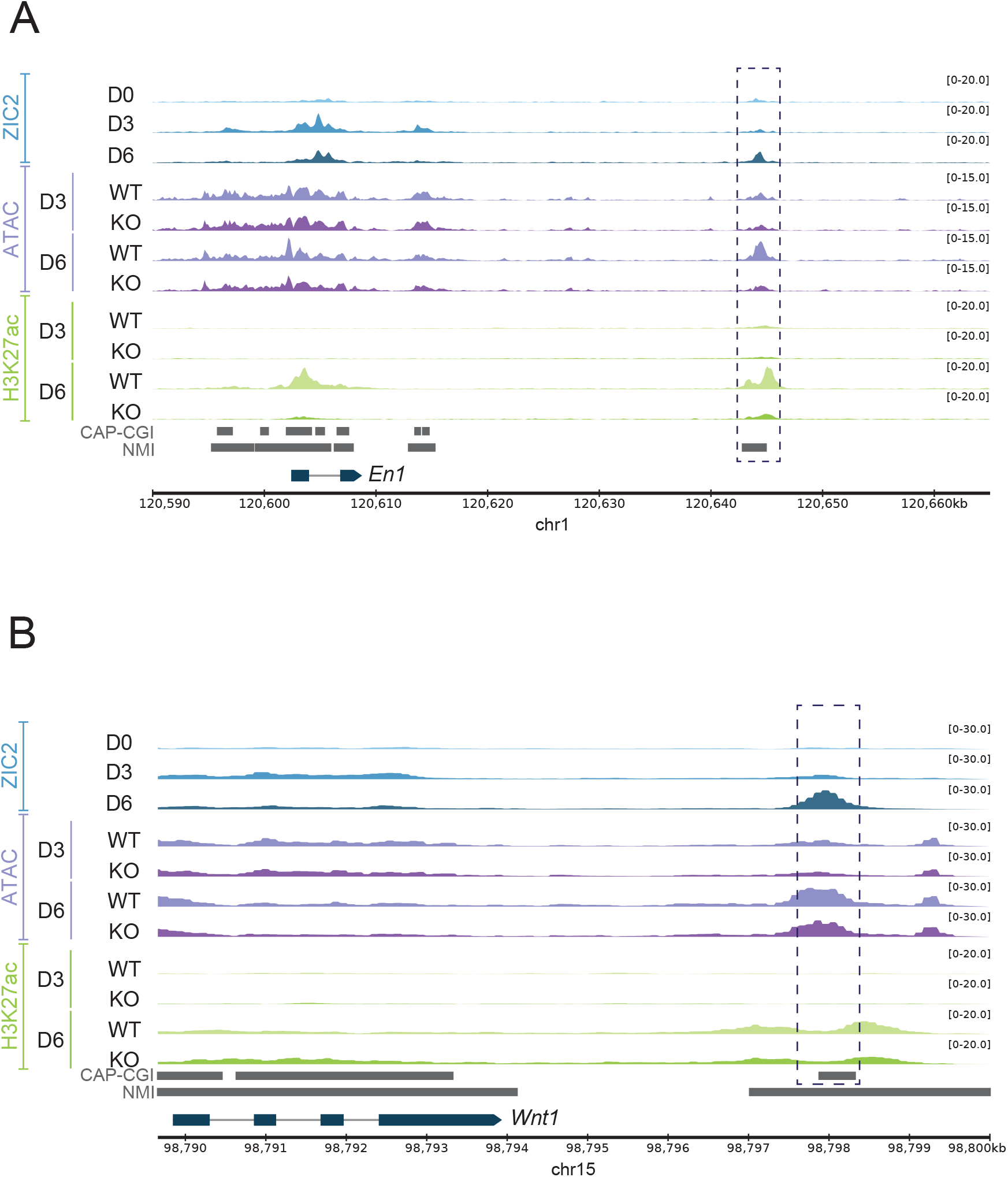
Chromatin dynamics of the ZIC2-bound enhancers located near *En1* and *Wnt1* during neural differentiation. **(A–B)** Genome browser tracks showing the epigenetic landscape of the *En1* **(A)** and *Wnt1* **(B)** loci during the differentiation of WT and *Zic2*^*-/-*^ (KO) mESC towards AntNPC. Tracks display ZIC2 ChIP-seq (blue), chromatin accessibility (ATAC-seq, purple) and H3K27ac ChIP-seq (green) profiles at the indicated differentiation time points (D0, D3 or D6). Two putative ZIC2-bound enhancers are highlighted with dashed boxes. CAP-CGI and NMI are shown as dark gray rectangles.

**Fig. S19.**
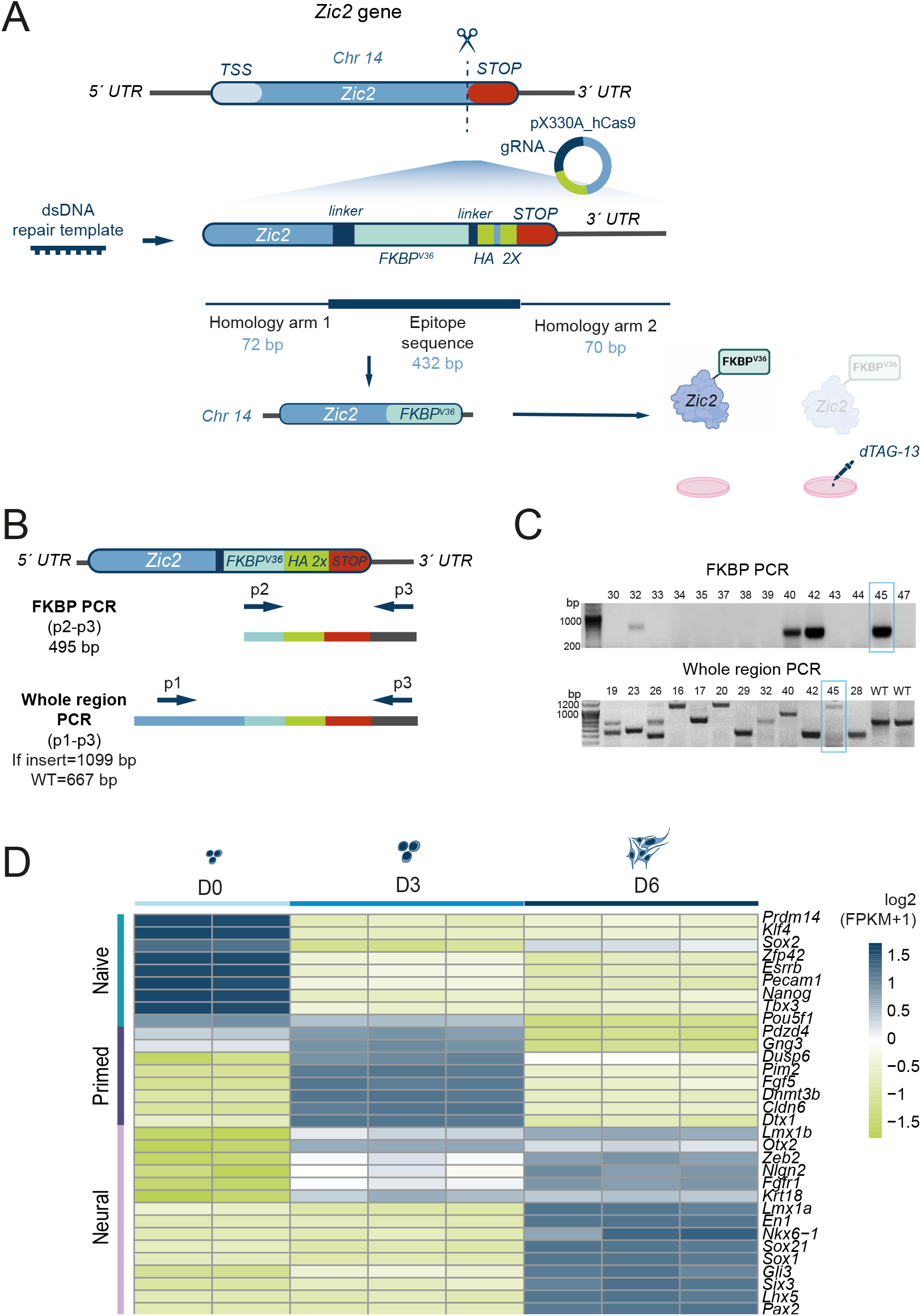
Generation and transcriptional characterization of the 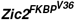 mESC line. **(A)** Graphical overview of the CRISPR-Cas9 genetic editing strategy employed to introduce the FKBP^V36^ epitope in the *Zic2* C-terminus region right before the stop codon. The scissors and the dashed line represent the cut site of the gRNA. The size of the different sequences included in the double stranded DNA (dsDNA) repair template used for epitope insertion are also shown. Upon insertion of the FKBP^V36^ epitope, ZIC2 is expressed as a fusion protein (ZIC2-FKBP^V36^) that gets degraded upon addition of d-TAG13 to the culture media. **(B)** PCR genotyping strategy to detect mESC clones positive for the insertion of the FKBP^V36^ epitope sequence. Primers used for genotyping are represented as arrows named as p1, p2 and p3. The PCR reactions employed (FKBP PCR and Whole region PCR) for genotyping and the expected sizes of the resulting PCR products are also depicted. **(C)** Agarose gel images showing the PCR genotyping results for several of the screened mESC clones. Clone #45 was selected as a representative 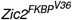 mESC line for subsequent ZIC2 depletion experiments.**(D)** Heat map showing the temporal expression patterns (as measured by RNA-Seq) of naive pluripotency, primed pluripotency and neural markers during differentiation of untreated 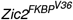 mESC into AntNPCs.

**Fig. S20.**
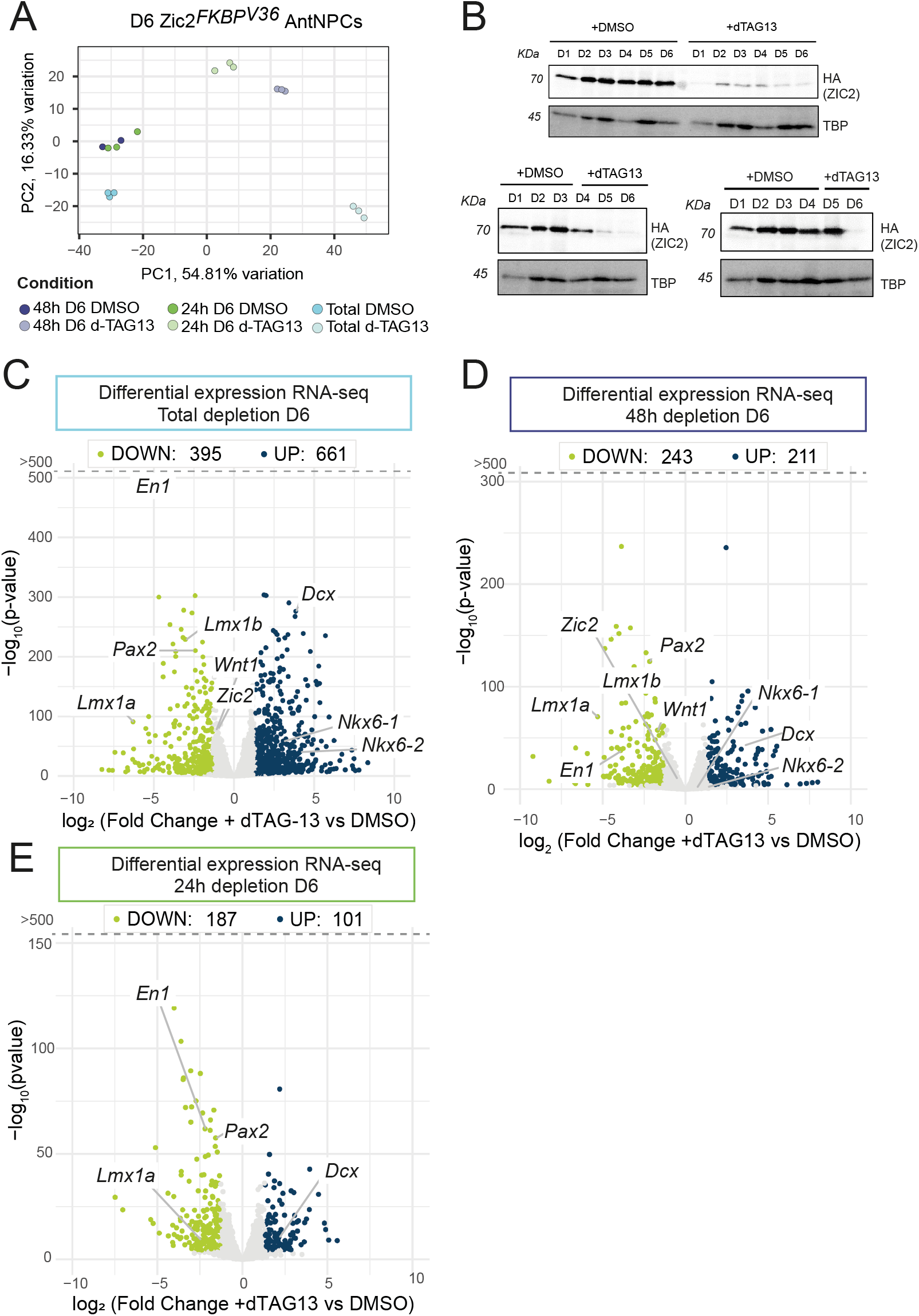
Transcriptional changes caused by ZIC2 depletion during AntNPC differentiation. **(A)** Principal Component Analysis (PCA) considering all the D6 RNA-Seq depletion samples described in Fig 7.A. **(B)** Western Blot analysis of ZIC2 protein levels in the ZIC2 depletion experiments described in Fig.7A. ZIC2 was detected using an anti-HA antibody. TBP is shown as a loading control. **(C-E)** Volcano plots showing the upregulated (blue) and downregulated (green) genes in dTAG13-treated vs DMSO-treated 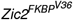D6 AntNPCs in the (C) “Total depletion”, (D) 48h and (E) 24h depletion experiments described in Fig. 4A.

**Fig. S21.**
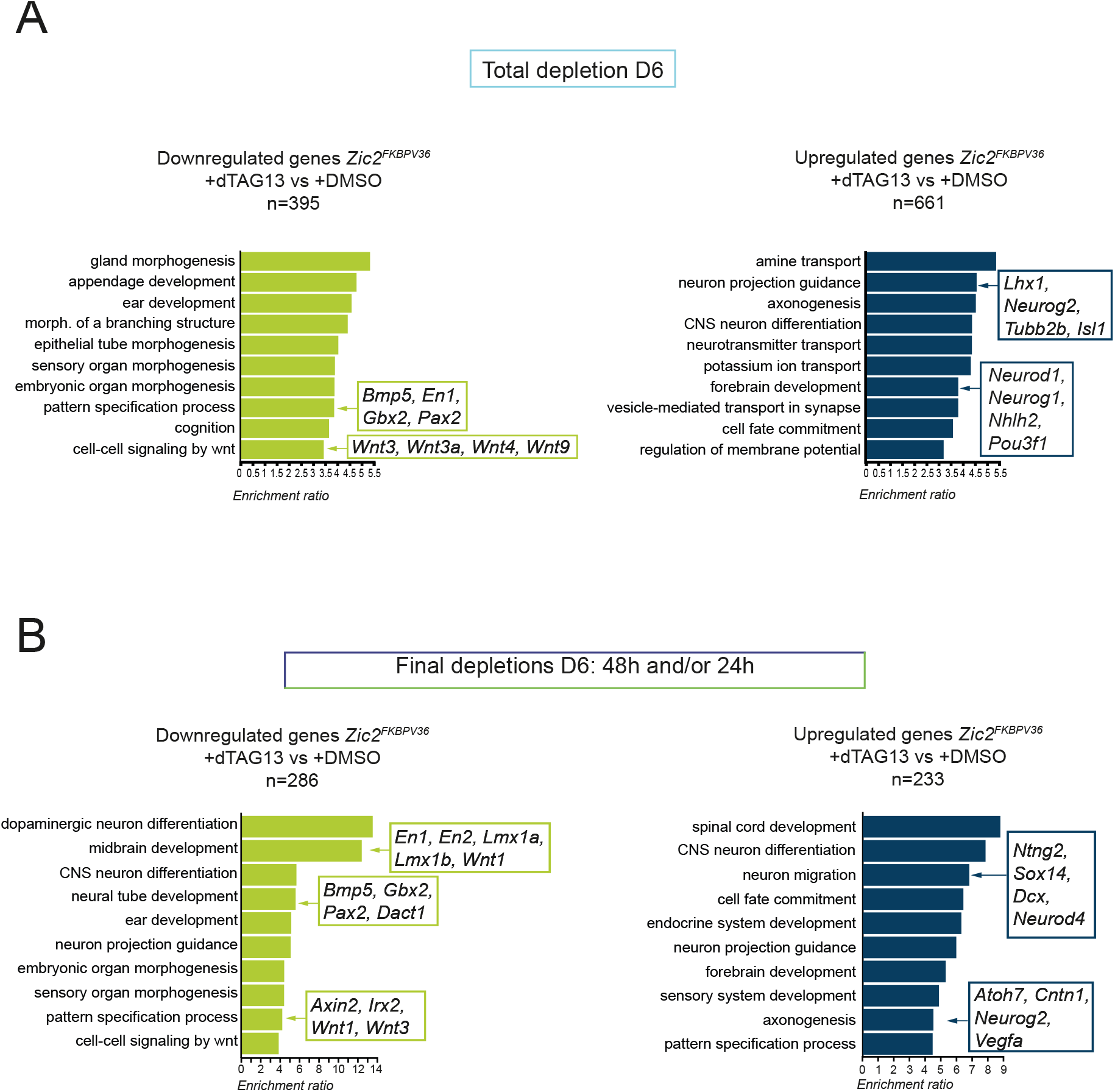
GO term enrichment of differentially expressed genes after ZIC2 targeted degradation. **(A-B)** Functional enrichment analysis (WebGestalt (65), GO terms for Biological Process non-redundant) of genes that were either upregulated (right panels; blue) or downregulated (left panels; green) in dTAG13-treated vs DMSO-treated 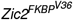 D6 AntNPCs in the (A) “Total depletion” or (B) 48h and/or 24h depletion experiments described in Fig. 4A. Genes associated with selected GO terms are indicated in boxes.

**Fig. S22.**
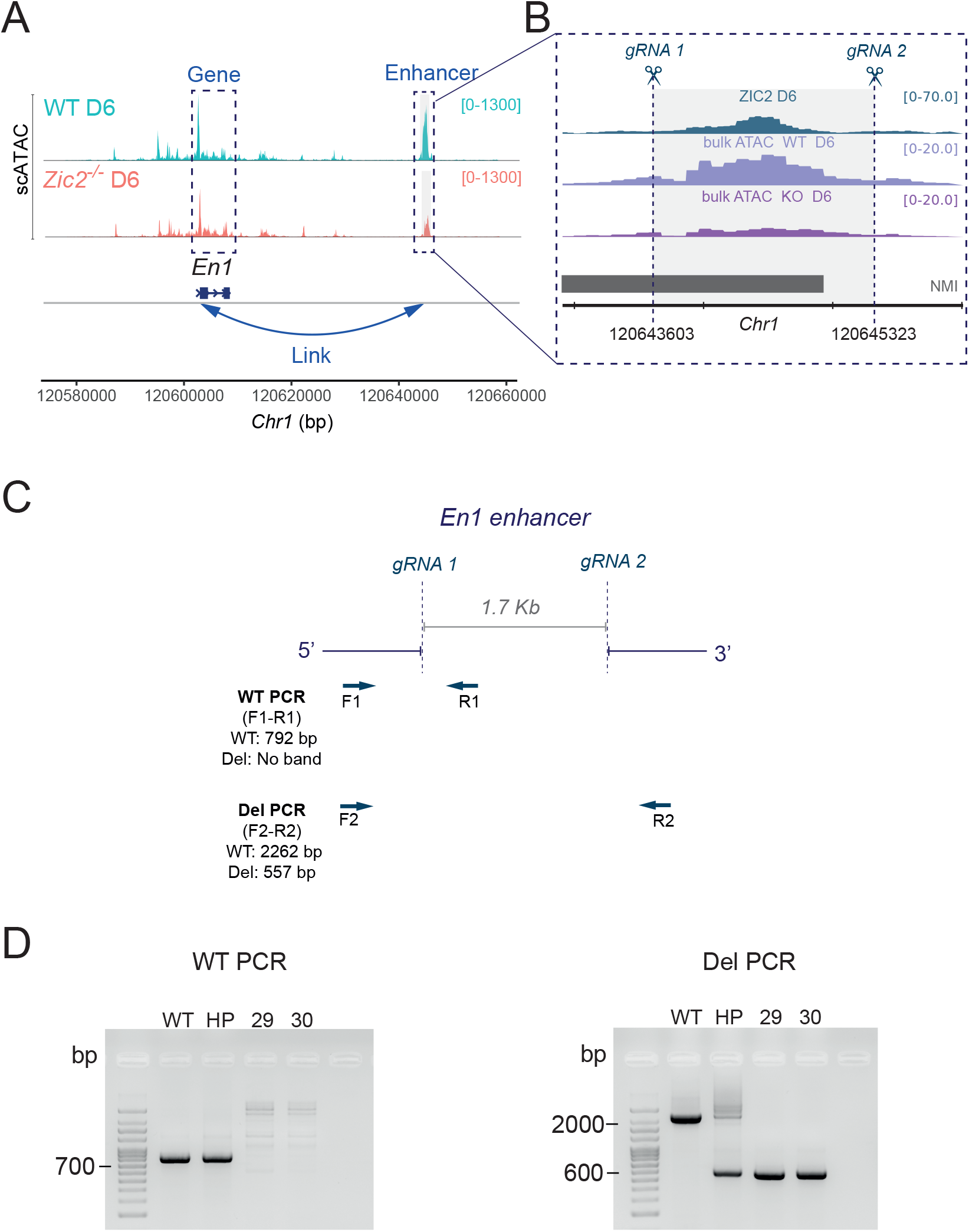
Functional validation of the ZIC2-bound enhancer linked to *En1*. **(A)** Genome browser snapshot showing chromatin accessibility (scATAC-seq) at the *En1* locus in WT (coral) and *Zic2*^-/-^ (turquoise) D6 AntNPCs. The predicted ZIC2-regulated enhancer is highlighted and linked to *En1* based on the Multiome analysis described in Fig. 7D and in Methods. **(B)** Close-up view of the *En1* enhancer highlighted in (A), showing ZIC2 ChIP-seq profiles in WT D6 AntNPC as well as bulk ATAC-seq profiles in WT and *Zic2*^*-/-*^ (KO) D6 AntNPC. Positions of the CRISPR gRNAs used to deleted the *En1* enhancer are indicated with scissors and dot lines. **(C)** Genotyping strategy to identify mESC lines with deletions spanning the *En1* enhancer. Arrows indicate the positions of the different genotyping primers (F1, R1, F2 and R2). The size of the expected amplicons for WT-specific and deletion-specific PCRs are shown. **(D)** Agarose gel images showing the PCR genotyping results obtained using genomic DNA from WT mESC, an heterogeneous population of mESC transfected with the gRNAs depicted in (B-C) (HP), and two clonal mESC lines with homozygous deletions of the *En1* enhancer(clones #29 and #30). WT-specific (WT PCR; top) and deletion-specific (Del PCR; bottom) PCR results are shown with the expected product sizes.

**Fig. S23.**
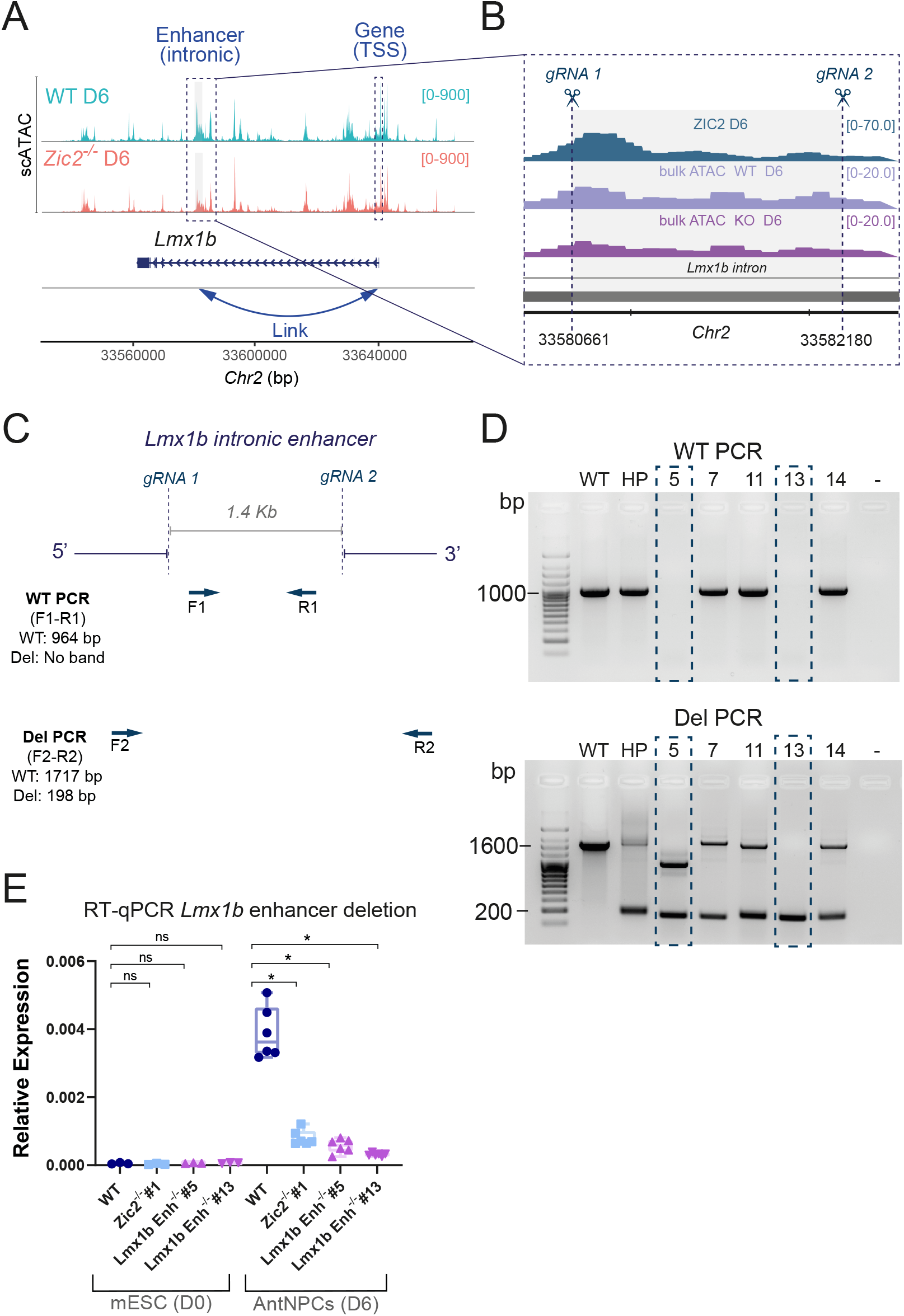
Functional validation of the ZIC2-bound enhancer linked to *Lmx1b*. **(A)** Genome browser snapshot showing chromatin accessibility (scATAC-seq) at the *Lmx1b* locus in WT (coral) and *Zic2*^-/-^ (turquoise) D6 AntNPCs. The predicted ZIC2-regulated enhancer is highlighted and linked to *Lmx1b* based on the Multiome analysis described in Fig. 7D. **(B)** Close-up view of the *Lmx1b* enhancer highlighted in (A), showing ZIC2 ChIP-seq profiles in WT D6 AntNPC as well as bulk ATAC-seq profiles in WT and *Zic2*^*-/-*^ (KO) D6 AntNPC. Positions of the CRISPR gRNAs used to deleted the *Lmx1b* enhancer are indicated with scissors and dot lines. **(C)** Genotyping strategy to identify mESC lines with deletions spanning the *Lmx1b* enhancer. Arrows indicate the positions of the different genotyping primers (F1, R1, F2 and R2). The sized of the expected amplicons for WT-specific and deletion-specific PCRs are shown. **(D)** Agarose gel images showing the PCR genotyping results obtained using genomic DNA from WT mESC, an heterogeneous population of mESC transfected with the gRNAs depicted in (B-C) (HP), and two clonal mESC lines with homozygous deletions of the *Lmx1b* enhancer (clones #5 and #13). WT-specific (WT PCR; top) and deletion-specific (Del PCR; bottom) PCR results are shown with the expected product sizes.**(E)** *Lmx1b* expression levels were measured by RT-qPCR in mESC and D6 AntNPC that were either WT, *Zic2*^*-/-*^ (clone #1) or homozygous for the *Lmx1b* enhancer deletion depicted in (C) (clones #5 and #13). For each cell line, *Lmx1b* expression was measured in the following number of replicates: D0 samples - one biological replicate with three technical replicates; D6 samples - two biological replicates with three technical replicates each . Expression values were normalized to two housekeeping genes (*Eef1a1* and *Hprt1*). The statistical significance of the expression differences among cell lines was calculated using two-sided unpaired t-tests with Welch’s correction (* fold-change > 2 and P < 0.0001; NS (not significant): fold-change < 2 or P > 0.05). Error bars represent the standard deviation.

**Fig. S24.**
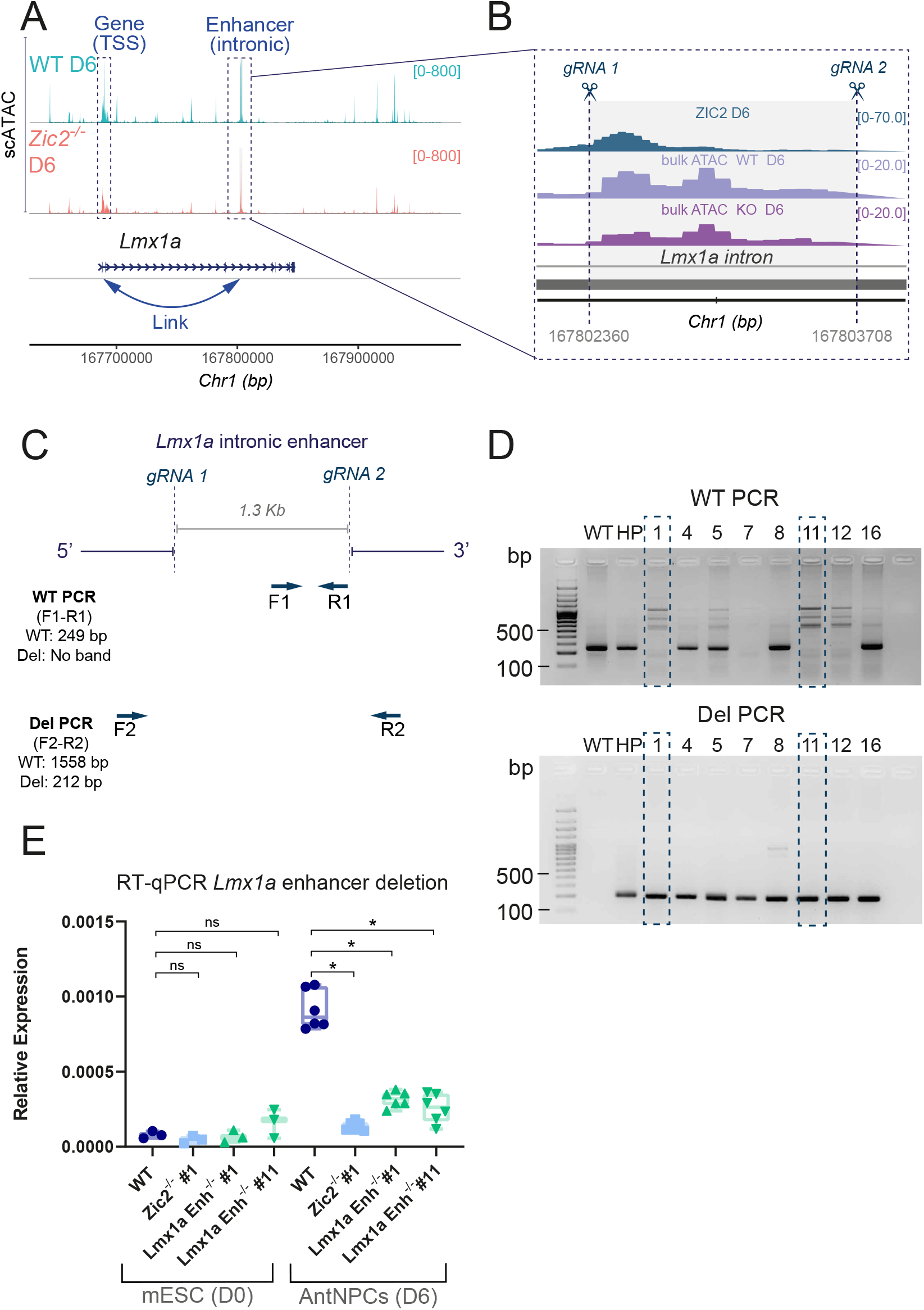
Functional validation of a ZIC2-bound enhancer linked to *Lmx1a*. **(A)** Genome browser snapshot showing chromatin accessibility (scATAC-seq) at the *Lmx1a* locus in WT (coral) and *Zic2*^-/-^ (turquoise) D6 AntNPCs. The predicted ZIC2-regulated enhancer is highlighted and linked to *Lmx1a* based on the Multiome analysis described in Fig. 7D. **(B)** Close-up view of the *Lmx1a* enhancer highlighted in (A), showing ZIC2 ChIP-seq profiles in WT D6 AntNPC as well as bulk ATAC-seq profiles in WT and *Zic2*^*-/-*^ (KO) D6 AntNPC. Positions of the CRISPR gRNAs used to deleted the *Lmx1a* enhancer are indicated with scissors and dot lines. **(C)** Genotyping strategy to identify mESC lines with deletions spanning the *Lmx1a* enhancer. Arrows indicate the positions of the different genotyping primers (F1, R1, F2 and R2). The sized of the expected amplicons for WT-specific and deletion-specific PCRs are shown. **(D)** Agarose gel images showing the PCR genotyping results obtained using genomic DNA from WT mESC, an heterogeneous population of mESC transfected with the gRNAs depicted in (B-C) (HP), and two clonal mESC lines with homozygous deletions of the *Lmx1a* enhancer (clones #1 and #11). WT-specific (WT PCR; top) and deletion-specific (Del PCR; bottom) PCR results are shown with the expected product sizes. **(E)** *Lmx1b* expression levels were measured by RT-qPCR in mESC and D6 AntNPC that were either WT, *Zic2*^*-/-*^ (clone #1) or homozygous for the *Lmx1a* enhancer deletion depicted in (C) (clones #1 and #11). For each cell line, *Lmx1a* expression was measured in the following number of replicates: D0 samples - one biological replicate with three technical replicates; D6 samples - two biological replicates with three technical replicates each . Expression values were normalized to two housekeeping genes (*Eef1a1* and *Hprt1*). The statistical significance of the expression differences among cell lines was calculated using two-sided unpaired t-tests with Welch’s correction (* fold-change > 2 and P < 0.0001; NS (not significant): fold-change < 2 or P > 0.05). Error bars represent the standard deviation.

**Fig. S25.**
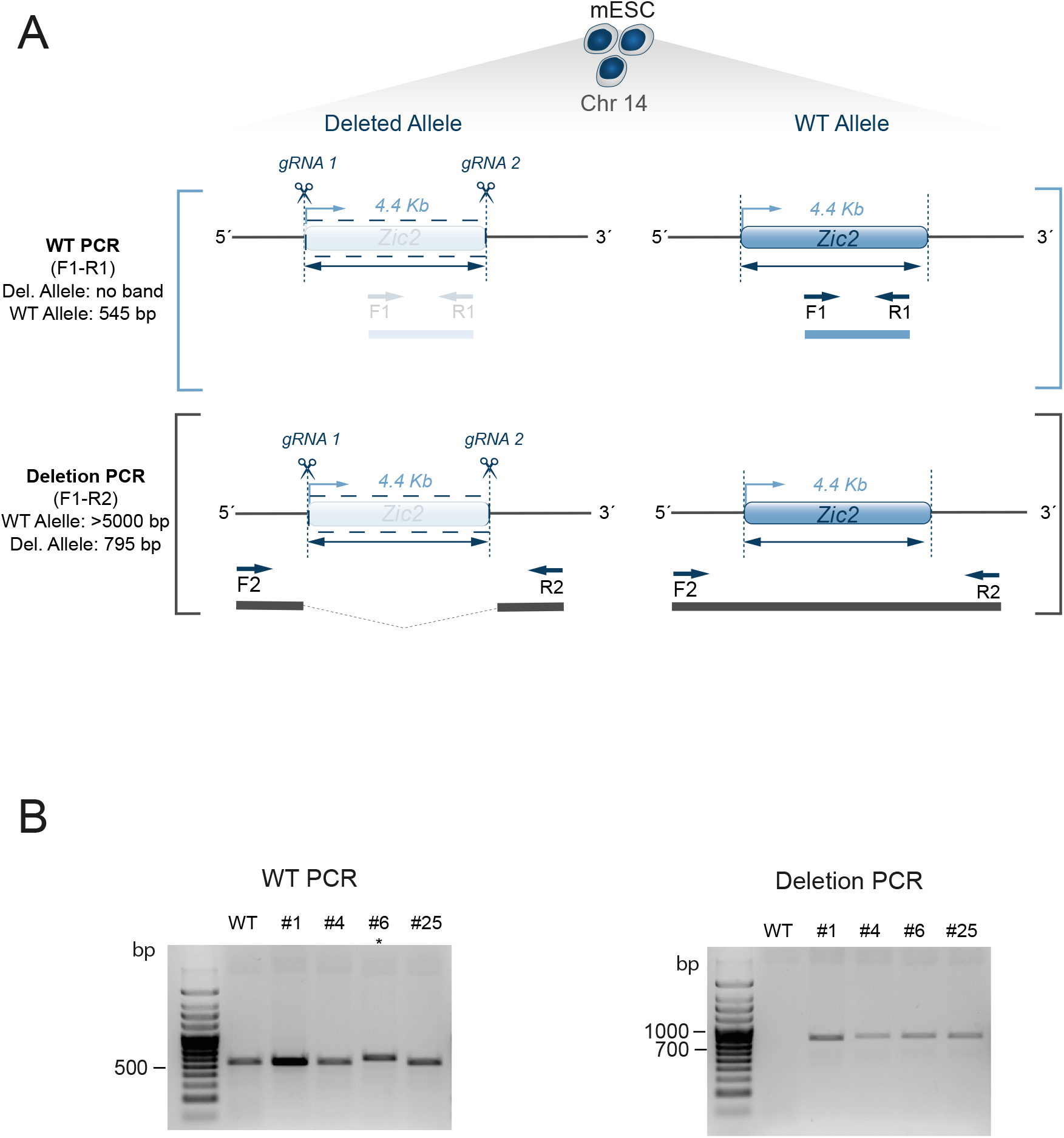
Generation of *Zic2*^*+/-*^ mESCs lines. **(A)** PCR genotyping strategy to identify *Zic2*^*+/-*^ mESC lines. Scissors represent the cut sites of the upstream 5’and downstream 3’ guide RNAs (gRNAs). Primers used for genotyping are represented as arrows named F1, R1, F2 and R2. The slightly higher amplicon size of clone #6 (*) is due to a FLAG-HA inserted at its C-terminus, for unrelated purposes. **(B)** Agarose gel images showing the genotyping results (WT and Deletion PCR) of four different *Zic2*^*+/-*^ heterozygous mESC clones (clones #1, #4, #6 and #25).

